# Massively parallel characterization and deep learning of enhancers in plant genomes

**DOI:** 10.64898/2026.04.26.720828

**Authors:** Tobias Jores, Nicholas A. Mueth, Sayeh Gorjifard, Sebastian Triesch, Dominic Schirmer, Jackson Tonnies, Kerry L. Bubb, Josh T Cuperus, Stanley Fields, Christine Queitsch

**Author notes:** Correspondence: Tobias Jores, Christine Queitsch.

## Abstract

Enhancers coordinate gene expression in response to developmental and environmental cues. Because plant enhancers lack the readily detectable molecular hallmarks of animal enhancers, their systematic functional characterization has yet to be accomplished. Here, we characterize the species- and condition-specific enhancer activity of over 350,000 sequences derived from accessible chromatin regions of the *Arabidopsis*, tomato, maize, and sorghum genomes. Enabled by the massive scale of the data, we developed plantGREP, a deep learning model that predicts enhancer strength and identifies the underlying functional sequence motifs. We apply plantGREP to evolve strong constitutive as well as species- and condition-specific enhancers, and to locate regions with enhancer activity upstream of developmental genes in crop genomes. These results should facilitate the targeted editing of enhancers in crop genomes and the design of cell-type-specific plant enhancers.

## Main text

Enhancers bind transcription factors and interact with the core transcription machinery to drive transcription^1–4^. They function independently of orientation, can reside upstream or downstream of the target core promoter, and are active over a wide range of distances^5–7^. By recruiting transcription factors specific to particular conditions or cell types, enhancers integrate environmental and developmental cues to control gene expression^1,2^. In animals, enhancers can facilitate expression in specific cell types while repressing it in others, reflecting cell-type-specific transcription factor repertoires^8^.

These fundamental features of enhancers are thought to be shared between animals and plants. However, plant enhancers lack the canonical flanking histone modifications and the highly expressed, short-lived enhancer RNAs that mark active animal enhancers^9–15^, which has hampered their large-scale identification in plant genomes^2,16^.

With the advent of efficient genome engineering, enhancers have emerged as promising targets for crop improvement. Introducing deletions to disrupt regions upstream of key developmental genes has improved traits in tomato, rice, and maize^17–21^. However, these studies relied on phenotyping many different deletion lines to find genotypes with improved traits. This laborious process can be avoided when the target regulatory elements are known^20,22^. Thus, the ability to rapidly identify or predict crop enhancers will accelerate genome engineering and crop improvement.

To acquire this ability, we characterized the species- and condition-specific enhancer activity of over 350,000 genomic sequences derived from the accessible chromatin regions of four diverse plant genomes. Leveraging the scale of these data, we built a deep learning model capable of predicting the enhancer strength of novel sequences. We demonstrate that this model can generate enhancers that rival in strength the commonly used viral 35S enhancer while also providing strict species and condition specificity. The model can localize enhancer regions in the genomes of common crop plants and identify the underlying sequence features contributing to enhancer activity.

## Results

### Plant enhancers show species and condition specificity and orientation independence

We focused on accessible chromatin regions (ACRs) enriched for active enhancers^1,2,4,11^ and selected 169,830 sequences derived from ACRs in the genomes of *Arabidopsis* (*Arabidopsis thaliana*), tomato (*Solanum lycopersicum*), maize (*Zea mays*), and sorghum (*Sorghum bicolor*), and 6,464 sequences from inaccessible regions in these genomes (Supplementary Table 1)^17,23–25^. We included 15,670 sequences tiled across the upstream regions of 147 tomato genes (Supplementary Table 2) and 36 sequences derived from regulatory elements^26,27^. These sequences were array-synthesized as 170-base oligonucleotides and cloned upstream of a 35S minimal promoter to drive transcription of a barcoded reporter gene (Fig. 1a). The sequences were cloned in either the forward or reverse orientation, resulting in a final “ACR library” of 384,000 unique sequences. Each sequence was linked to multiple unique barcodes by next-generation sequencing. The ACR library was subjected to the massively parallel reporter assay Plant STARR-seq^28^. After transformation, the reporter mRNA was extracted from the plant tissue, and the relative abundance of the barcode sequences in the input DNA and output RNA was determined by sequencing. We define the enrichment of a barcode in the RNA relative to its DNA input as the measure of the enhancer strength of the corresponding tested sequence, depicted relative to core promoter strength (a control construct without an enhancer that is scored as log_2_ = 0)^28^. As the library contains sequences that show little or repressive activity, enhancer strength as defined here ranges from negative to positive values.

**Fig. 1.**
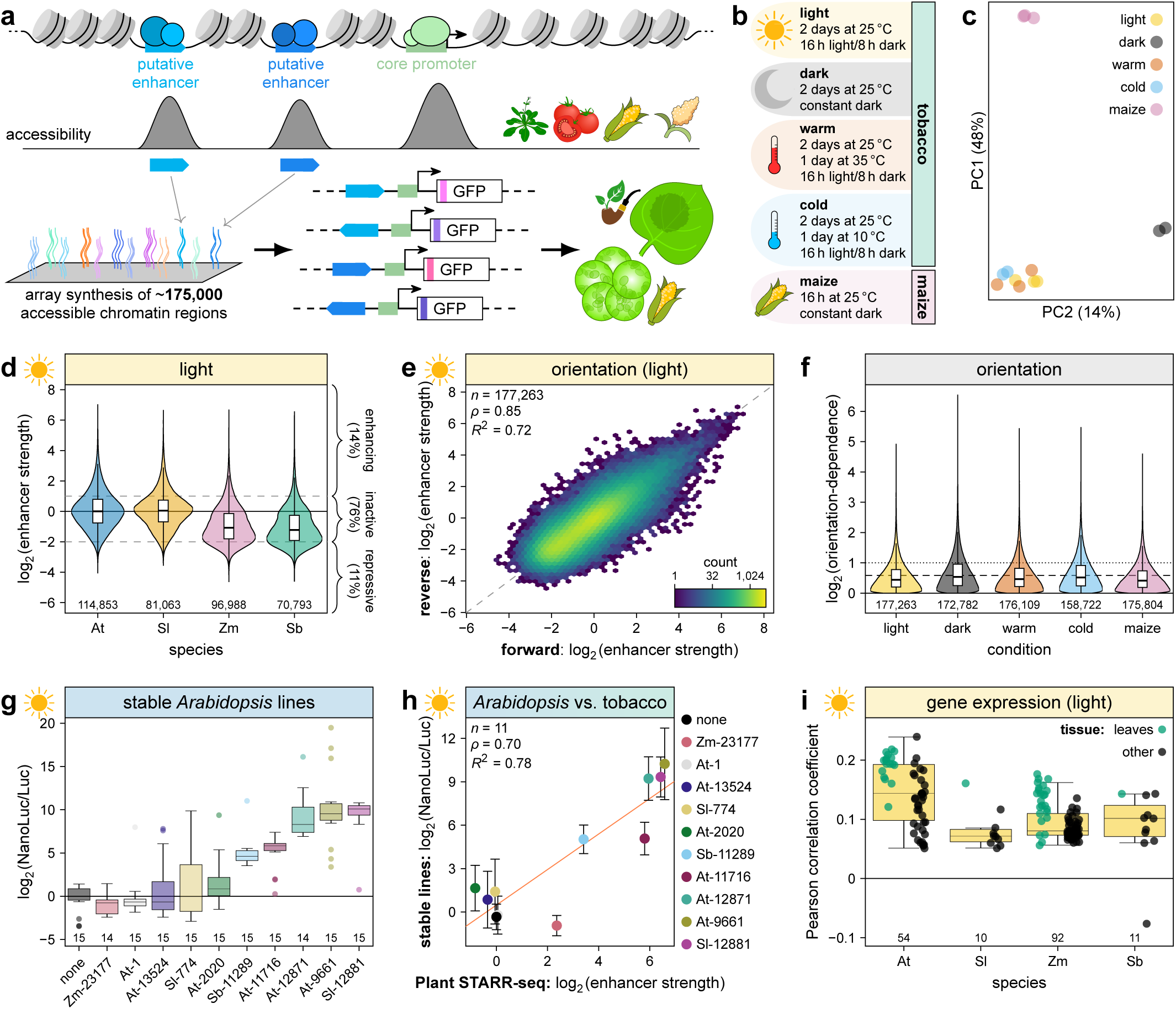
| Characterization of enhancer strength across species and conditions. **a**,**b**, Test sequences (170 bp) were derived from accessible chromatin regions (ACRs) in the genomes of *Arabidopsis* (At), tomato (Sl), maize (Zm), and sorghum (Sb). The array-synthesized test sequences were cloned in the forward and reverse orientation upstream of a 35S minimal promoter driving the expression of a barcoded GFP reporter gene. The plasmid library was subjected to Plant STARR-seq in transiently transformed tobacco leaves and maize leaf protoplasts (**a**). After transformation, the plants or protoplasts were subjected to different light and temperature conditions before RNA extraction (**b**). **c**, Principal component analysis of enhancer strength determined in Plant STARR-seq replicate experiments. **d**, Enhancer strength of test sequences grouped by species of origin. The percentage of enhancing (log_2_ (enhancer strength) > 1), inactive (log_2_ (enhancer strength) between 1 and −2), and repressive (log_2_ (enhancer strength) < −2) sequences is indicated. **e**,**f**, Correlation of (**e**) and fold-change in (**f**) enhancer strength between sequences tested in the forward and reverse orientation across the indicated Plant STARR-seq conditions. The dashed and dotted lines in (**e**) represent a 1.5-fold and 2-fold difference, respectively. **g**,**h**, The enhancer strength of ten ACR sequences selected from the Plant STARR-seq library and a negative control without an enhancer (none) was determined using a dual-luciferase assay in stable *Arabidopsis* lines (**g**) and compared to the enhancer strength determined by Plant STARR-seq in tobacco leaves in the light (**h**). **i**, Correlation between enhancer strength and gene expression in various tissues of the indicated species. ACR sequences were linked to the closest transcription start site. For each gene, only the ACR sequence with the highest enhancer strength was retained. Violin plots in (**d**,**f**) represent the kernel density distribution. Box plots inside violin plots and in (**g**,**i**) represent the median (center line), upper and lower quartiles (box limits), 1.5× interquartile range (whiskers), and outliers (points) for all corresponding samples. Numbers at the bottom of each violin or box plot indicate the number of samples in each group. In the hexbin plot in (**d**), color indicates the number of observations in each hexagon. In (**h**), the solid line represents a linear regression line and error bars represent the 95% confidence interval. Pearson’s *R*^2^, Spearman’s *ρ*, and the number (*n*) of samples are indicated in (**e**,**h**).

To detect species-specific enhancer activity, we performed Plant STARR-seq in transiently transformed tobacco (*Nicotiana benthamiana*) leaves and maize protoplasts. In tobacco, we measured enhancer strength after exposing the transformed plants to four growth conditions (Fig. 1b): two days in normal light/dark cycles (light condition) or completely in the dark (dark condition); and two days at the normal growth temperature of 25°C and normal light/dark cycles, followed by one day at increased (35°C; warm condition) or decreased (10°C; cold condition) temperature.

Experimental replicates were highly correlated (Fig. 1c and Supplementary Fig. 1). Across all experiments, transcriptional activity ranged over 1000-fold between the strongest enhancers and the most repressive elements (Fig. 1d, Extended Data Fig. 1a and Supplementary Data 1). Overall, 33% of the sequences enhanced expression (log_2_(enhancer strength) > 1) and 27% decreased expression (log_2_(enhancer strength) < −2) in at least one condition or assay system (Fig. 1d and Supplementary Table 3). Nearly 43% of test sequences showed little activity (log_2_(enhancer strength) between −2 and 1) across all conditions and in both assay systems (Supplementary Table 3). Enhancer strength in tobacco and maize was uncorrelated (*R*^2^ between 0.01 and 0.08; Fig. 1b and Supplementary Fig. 2). Because light had a profound effect on enhancer activity there were only moderate correlations between results obtained from plants kept in the light (light, warm, and cold conditions) and dark (*R*^2^ between 0.47 and 0.56). Enhancer activity in warm and cold conditions was well correlated with that in control conditions (light condition; *R*^2^ of 0.83 and 0.85; Fig. 1b and Supplementary Fig. 2). The function of individual plant enhancers is largely orientation-independent^26,28–32^. By systematically testing all test sequences in the forward and reverse orientation, we find that orientation-independence is indeed a general property of plant enhancers (Figs. 1e, f and Extended Data Fig. 1b).

To ensure that the results obtained with Plant STARR-seq reflect enhancer activity in a genomic context, we measured the enhancer strength of ten ACR sequences in stable transgenic *Arabidopsis* lines using a dual-luciferase assay (Supplementary Fig. 4). The results of the two assays were well correlated (*R*^2^ = 0.78; Figs. 1g, h).

### Features associated with enhancer strength in Plant STARR-seq

ACR sequences showed significantly higher enhancer strength than randomly sampled sequences from inaccessible regions of the same genomes (Supplementary Fig. 4). Moreover, the level of accessibility of an ACR sequence in the genome was predictive of its enhancer strength in Plant STARR-seq (Extended Data Fig. 2a). Genomic regions with acetylated histones yielded ACR sequences with only slightly more enhancer strength than did regions with methylated histones (Extended Data Fig. 2b), consistent with the inability of histone marks to serve as reliable hallmarks of plant enhancer activity.

Enhancer strength was highest for intergenic sequences and lowest for sequences from coding regions (Extended Data Fig. 2c). Grouping ACR sequences by their position relative to the nearest transcription start site, sequences residing directly upstream of the transcription start site in the genomic context showed the highest enhancer strength, with strength diminishing as distance from the transcription start site increased (Extended Data Fig. 2d).

We found a positive correlation (Pearsons correlation coefficient of up to 0.24) between Plant STARR-seq-determined enhancer strength and gene expression data from several tissues of light-grown plants^33–37^, with the strongest correlations observed in leaf samples, the tissue used in the Plant STARR-seq experiments (Fig. 1i and Extended Data Fig. 3a). Moreover, the Plant STARR-seq-determined enhancer condition specificities resembled the condition specificities observed in gene expression data^38^ (Extended Data Fig. 3b, c).

Across conditions, the genes linked to the top 5% of sequences with the highest enhancer strength were enriched (adjusted *p* value ≤ 0.05) for stimulus-responsive Gene Ontology (GO) terms; GO terms associated with plastid proteins, especially proteins in thylakoids, were enriched in the light condition (Extended Data Fig. 3d and Supplementary Data 2).

GC content exerted a strong negative effect on enhancer strength in the tobacco system under all tested conditions, but showed the opposite trend in maize (Extended Data Fig. 4a). This difference likely reflects the more AT-rich nature of dicot genomes than of monocot genomes, a feature even more pronounced in their ACRs (Extended Data Fig. 4b,c).

Because enhancers recruit transcription factors, we sought to quantify the effect of transcription factor binding sites on enhancer strength. Plants have large repertoires of transcription factors, which often bind similar sequences^39–42^. Here, we used 72 consensus binding motifs (Supplementary Table 4)^39^ to identify putative transcription factor binding sites in the ACR sequences. We calculated the transcription factor effect by comparing the enhancer strength of ACR sequences with a predicted binding site to that of ACR sequences lacking it. While we identified both activating motifs, which increase enhancer strength up to 70%, and repressive motifs, which decrease it up to 20%, activating motifs were more prevalent and showed higher absolute effect sizes (Fig. 2a, Extended Data Fig. 5, Supplementary Fig. 5). The correlation of transcription factor effects between tobacco and maize was low, consistent with differences in their transcription factor repertoires (Fig. 2a and Supplementary Fig. 6).

**Fig. 2.**
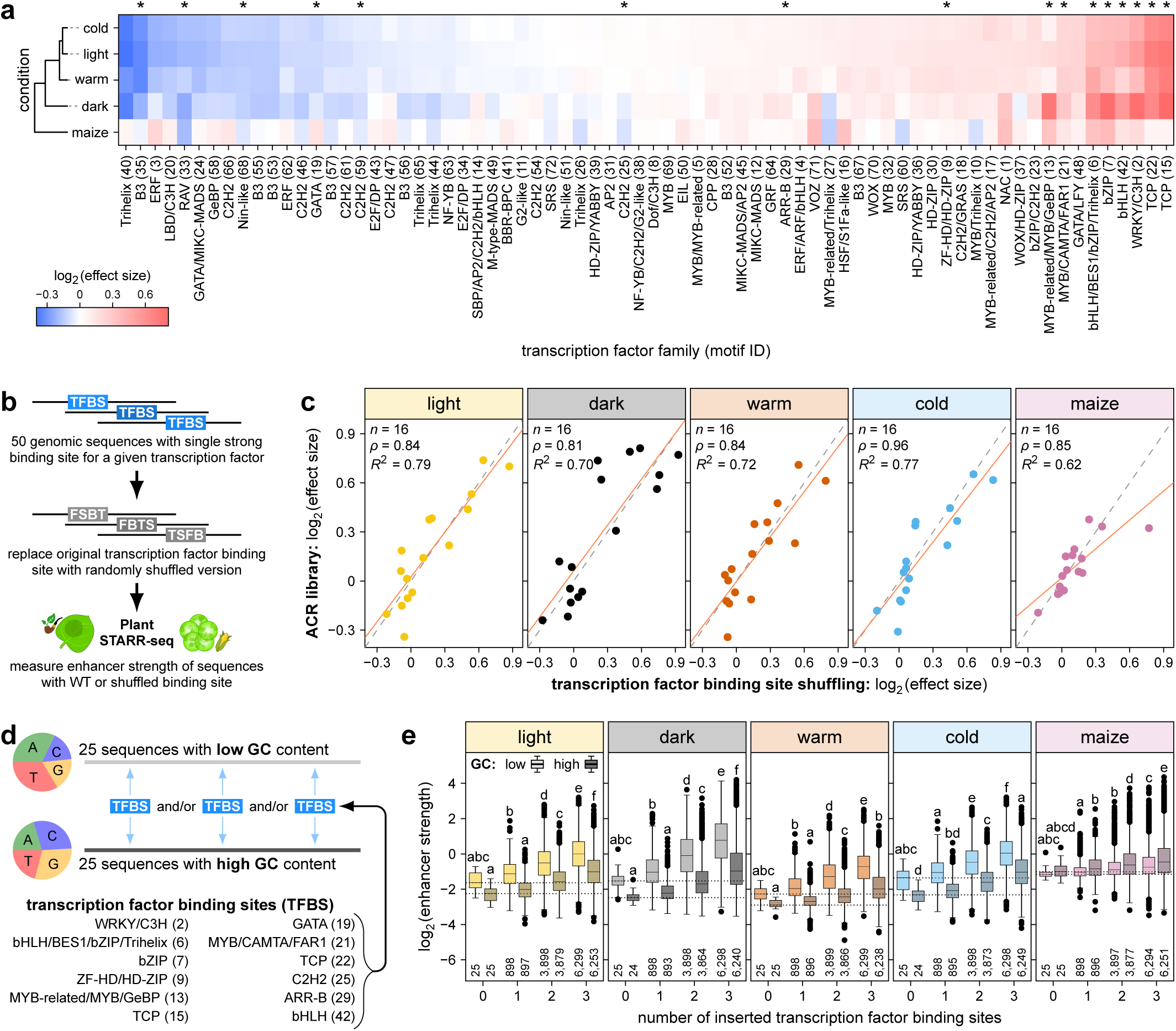
| Transcription factor binding sites are key determinants of enhancer strength. **a**, Putative transcription factor binding sites were identified by scanning the test sequences for 72 known binding motifs of the indicated transcription factor families. Effect size, calculated as the fold-change in GC-normalized (see Methods) enhancer strength between test sequences with or without one putative binding site, is indicated by color. **b**,**c**, To test the effect of a transcription factor binding site (TFBS), 50 test sequences containing a corresponding binding site were selected and mutated by replacing the binding site with a shuffled version. All sequences were subjected to Plant STARR-seq in the indicated conditions and species (**b**). The effect size of 16 different binding sites (marked with asterisks in **a**) was determined as the fold-change between the wild-type and mutated sequences, and was compared to the effect size in the ACR library (**c**). The solid and dashed lines represent a linear regression line and a *y* = *x* line, respectively. Pearson’s *R*^2^, Spearman’s *ρ*, and the number (*n*) of samples are indicated. **d**,**e**, Synthetic enhancers were designed by inserting one to three transcription factor binding sites into 25 random sequences each with a nucleotide composition similar to an average dicot (low GC) or monocot (high GC) ACR (**d**). The synthetic enhancers were subjected to Plant STARR-seq and their enhancer strengths are shown in box plots as defined in Fig. 1 (**e**). Letters above the box plots indicate significance groups determined by post-hoc Tukey tests performed separately for each condition. Exact *p* values are listed in Supplementary Data 4.

To validate our findings, we created a second Plant STARR-seq library containing sequences with either shuffled or added transcription factor binding sites. We selected 50 ACR sequences, each harboring a single strong binding site (*p* value ≤ 0.0001) for one of 16 different transcription factor families (marked with asterisk in Fig. 2a), and created mutated binding site variants by shuffling the respective DNA sequences (Fig. 2b). The loss of enhancer strength resulting from shuffling a given binding site was highly correlated with the effect size of the same binding site across all conditions and in both assay systems (Fig. 2c and Supplementary Figs. 7, 8, 9 and Supplementary Data 3). Inserting a binding site for one of 12 different transcription factor families into 50 random sequences increased enhancer strength (Extended Data Fig. 6).

### Rational design of synthetic enhancers

We sought to design synthetic enhancers based on 25 random sequence backgrounds, each with a nucleotide frequency resembling either an average dicot or monocot ACR sequence, and containing one, two or three transcription factor binding sites (Fig. 2d). Synthetic enhancers based on AT-rich, dicot-like random sequences showed higher strength in tobacco than their more GC-rich, monocot-like counterparts, whereas the opposite was true in maize (Fig. 2e). The strength of the synthetic enhancers increased with the number of inserted transcription factor binding sites (Fig. 2e). Synthetic enhancers with three binding sites were up to 140 times stronger than their corresponding random sequences. The strength of the synthetic enhancers could be predicted with a simple additive model based on the underlying random sequence and the individual effects of the inserted binding sites (*R*^2^ of 0.6 to 0.8; Extended Data Fig. 7). The position where a transcription factor binding site was inserted had only a modest effect on enhancer strength (Supplementary Fig. 10 and Extended Data Fig. 7b).

Because some of the transcription factor binding sites used to create synthetic enhancers showed species- or condition-specific differences in effect size (Fig. 2a), we asked if their insertion would suffice to yield enhancers with corresponding specificities. Synthetic enhancers with a tobacco-specific transcription factor binding site were more active in all tobacco conditions than they were in the maize system (Extended Data Fig. 8). However, synthetic elements with maize-specific or condition-specific transcription factor binding sites were weak and showed little evidence of species- or condition-specific activity (Extended Data Fig. 8b, c).

### The plantGREP model can predict enhancer strength and generate species- or condition-specific enhancers

To generalize our findings to genomic sequences, we developed computational models that predict enhancer strength. Models were trained with 90% of the sequences (304,042 unique sequences) and tested with the remaining 10%. Linear models based on transcription factor binding site counts or motif scores showed low predictive power (*R*^2^ of 0.2 to 0.4 in tobacco and 0.1 to 0.15 in maize) and systematically underestimated strong enhancers (log_2_(enhancer strength) > 2, Extended Data Fig. 9a–d, Extended Data Fig. 9e, Supplementary Fig. 11). A k-mer-based linear model performed better (*R*^2^ of 0.4 to 0.5 in tobacco and 0.3 in maize) but underpredicted strong enhancers (log_2_(enhancer strength) > 2, Extended Data Fig. 10).

Next, we built a convolutional neural network model (Fig. 3a) that uses a bidirectional convolutional layer to scan the input sequences with regular and reverse-complemented kernels. The outputs of this layer are fed into a DenseNet architecture^43^, which had success in predicting the strength of plant core promoters and terminators^44,45^. The resulting model—termed plantGREP (plant Gene Regulatory Element Predictor)—predicted enhancer strength well (*R*^2^ of 0.6 to 0.7 in tobacco and 0.5 in maize), including for strong enhancers (log_2_(enhancer strength) > 2, Fig. 3b).

**Fig. 3.**
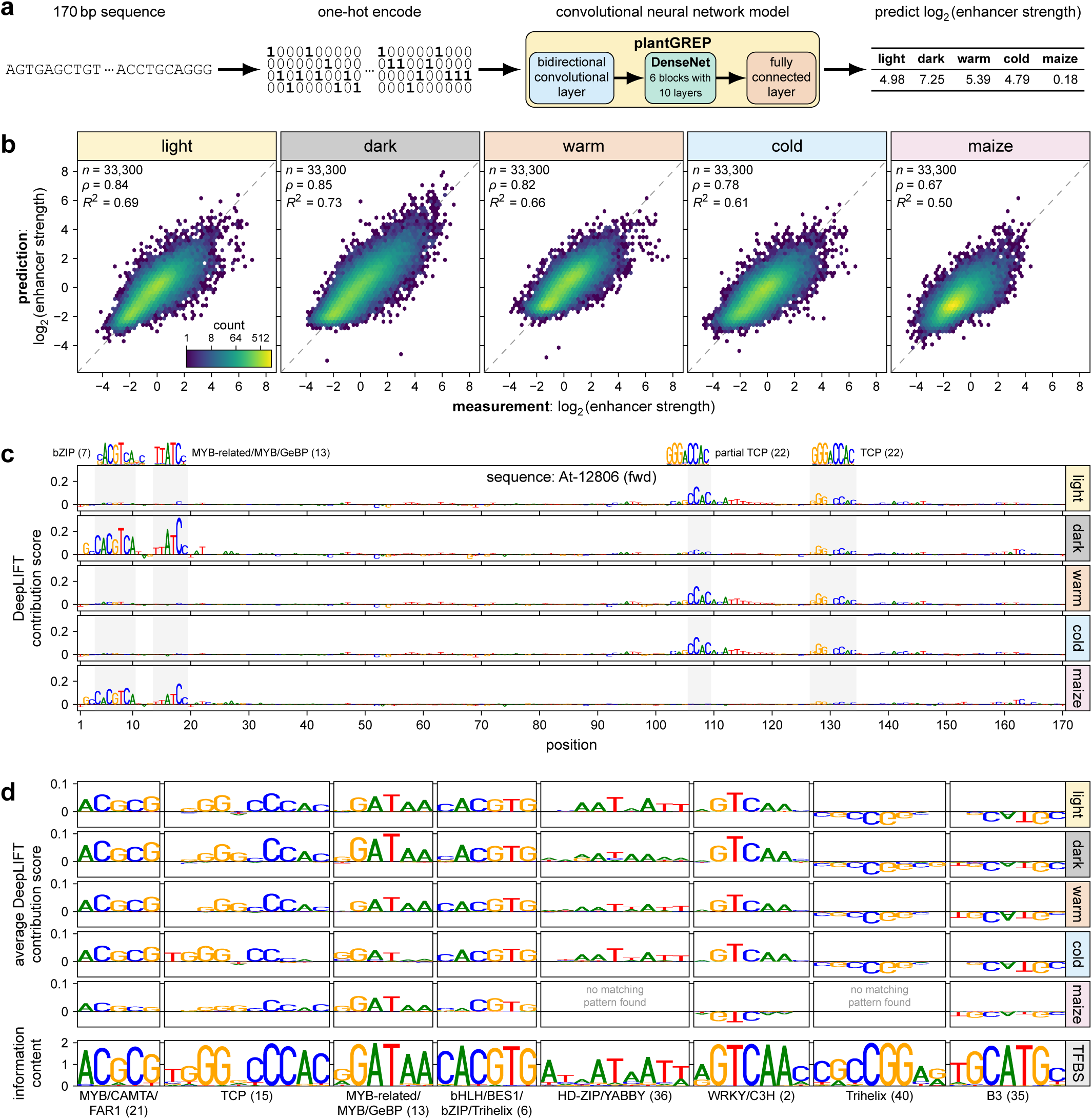
| A deep learning model predicts enhancer strength and identifies underlying features. **a**,**b**, A convolutional neural network model, termed plantGREP, consisting of a bidirectional convolutional layer followed by a DenseNet architecture with six blocks of ten layers each and a final fully connected layer was developed to predict enhancer strength across five conditions using one-hot encoded DNA sequences as input (**a**). The model was trained and validated on 90% of the data. The remaining 10% of the data (test set) were used to compare model predictions to the experimentally measured enhancer strength (**b**). The color in the hexbin plots indicate the number of observations in each hexagon and the dashed line represents a *y* = *x* line. Pearson’s *R*^2^, Spearman’s *ρ*, and the number (*n*) of samples are indicated. **c**, DeepLIFT was used to calculate contribution scores for an example ACR sequence (At-12806). Known transcription factor binding motifs that match the observed pattern are shown above the plot. **d**, TF-MoDISco was used to identify commonly occurring patterns in the DeepLIFT scores across all sequences of the test set. Examples of patterns detected by TF-MoDISco (top) and matching transcription factor binding motifs (TFBS; bottom) are shown. Patterns were truncated to the size of respective TFBSs.

To identify the features plantGREP learned, we used DeepLIFT, an explainable AI algorithm that attributes importance scores to all nucleotides in a sequence based on their contribution to the model output^46^. DeepLIFT analysis of an example ACR sequence showed cluster of nucleotides with high contribution scores that match known transcription factor binding motifs (Fig. 3c). We aggregated DeepLIFT contribution scores using TF-MoDISco^47^ to identify recurring patterns linked to enhancer strength. Most of the TF-MoDISco patterns (∼83%; 153 out of 184) matched known transcription factor binding sites (Fig. 3d and Supplementary Fig. 12–21).

We further improved prediction accuracy by averaging the predictions of an ensemble consisting of multiple (up to 50) plantGREP models trained with different, randomly initialized starting weights (*R*^2^ of up to 0.75 in tobacco and 0.55 maize). However, increasing the number of models in the ensemble showed diminishing returns while linearly increasing the required computational resources (Supplementary Fig. 22).

Taken together, plantGREP accurately predicts the enhancer strength of novel sequences and identifies the underlying functional sequence motifs. The trained model is available in a Google Colab notebook (https://colab.research.google.com/github/tobjores/plantGREP/blob/main/code/plantGREPcolab/plantGREPcolab.ipynb) and command-line interface (https://github.com/tobjores/plantGREP/tree/main/code/plantGREPcli).

We applied plantGREP to create novel enhancers by *in silico* evolution, scoring all possible single-nucleotide substitution variants of a sequence. The highest-scoring variant was used as the starting point for the next evolution round, with each iteration introducing a single mutation predicted to improve enhancer strength (Fig. 4a). We subjected 150 sequences (50 random sequences and 100 randomly selected sequences from the ACR library) to twelve rounds of *in silico* evolution. The starting sequences, and the round six and round twelve evolved sequences, were tested experimentally. *In silico* evolution increased enhancer strength up to 250-fold and resulted in several enhancers with high activity (log_2_(enhancer strength) > 3) in both tobacco and maize, under all tested conditions (Fig. 4b). Introducing multiple mutations per round did not yield enhancers of greater strength but substantially increased computational demands (Supplementary Fig. 23).

**Fig. 4.**
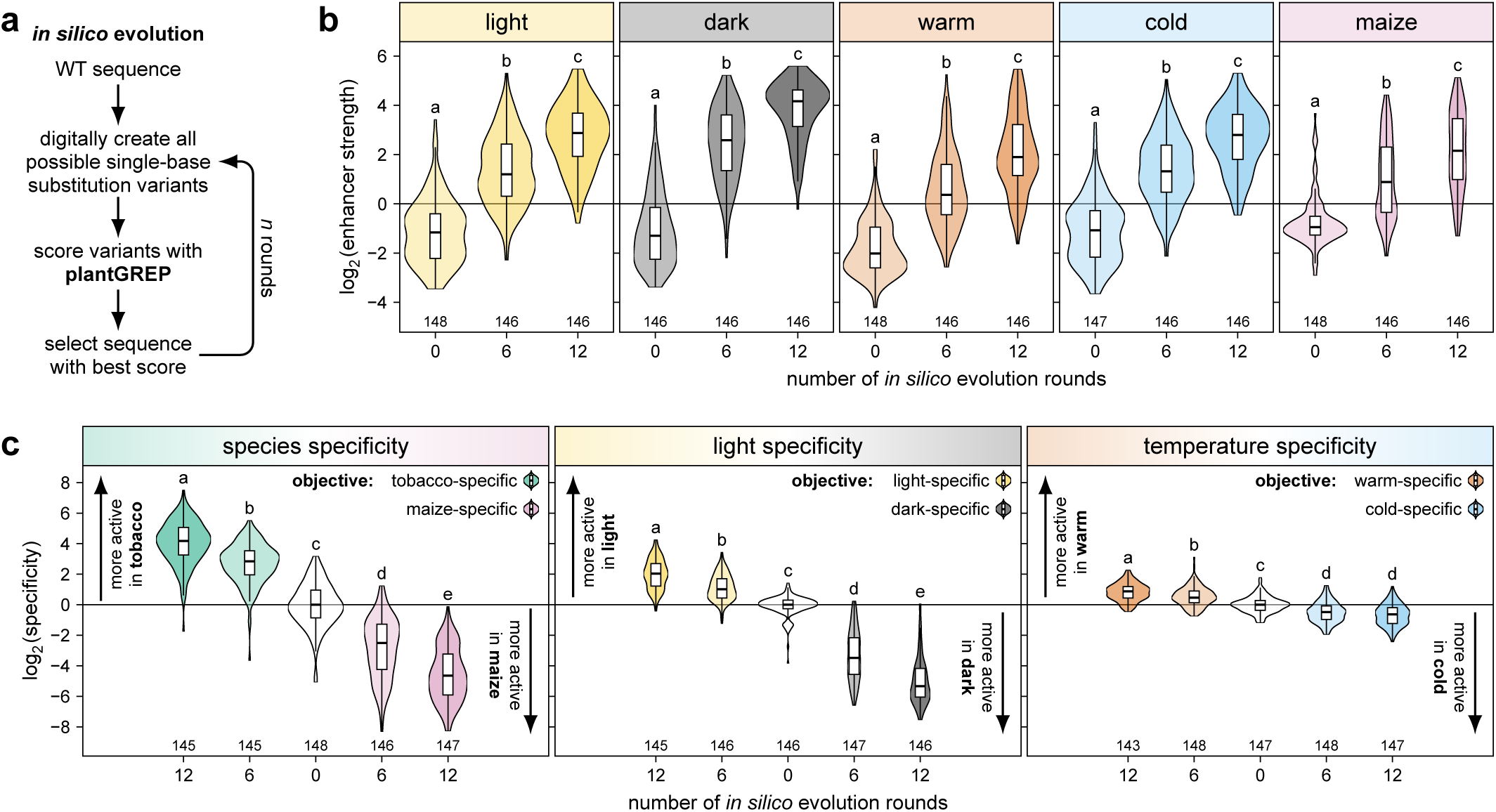
| Building species- and condition-specific synthetic enhancers through *in silico* evolution. **a**–**c**, Improved enhancers were generated through *in silico* evolution using plantGREP (**a**). The starting sequences and sequences evolved for six or twelve rounds were subjected to Plant STARR-seq in the indicated conditions and species (**b**,**c**). The objective was to evolve constitutive enhancers (**b**) or species- or condition-specific enhancers (**c**). Species and condition specificity was calculated as the fold-change in enhancer strength between two species (tobacco [mean of light, dark, warm and cold] and maize) or conditions (light and dark for light specificity; warm and cold for temperature specificity). Violin plots in (**b**) and (**c**) are as defined in Fig. 1 and letters above the violin plots indicate significance groups determined by post-hoc Tukey tests performed separately for each condition, species, or specificity. Exact *p* values are listed in Supplementary Data 4.

We tested whether modifying the *in silico* scoring function could generate species-or condition-specific enhancers, rather than only constitutive ones. Indeed, when scoring was based on predicted species specificity (the fold-change in enhancer strength between tobacco and maize), only with the target species did the evolved enhancers show very high strength (up to log_2_(enhancer strength) > 5). Similarly, enhancers evolved for light- or dark-specific activity exhibited the desired condition-specific activity (Fig. 4c and Supplementary Fig. 24). However, the light specificity achieved was substantially lower than the dark specificity (19-fold vs. 188-fold higher activity in the target condition). This result is consistent with our finding that transcription factor binding sites contribute additively to plant enhancer activity in the dark but require cooperativity in the light^26^.

Enhancer strength in warm and cold conditions was highly correlated (Supplementary Fig. 2) and transcription factor binding sites did not show specificity for either temperature (Fig. 2a). Nonetheless, enhancers evolved for temperature-specific activity showed up to 5-fold higher strength in the targeted temperature (Fig. 4c), indicating that plantGREP recognizes even subtle differences in condition-specific enhancer activity.

### Identification of likely enhancers in crop genes

We asked whether Plant STARR-seq could identify candidate enhancer regions for engineering in crop genomes. First, we focused on a study by Wang *et al.*, in which CRISPR-Cas9 mutagenesis was used to introduce deletions into the upstream region of the tomato *CLAVATA3* (*SlCLV3*) gene^18^. A ∼1-kb deletion just upstream of the *SlCLV3* start codon did not affect phenotype, while a deletion of similar size further upstream resulted in larger tomato fruits due to reduced *SlCLV3* expression (Fig. 5a). We used Plant STARR-seq to measure the enhancer strength of 170 bp overlapping fragments from the *SlCLV3* upstream region. The fragments with the highest enhancer strength resided within the distal region whose deletion results in larger fruits (Fig. 5b). We obtained similar results when scoring the fragments using plantGREP predictions (Fig. 5b), suggesting that this strategy could be broadly applicable to any gene, even in the absence of Plant STARR-seq data.

**Fig. 5.**
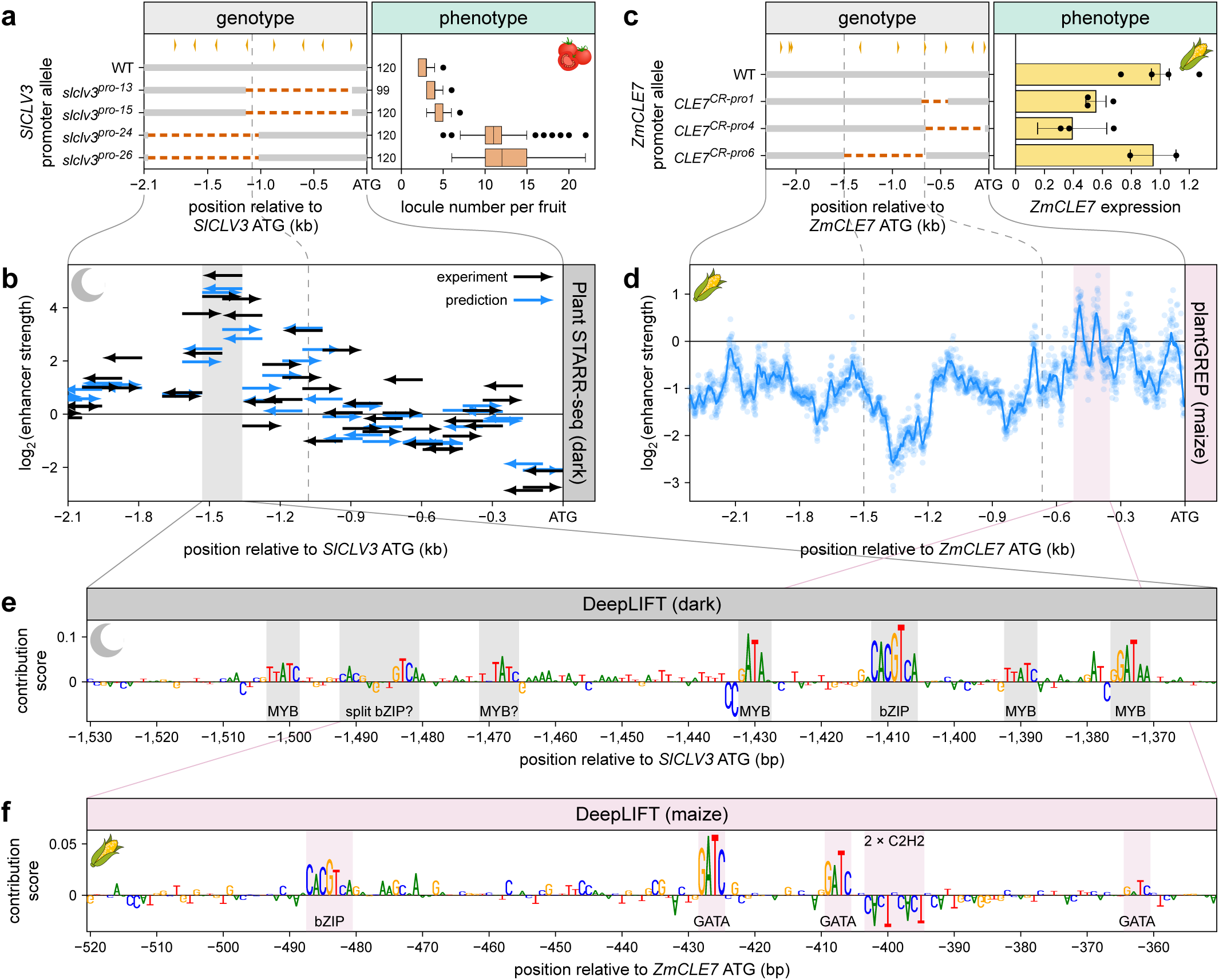
| Plant STARR-seq data and models can guide genome engineering of regulatory elements. **a**, Wang *et al.* used CRISPR-Cas9 to introduce deletions in the upstream region of *SlCLV3* and measured the number of locules per tomato fruit^18^. Increased numbers of locules are indicative of decreased *SlCLV3* expression levels. **b**, The enhancer strength in the dark of fragments from the *SlCLV3* upstream region was determined experimentally by Plant STARR-seq (black arrows) and predicted using plantGREP (blue arrows). The orientation of the tested fragments is indicated by the direction of the arrows. **c**, Liu *et al.* used CRISPR-Cas9 to introduce deletions in the upstream region of *ZmCLE7* and measured its expression levels^19^. **d**, The enhancer strength in maize of all possible 170-bp fragments from the *ZmCLE7* upstream region was predicted with plantGREP. Points show individual predictions and the line represents a sliding average (window size 20 bp). **e**,**f**, DeepLIFT contribution scores for regions with enhancer activity (shaded in **b** and **d**) of the *SlCLV3* (**e**) and *ZmCLE7* (**f**) upstream regions were calculated with plantGREP for the indicated conditions. Matches to known transcription factor binding sites are shaded and labeled. Data in (**a**) and (**c**) was obtained from ref. ^18^ and ref. ^19^, respectively.

To test this hypothesis, we turned to a study by Liu *et al.* that introduced deletions into the upstream region of the maize *CLAVATA3*/*EMBRYO SURROUNDING REGION-RELATED7* (*ZmCLE7*) gene^19^. *ZmCLE7* expression was reduced in response to deletions within a ∼700-bp region directly upstream of the start codon (Fig. 5c). Using plantGREP, we predicted the enhancer strength of every possible 170-bp fragment across the *ZmCLE7* upstream region. The strongest fragments were located within the critical ∼700-bp region required for *ZmCLE7* expression (Fig. 5d).

Finally, we used DeepLIFT to identify functional sequence motifs that drive the expression of *SlCLV3* and *ZmCLE7*. The contribution scores for the upstream regions with the highest enhancer strength showed several clusters of nucleotides matching known binding motifs for MYB (*SlCLV3*), bZIP (*SlCLV3* and *ZmCLE7*), and GATA (*ZmCLE7*) family transcription factors (Fig. 5e,f). Because genome engineering in crops remains laborious and time-consuming, applying plantGREP could direct these efforts towards promising candidate loci.

## Discussion

This work confirms that plant enhancers share key features with animal enhancers: their activity is independent of orientation and driven by transcription factor binding. Sequences that show higher chromatin accessibility in the genomic context—presumably due to more consistent or higher protein occupancy—showed higher enhancer strength in the Plant STARR-seq assay. At a deeper level, our study revealed plant-specific nuances in the features of enhancers and ACRs that might reflect differences between plants and animals in genome organization and gene regulation. For example, sequences showing the highest enhancer strength in the Plant STARR-seq assay tended to reside close to the core promoter and transcription start site. While there are examples of distal gene regulation in plants, such as the *tb1* enhancer in maize^11,48,49^, more common is promoter-proximal gene regulation. The reason could be the much greater prevalence of transposons in plants (Groover et al., 2026), which are more likely to disrupt distal enhancer-promoter interactions than proximal ones. Consistent with promoter-proximal gene regulation, introns in plant genomes are much smaller than in animal genomes^15^. Introns, particularly the first intron, commonly contain regulatory elements^11,50^.

Enhancer strength was primarily driven by the additive action of transcription factor binding sites, which increased enhancer strength regardless of their position. However, our use of consensus transcription factor binding sites might have diminished the effects of flanking bases. Doubling and tripling the number of transcription factor binding sites generated synthetic enhancers comparable in strength to the commonly used viral 35S enhancer, and these data were captured by a simple additive model, reminiscent of studies in yeast^51^.

However, the additive models failed to account for enhancer condition specificity and for the activity of strong enhancers (log_2_(enhancer strength) > 2). Using plantGREP, we could generate enhancers that showed strong and specific activity in the light or dark. We previously dissected the activity in the dark and light of three plant enhancers by saturation mutagenesis, mutation-sensitive domain shuffles and chimeras. In the dark, enhancer strength adhered to the “billboard” model, with transcription factors binding independently and acting additively^51,52^. In contrast, in the light, the order and spacing of the mutation-sensitive domains likely bound by transcription factors significantly affected enhancer activity, consistent with the “enhanceosome” model which assumes cooperativity^53,54^. Thus, plantGREP appears to learn some aspects of transcription factor cooperativity. However, *in silico* evolution with plantGREP produced far higher specificity for dark-specific enhancers, suggesting that the difference between enhancer modes in the dark and light is not fully captured in the model.

The success of generating strong species- and condition-specific enhancers suggests an experimental strategy for building cell-type-specific plant enhancers. Single-cell and spatial genomics approaches can be used to identify the transcription factors present and ACR-bound in distinct plant cell types^24,55–58^. Because plant transcription factor families tend to be large and contain members with highly similar binding preferences^39–42^, insertion of single binding sites into random sequence backgrounds is unlikely to yield cell-type specificity. However, insertion of binding sites for two or three different cell-type-specific transcription factors could feasibly generate expression specificity in major plant cell- or tissue-types, especially with plants having far fewer terminally differentiated cell- and tissue-types than animals^15,59^. Moreover, specificity could be enhanced by adding binding sites for repressors not expressed in the target cell or tissue type. Order and spacing of the binding sites will likely matter and would need to be determined experimentally. However, this approach would greatly decrease the search space compared to current approaches that rely on conservation and laborious genomic edits.

Over 40% of the tested sequences failed to drive gene expression in any condition or assay system. This result reflects the weak correlation between chromatin accessibility and gene expression in both plant bulk and single-cell genomics experiments^15,23,50,56,60^. The constant integration of environmental cues with gene expression in plants requires that many ACRs be poised for activation: occupied by trans-acting factors but not triggering transcription until a signal is perceived. This mode of transcriptional regulation is likely not captured in our experiments.

About a third of the tested sequences repressed the activity of the core promoter in at least one condition or assay system (log_2_(enhancer strength) < -2). This result might reflect the action of repressive transcription factors, and the sequences might act as enhancers in other tissues, as in animals^8^. Alternatively, the sequences that repressed the activity of the core promoter may act as silencers, a class of regulatory elements whose in-depth exploration in plants would require a different experimental set-up than used here.

In summary, context—both genomic (sequences) and cellular (transcription factor repertoire)—contributes to the enhancer activity of ACR sequences more than often assumed from chromatin accessibility assays and MPRA approaches. In humans, the vast majority of the over one million trait- and disease-associated variants reside in ACRs^61,62^. However, with few exceptions, efforts to tie these ACR-residing variants to changes in gene expression have failed^63–66^. Combining STARR-seq with multiplexed expression of specific transcription factors, approaches like Fiber-seq^67–69^ that detect protein footprints on single molecules, and modeling could shed light on the mechanisms that underpin the different ACR effects.

## Methods

### Library design and construction

Candidate sequences for the ACR library were derived from publicly available accessibility data for Arabidopsis^23^, tomato^17^, maize^24^, and sorghum^25^. We used 170-bp sequences surrounding the center of all accessibility peaks as candidate sequences for our library. Candidate sequences overlapping with a core promoter (defined as the region from −60 to +5 relative to the annotated transcription start site) were shifted upstream of the core promoter. Sequences, coordinates, and genomic features of the ACR sequences are based on the Araport11 (Arabidopsis; ref. ^70^), B73_RefGen_v4.50 (maize; ref. ^71^), or Sorghum_bicolor_NCBIv3.50 (sorghum; ref. ^72^) genomes and annotations. For tomato, we used a preliminary version of the M82 genome and annotation supplied by Xingang Wang and Zachary B. Lippman^73^. For maize, the annotated transcription start sites were replaced with experimentally determined ones where available^74^.

To validate and extend our findings from the ACR library, we created a second, smaller library (validation library). This library included ACR sequences with a single high-confidence (*p* value ≤ 0.0001) binding site for one of 16 different transcription factor families (marked with an asterisk in Fig. 2a; 50 sequences per family) and variants thereof where the binding site was randomly shuffled. Additionally, the validation library contained synthetic enhancers created by inserting one to three binding sites for twelve different transcription factor families into 25 random sequences each with a nucleotide composition similar to an average dicot (32% A, 34% T, 16% C, and 18% G) or monocot (20% A, 21% T, 29% C, and 30 % G) ACR. Binding sites were inserted with their center at positions 29, 85, and/or 141 (numbered starting from the 5′ end) of the random sequences. Finally, the validation library also contained the sequences from *in silico* evolution (see below).

All libraries used in this study were constructed using pPSnt, a variant of pPSup (Addgene no. 149416; https://www.addgene.org/149416/; ref. ^28^), as the base plasmid. The plasmid’s T-DNA region harbors a GFP reporter construct terminated by the poly(A) site of the Cauliflower Mosaic Virus (CaMV) 35S RNA. Two versions of pPSnt were created to receive enhancer candidates in the forward (pPSntF) or reverse (pPSntR) orientation by changing the BsaI scars to ACTC and CTGT or ACAG and GAGT, respectively. The plasmids were deposited at Addgene (Addgene no. 256230 and 256231; https://www.addgene.org/256230/; https://www.addgene.org/256231/). The CaMV 35S minimal promoter (−46 to +5 relative to the 35S transcription start site) followed by the 5′ UTR from a maize histone H3 gene (Zm00001d041672) and an 18-bp random barcode (VNNVNNVNNVNNVNNVNN; V = A, C, or G) downstream of an ATG start codon was cloned in front of the second codon of GFP by Golden Gate cloning^75^ using BbsI-HF (NEB). To distinguish between the pPSntF and pPSntR versions of the library, positions 7 and 13 of the barcodes were set to fixed bases. Enhancer candidates were ordered as an oligo pool from Twist Biosciences and inserted into the pPSnt plasmids by Golden Gate cloning using BsaI-HFv2 (NEB). The resulting libraries were bottlenecked to yield on average about 10 (ACR library) or 20 (validation library) barcodes per enhancer candidate.

The base plasmids for dual-luciferase constructs were derived from pDL (Addgene no. 208978; https://www.addgene.org/208978/; ref. ^26^) by changing the BsaI scars to ACTC and CTGT (pDLenhF; to test sequences in the forward orientation) or ACAG and GAGT (pDLenhR; to test sequences in the reverse orientation). The plasmids were deposited at Addgene (Addgene no. 256232 and 256233; https://www.addgene.org/256232/; https://www.addgene.org/256233/). Enhancer candidates were ordered as synthesized gene fragments from Twist Biosciences and inserted into pDLenh plasmids by Golden Gate cloning using BsaI-HFv2 (NEB).

The sequences of the oligonucleotides used in this study for cloning and sequencing are listed in Supplementary Table 5.

### Tobacco cultivation and transformation

Tobacco (*N. benthamiana*) was grown in soil (Sunshine Mix no. 4) at 25 °C in a long-day photoperiod (16 h light and 8 h dark; cool-white fluorescent lights [Philips TL-D 58W/840]; intensity 300 μmol m^−2^ s^−1^). Plants were transformed approximately 3 weeks after germination. For transient transformation of tobacco leaves, Plant STARR-seq libraries were introduced into Agrobacterium tumefaciens strain GV3101 (harboring the virulence plasmid pMP90 and the helper plasmid pMisoG) by electroporation. An overnight culture of the transformed A. tumefaciens was diluted into 200 mL YEP medium (1% [w/v] yeast extract and 2% [w/v] peptone) and grown at 28 °C for 8 h. A 10-mL input sample of the cells was collected, and plasmids were isolated from it using the QIAprep Spin Miniprep Kit (Qiagen) according to the manufacturer’s instructions. The remaining cells were harvested and resuspended in 200 mL induction medium (M9 medium [3 g/L KH_2_PO_4_, 0.5 g/L NaCl, 6.8 g/L Na_2_HPO_4_, and 1 g/L NH_4_Cl] supplemented with 1% [w/v] glucose, 10 mm MES, pH 5.2, 100 μM CaCl_2_, 2 mm MgSO_4_, and 100 μM acetosyringone). After overnight growth, the Agrobacteria were harvested, resuspended in infiltration solution (10 mm MES, pH 5.2, 10 mm MgCl_2_, 150 μm acetosyringone, and 5 μm lipoic acid) to an optical density of 1 and infiltrated into leaves 3 and 4 of 24 (ACR library) or 12 (validation library) tobacco plants. The plants were further grown for 48 h at 25 °C under normal light (16 h light and 8 h dark; “light” condition) or in complete darkness (“dark” condition) before mRNA extraction. For the temperature treatments, the plants were first grown for 48 h under normal conditions (16 h light and 8 h dark at 25 °C) followed by 24 h with the same light settings but increased (35 °C; “warm” condition) or decreased (10 °C; “cold” condition) temperature.

### Maize cultivation and transformation

To transform maize (*Zea mays* L. cultivar B73) protoplasts for Plant STARR-seq, we used the PEG transformation method as previously described^76^. Maize seeds were germinated in soil at 25 °C in a long-day photoperiod (16 h light and 8 h dark; cool-white fluorescent lights [Philips TL-D 58W/840]; intensity 300 μmol m^−2^ s^−1^). After 3 days, the seedlings were moved to complete darkness at 25 °C and grown for 10 to 11 days. From each seedling, 10 cm sections from the second and third leaves were cut into thin 0.5 mm strips perpendicular to veins and immediately submerged in 10 mL of protoplasting enzyme solution (0.6 M mannitol, 10 mm MES pH 5.7, 15 mg/mL cellulase R10, 3 mg/mL macerozyme, 1 mm CaCl_2_, 0.1% [w/v] BSA, and 5 mm β-mercaptoethanol). The mixture was covered in foil to keep out light, vacuum infiltrated for 3 min, and incubated on a shaker at 40 rpm for 2.5 h. Protoplasts were released by incubating an extra 10 min at 80 rpm. To quench the reaction, 10 mL ice-cold MMG (0.6 M mannitol, 4 mm MES pH 5.7, 15 mm MgCl_2_) was added to the enzyme solution and the whole solution was filtered through a 40 µm cell strainer. To pellet protoplasts, the filtrate was split into equal volumes of no more than 10 mL in chilled round-bottom glass centrifuge vials and centrifuged at 100 × *g* for 4 min at room temperature (RT). Pellets were resuspended in 1 mL cold MMG each and combined into a single round-bottom vial. To wash, MMG was added to make a total volume of 5 mL and the solution was centrifuged at 100 × *g* for 3 min at RT. This wash step was repeated 2 more times. The final pellet was resuspended in 1 to 2 mL of MMG. A sample of the resuspended protoplasts was diluted 1:20 in MMG and used to count the number of viable cells using fluorescein diacetate as a dye. For each replicate, 5 to 10 million protoplasts were mixed with 15 µg per million protoplasts of the Plant STARR-seq plasmid library in a fresh tube, topped with MMG to a volume of 114.4 µL per million protoplasts, and incubated on ice for 30 min. For PEG transformation, 105.6 µL per million protoplasts of PEG solution (0.6 m Mannitol, 0.1 m CaCl_2_, 25% [w/v] poly-ethylene glycol MW 4000) was added to reach a final concentration of 12% (w/v) PEG. The mixture was incubated for 10 min in the dark at RT. After incubation, the transformation solution was diluted with 5 volumes of incubation solution (0.6 m Mannitol, 4 mm MES pH 5.7, 4 mm KCl), and centrifuged at 100 × *g* for 4 min at RT. The protoplast pellet was washed with 5 mL of incubation solution, centrifuged at 100 × *g* for 3 min at RT, and resuspended in incubation solution to a concentration of 500 cells/µL. Protoplasts were incubated overnight in the dark at RT to allow for transcription of the plasmid library and then pelleted (4 min, 100 × *g*, RT). The pellet was washed with 1 to 5 mL incubation solution and centrifuged (3 min, 100 × *g*, RT). The pellet was resuspended in 1 to 5 mL incubation solution. An aliquot of the solution was used to check transformation efficiency under a microscope. Cells were pelleted (4 min, 100 × *g*, RT) and resuspended in 12 mL Trizol for subsequent mRNA extraction. An aliquot of the plasmid library used for PEG transformation was used as the input sample for Plant STARR-seq.

### Arabidopsis cultivation and transformation

*Arabidopsis thaliana* Col-0 was grown in soil (Sunshine Mix no. 4) at 20 °C in a long-day photoperiod (16 h light and 8 h dark; cool-white fluorescent lights [Sylvania FO32/841/ECO 32W]; intensity 100 μmol m^−2^ s^−1^). For transformation, dual-luciferase plasmids were introduced into Agrobacterium tumefaciens strain GV3101 (harboring the virulence plasmid pMP90 and the helper plasmid pMisoG) by electroporation. Transgenic *Arabidopsis* plants were generated by floral dipping^77^ and selected for by spraying with a 0.01% (w/v) glufosinate solution. Successful genome integration of the dual-luciferase constructs was verified by genotyping PCR.

### Plant STARR-seq

For all Plant STARR-seq experiments, at least 2 independent biological replicates were performed. Different plants and fresh *Agrobacterium* cultures were used for each biological replicate.

For Plant STARR-seq in tobacco, whole leaves were harvested two (light and dark conditions) or three (warm and cold conditions) days after infiltration and partitioned into batches of 8 leaves. The leaf batches were frozen in liquid nitrogen, finely ground with mortar and pestle, and immediately resuspended in 20 mL QIAzol (Qiagen). The suspensions were cleared by centrifugation (5 min, 4,000 × *g*, 4 °C). The supernatant was transferred to a 50-mL MaXtract High Density tube (Qiagen) and mixed with 5 mL chloroform. After centrifugation (10 min, 1,000 × *g*, 4 °C), another 5 mL chloroform were added. The samples were mixed and centrifuged (10 min, 1,000 × *g*, 4 °C) again. The supernatant (approximately 14 mL) was poured into a new tube, and mixed with 7 mL 100% ethanol. The solution from three leaf batches was loaded onto a RNeasy Maxi column (Qiagen) in 15 mL steps. The column was washed according to the instructions of the RNeasy Maxi prep kit (Qiagen) and the RNA was eluted with 1.2 mL and 1 mL nuclease-free water in two subsequent elution steps. The eluates were pooled (yielding approximately 2 mL) and mixed with 112 µL 20X DNase I buffer (1 mM CaCl_2_, 100 mM Tris pH 7.4), 112 µL 200 mM MnCl_2_, 12.5 µL DNase I (Thermo Fisher Scientific), and 2.5 µL RNaseOUT (Thermo Fisher Scientific). After 1 h incubation at 37 °C, the reaction was stopped with 250 µL 500 mM EDTA. To precipitate the RNA, the solution was split into five 2-mL tubes (approximately 0.5 mL each), and mixed with 250 µL high salt buffer (0.8 M sodium citrate, 1.2 M NaCl) and 250 µL isopropanol. After incubation for 15 min at room RT, the RNA was pelleted by centrifugation (20 min, 20,000 × *g*, 4 °C). The pellet was washed with 1 mL ice-cold 70% ethanol, centrifuged (5 min, 20,000 × *g*, 4 °C), air-dried, and resuspended in 100 µL nuclease-free water. All batches of the same sample were pooled, and the solution was supplemented with 1 µL RNaseOUT.

For cDNA synthesis, 8 (ACR library) or 4 (validation library) reactions with 11 µL RNA solution, 1 µL 10 µm GFP-specific reverse transcription primer, and 1 µL 10 mM dNTPs were incubated at 65 °C for 5 min then immediately placed on ice. The reactions were supplemented with 4 µL 5X SuperScript IV buffer, 1 µL 100 mM DTT, 1 µL RNaseOUT, and 1 µL SuperScript IV reverse transcriptase (Thermo Fisher Scientific). To ensure that the samples were largely free of DNA contamination, 4 additional reactions were used as controls, where the reverse transcriptase and RNaseOUT were replaced with water. Reactions were incubated for 10 min at 55 °C, followed by 10 min at 80 °C. Sets of 4 reactions each were pooled and mixed with 4 µL RNase H (NEB). After 20 min incubation at 37 °C, the cDNA was purified with the Clean&Concentrate-5 kit (Zymo Research), and eluted in 25 µL 10 mm Tris. The barcodes portion of the cDNA was amplified with 10 to 20 cycles of polymerase chain reaction (PCR) and read out by next-generation sequencing.

For Plant STARR-seq in maize protoplasts, the protoplast-containing Trizol solution from PEG transformation was transferred to 2-mL Phasemaker tubes (1 mL per tube; Thermo Fisher Scientific), mixed thoroughly with 300 µL chloroform, and centrifuged (5 min, 15,000 × *g*, 4 °C). RNA was extracted using the RNeasy Plant Mini Kit (Qiagen). The supernatant from two Phasemaker tubes was transferred to a QIAshredder column and centrifuged (2 min, 20,000 × *g*, RT). The flowthrough was transferred to a new 1.5-mL tube and mixed with 300 µL 100% ethanol. This process was repeated with the same QIAshredder column and the supernatant from a second Phasemaker tube. The two flowthroughs were pooled and up to 500 µL of the solution was loaded on an RNeasy mini spin column. After centrifugation (10 s, 16,100 × *g*, RT) the flowthrough was discarded. This was repeated until the whole solution had been added to the column. The column was washed with 350 µL RW1 buffer followed by centrifugation (30 s, 16,100 × *g*, RT). An on-column DNase I digestion was performed with 70 µL RDD buffer and 10 µL DNase I (Qiagen) for 15 min at RT. The column was washed once with 350 µL RW1 buffer and twice with 500 µL RPE buffer. After each wash step, the column was centrifuged (30 s, 16,100 × *g*, RT) and the flowthrough was discarded. The column was dried with an extra centrifugation step (30 s, 16,100 × *g*, RT) and transferred to a 1.5-mL collection tube. For elution, 50 µL of RNase-free water was added, and the column was incubated for 1 min, and centrifuged (1 min, 16,100 × *g*, RT). This elution step was repeated with an additional 40 µL of RNase-free water. The eluate was treated with DNase I (5 µL of 20x DNaseI buffer, 5 µL 200 mm MnCl_2_, 1 µL RNaseOUT, and 2 µL DNase I) for 1 h at 37 °C. The solution was supplemented with 20 µL 500 mm EDTA, 1 µL 20 mg/mL glycogen, 12 µL ice-cold 8 m LiCl, and 300 µL ice-cold 100% ethanol. The solution was incubated 15 min at −80 °C and centrifuged (20 min, 20,000 × *g*, 4 °C). The pellet was washed with 500 µL ice-cold 70% ethanol and centrifuged (3 min, 20,000 × *g*, 4 °C). The pellet was air-dried and resuspended in 50 µL RNase-free water. All samples from the same experiment were pooled and subjected to reverse transcription, purification, PCR amplification, and sequencing as for the tobacco samples.

### Dual-luciferase assay

Transgenic Arabidopsis lines (T2 generation) with dual-luciferase constructs were grown in soil for 3 weeks. A cork borer (4 mm diameter) was used to collect a total of 4 leaf discs from the third and fourth leaves of the plants. The leaf discs were transferred to 1.5-mL tubes filled with approximately 10 glass beads (1 mm diameter), snap-frozen in liquid nitrogen, and disrupted by shaking twice for 5 s in a Silamat S6 (Ivoclar) homogenizer. The leaf disc debris was resuspended in 100 µL 1X Passive Lysis Buffer (Promega). The solution was cleared by centrifugation (5 min, 20,000 × *g*, RT) and 10 µL of the supernatant were mixed with 90 µL 1X passive lysis buffer. Luciferase and nanoluciferase activity were measured on a Biotek Synergy H1 plate reader using the Promega Nano-Glo Dual-Luciferase Reporter Assay System according to the manufacturer’s instructions. Specifically, 10 µL of the leaf extracts were combined with 75 µL ONE-Glo EX Reagent, mixed for 3 min at 425 rpm, and incubated for 2 min before measuring luciferase activity. Subsequently, 75 µL NanoDLR Stop&Glo Reagent were added to the sample. After 3 min mixing at 425 rpm and 12 min incubation, nanoluciferase activity was measured. Three independent biological replicates were performed.

### Subassembly and barcode sequencing

All sequencing was performed on an Illumina NextSeq 2000 system. To link insulator fragments to their respective barcodes, the insulator region was sequenced using paired-end reads (151 bp), and 2 18-bp indexing reads were used to sequence the barcodes. For each Plant STARR-seq experiment, barcodes were sequenced using 18-bp paired-end reads. Paired reads were assembled using PANDAseq (version 2.11; ref. ^78^). Insulator fragment-barcode pairs with less than 5 reads and insulator fragments with a mutation or truncation were discarded.

### Computational methods

For analysis of the Plant STARR-seq experiments, the reads for each barcode were counted in the input DNA and cDNA samples. Barcode counts below 5 were discarded. Barcode counts were normalized to the sum of all counts in the respective sample. The sum of the normalized counts for all barcodes associated with a given enhancer candidate was used to calculate its enhancer strength. For each replicate, the enhancer strength was normalized to the median enhancer strength. The mean enhancer strength across all replicates was normalized to a control construct without an enhancer and used for all analyses.

Spearman and Pearson’s correlation were calculated using base R (version 4.4.1). Linear regression analysis was performed using the lm() function in base R with default parameters (e.g. lm(*y* ∼ *x*)). The base R function t.test() was used to calculate 95% confidence intervals. To test for statistic significance, the base R functions aov(), TukeyHSD(), wilcox.test(), and pairwise.wilcox.test() were used. The *p* values obtained from Wilcoxon tests were corrected for multiple comparisons using the Bonferroni method as implemented in p.adjust() in base R. Exact *p* values are listed in Supplementary Data 4. Principal component analysis was performed on normalized replicate data using the prcomp() function with options ‘center’ and ‘scale.’ set to TRUE. All data for figures was processed in R and exported for plotting in LaTeX using the pgfplots package.

ACR sequences were linked to genomic features using BEDTools (version 2.29.2; ref. ^79^) and BEDOPS (version 2.4.35; ^80^). Specifically, the tool ‘closest-features’ was used to extract the closest transcription start site for each ACR sequence, and ‘bedmap’ was used to identify the region (intergenic, 5 UTR, CDS, intron, 3 UTR, or ncRNA) in which the center of an ACR sequence resides. Histone modifications occurring in a 1-kb window surrounding the center of *Arabidopsis* ACR sequences were attributed to that sequence using ‘bedmap’ and publicly available histone ChIP data from ReMap2022 (ref. ^81^).

Processed gene expression data for *Arabidopsis*^36^, tomato^33^, maize^34^, and sorghum^35^ was obtained from the EMBL-EBI Expression Atlas^37^. Additionally, *Arabidopsis* genes differently expressed in the light and dark were obtained from ref. ^82^. ACR sequences were linked to genes based on the closest transcription start site. For each gene, only the ACR sequence with the highest enhancer strength was retained and genes with less than 10 FPKM were discarded. The correlation between enhancer strength (log_2_-transformed) and gene expression (log_10_-transformed FPKM) was calculated in each condition and species for all available RNA-seq samples. To calculate the specificity of enhancer strength across the tested conditions and species or of gene expression across the samples tissues, the tau index was calculated as previously published^38^. For GO term enrichment analysis, GO terms were linked to ACRs based on GO term annotations of the closest gene. GO term enrichment analysis was performed using the ggprofiler2 (version 0.2.3; ref. ^83^) library for R and a custom gmt file with GOslim terms.

Putative transcription factor binding sites were identified in the ACR sequences using the scan_sequences() function of the universalmotif (version 1.22.2; ref. ^84^) library and 72 consensus transcription factor binding motifs^39^ (Supplementary Table 4). Both strands of the ACR sequences were searched for matches to the consensus transcription factor binding motifs and matches with a *p* value below 0.0001 were counted. For overlapping matches, only the highest-scoring match was counted. To calculate the cumulative match scores, both strands of the ACR sequences were searched for matches to the consensus transcription factor binding motifs and the scores of all matches with a logodds score above 0 were summed up. The effect size of transcription factor binding sites in the ACR library was calculated based on enhancer strength that was GC-normalized by fitting a linear model (formula: log_2_(enhancer strength) ∼ GC content) for each condition. The fitted GC content trend was then subtracted from the observed log_2_(enhancer strength).

### Computational modelling of promoter strength

To predict the enhancer strength of the ACR sequences, we built linear models and a convolutional neural network in python (version 3.10.12). The models were trained on a set of 80% of all measured ACR sequences and tested against a held-out set of 10% of the ACR sequences (hereafter referred to as the test set). For the convolutional neural network, the remaining 10% of the ACR sequences was used as a validation set during model training. The two orientations of a given ACR sequence were always kept in the same set. All models were tasked to predict log_2_-transformed enhancer strength.

Linear models were built using LinearRegression() from the scikit-learn (version 1.2.0; ref. ^85^) library. For each assay condition and species, three separate linear models were trained using counts of transcription factor binding sites, cumulative scores of transcription factor binding motif matches, or counts of all possible 6-mers as input features.

The convolutional neural network model was built using EUGENe (version 0.1.2; ref. ^86^) and PyTorch (version 2.4.0; ^87^). The model takes 170-bp long, one-hot encoded DNA sequences as an input which is fed to a convolutional layer with 32 filters and a kernel size of 9. The input is also scanned twice with a bidirectional convolutional layer with 256 filters and a kernel size of 9. Before the second pass, the filters of the bidirectional layer are reverse-complemented and the maximum activation of the two passes is combined with the output of the first convolutional layer. This is followed by six dense blocks consisting of ten convolutional layers with 24 filters each and a kernel size of 3. In each dense block, the output of a convolutional layer is appended to its input and the combined output is used as the input for the subsequent layer. Between each dense block we inserted a transition layer that reduces the output feature map in size through convolution (using half as many filters as the input and a kernel size of 1) and average pooling (with a kernel size of 2). Each dense block and transition layer was preceded by a batch normalization layer. The output of the final dense block is fed into a fully connected layer with the output corresponding to the enhancer strength in the different assay conditions and species. To avoid a bias caused by the higher number of data from tobacco (four conditions versus only one for maize), the data from the three individual replicate experiments for maize were added as additional model outputs for a total of 8 outputs (light, dark, warm, cold, maize[mean], maize[replicate 1], maize[replicate 2], and maize [replicate 3]). After training, the model was truncated to yield only the first five outputs (light, dark, warm, cold, maize). Mean squared error was used as a loss function during model training with the Adam optimizer^88^ using an initial learning rate of 0.001, which was reduced by a factor of 2 if the validation loss did not improve for two epochs. Model training was stopped after the validation loss did not improve for five epochs and the model weights after the epoch with the lowest validation loss were used for the final model.

Contribution scores for all nucleotides in a given 170-bp DNA sequence were calculated using the DeepLIFT method^46^ implemented in captum (version 0.5.0; ref. ^89^). We observed substantial differences in the contribution scores for the same sequence when it was placed in different positions within the batch of sequences subjected to DeepLIFT. Therefore, for each sequence contribution scores were calculated at least ten times with the sequence in a different batch position each time, and the average score across all positions was used as the final contribution score. The DeepLIFT contribution scores for all sequences in the test set were used as input for TF-MoDISco^47^ as implemented in modisco-lite (version 2.2.1) to extract motifs that positively or negatively contribute to the model predictions.

### In *silico* evolution

We used plantGREP to improve enhancer performance in an iterative fashion. In each round, we generated all possible single-nucleotide variants of a given 170-bp enhancer sequence, scored them with plantGREP and kept the variant with the highest score for the next round. In a parallel approach, we created variant sequences for *in silico* evolution by introducing two or three mutations within an 8-bp window. These sequence variants were scored in the same way as the single-nucleotide variants. Different objective functions were used to score the sequences during *in silico* evolution. Enhancers were evolved for high activity in each individual condition or species (light, dark, warm, cold, or maize) by scoring variants with the corresponding plantGREP prediction. To evolve constitutive enhancers, they were scored using the sum of all plantGREP predictions. Species-specific enhancers were evolved by using the difference between the prediction for maize and the sum of the predictions for tobacco as the score. The difference between predictions for light and dark or warm and cold conditions was used as a score to evolve light- or temperature-specific enhancers, respectively.

As starting sequences for the *in silico* evolution, we randomly chose 100 ACR sequences from the test set used to assess the performance of plantGREP. Additionally, the 50 random sequences used to build synthetic enhancers (see above) were also subjected to *in silico* evolution. Finally, we used Plant STARR-seq to experimentally determine the enhancer strength of the original sequences and the evolved variants with six or twelve mutations.

## Data availability

The raw sequencing data underlying this article are available in the National Center for Biotechnology Information (NCBI) Sequence Read Archive at http://www.ncbi.nlm.nih.gov/bioproject/1427589. The processed data underlying this article are available on GitHub at https://github.com/tobjores/plantGREP. The plantGREP model can be used through a Google Colab notebook at https://colab.research.google.com/github/tobjores/plantGREP/blob/main/code/plantGREPcolab/plantGREPcolab.ipynb and through a command-line interface available at https://github.com/tobjores/plantGREP/tree/main/code/plantGREPcli.

## Code availability

The code used for the analysis and to generate the figures is available on GitHub at https://github.com/tobjores/plantGREP.

## Acknowledgements

We thank Xingang Wang and Zachary B. Lippman for supplying us with a preliminary version of the M82 genome and annotation.

This work was supported by the National Science Foundation (RESEARCH-PGR grant no. 1748843 to S.F. and C.Q. and PlantSynBio grant no. 2240888 to C.Q.), the German Research Foundation (DFG; postdoctoral fellowship no. 441540116 to T.J., Emmy Noether program grant no. 517938232 to T.J., and Germany’s Excellence Strategy - EXC-2048/1 - project ID 390686111 to T.J.), the National Institutes of Health (T32 training grant no. HG000035 to J.T., NIGMS grant no. R01-GM079712 to J.T.C. and C.Q., and NIGMS MIRA grant no. 1R35GM139532 to C.Q.), and the United States Department of Agriculture (NIFA postdoctoral fellowship no. 2023-67012-39445 to N.A.M.).

## Author contributions

All authors conceived and interpreted experiments; T.J., N.A.M., S.G., and J.T. performed experiments; T.J., S.G., S.T., D.S., and K.L.B. analyzed the data; T.J. prepared figures; T.J., S.F., and C.Q. wrote the manuscript. All authors read and revised the manuscript.

## Competing interests

The authors declare no competing interests.

---

**Extended Data Fig. 1.**
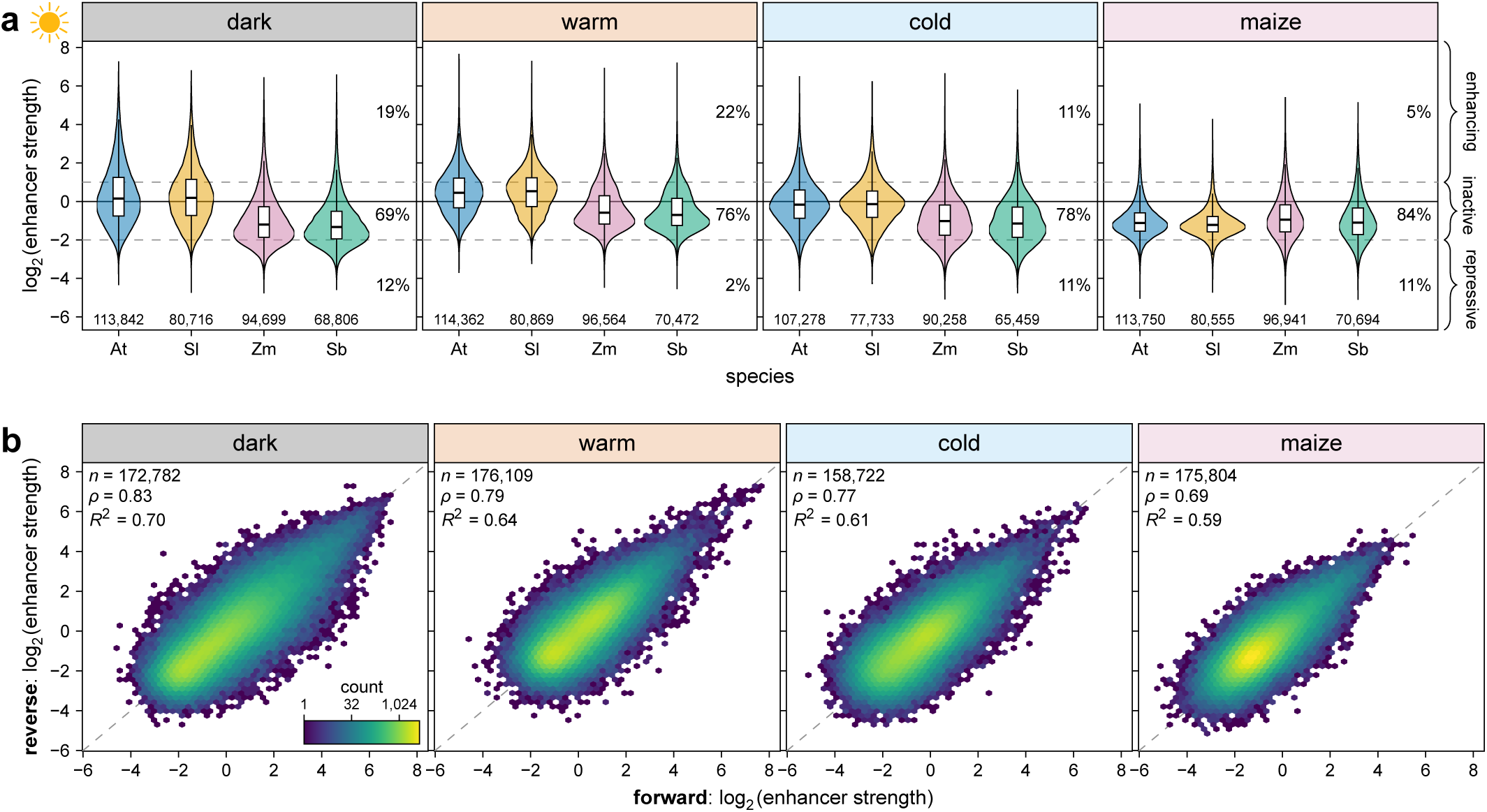
| Regulatory activity spans large dynamic range and is orientation-independent. **a**, Violin plots (as defined in Fig. 1) of the enhancer strength of test sequences grouped by species of origin. The percentage of enhancing (log_2_ (enhancer strength) > 1), inactive (log_2_ (enhancer strength) between 1 and −2), and repressive (log_2_ (enhancer strength) < −2) sequences is indicated. **b**, Hexbin plots (as defined in Fig. 1) for the correlation of enhancer strength between sequences tested in the forward and reverse orientation.

**Extended Data Fig. 2.**
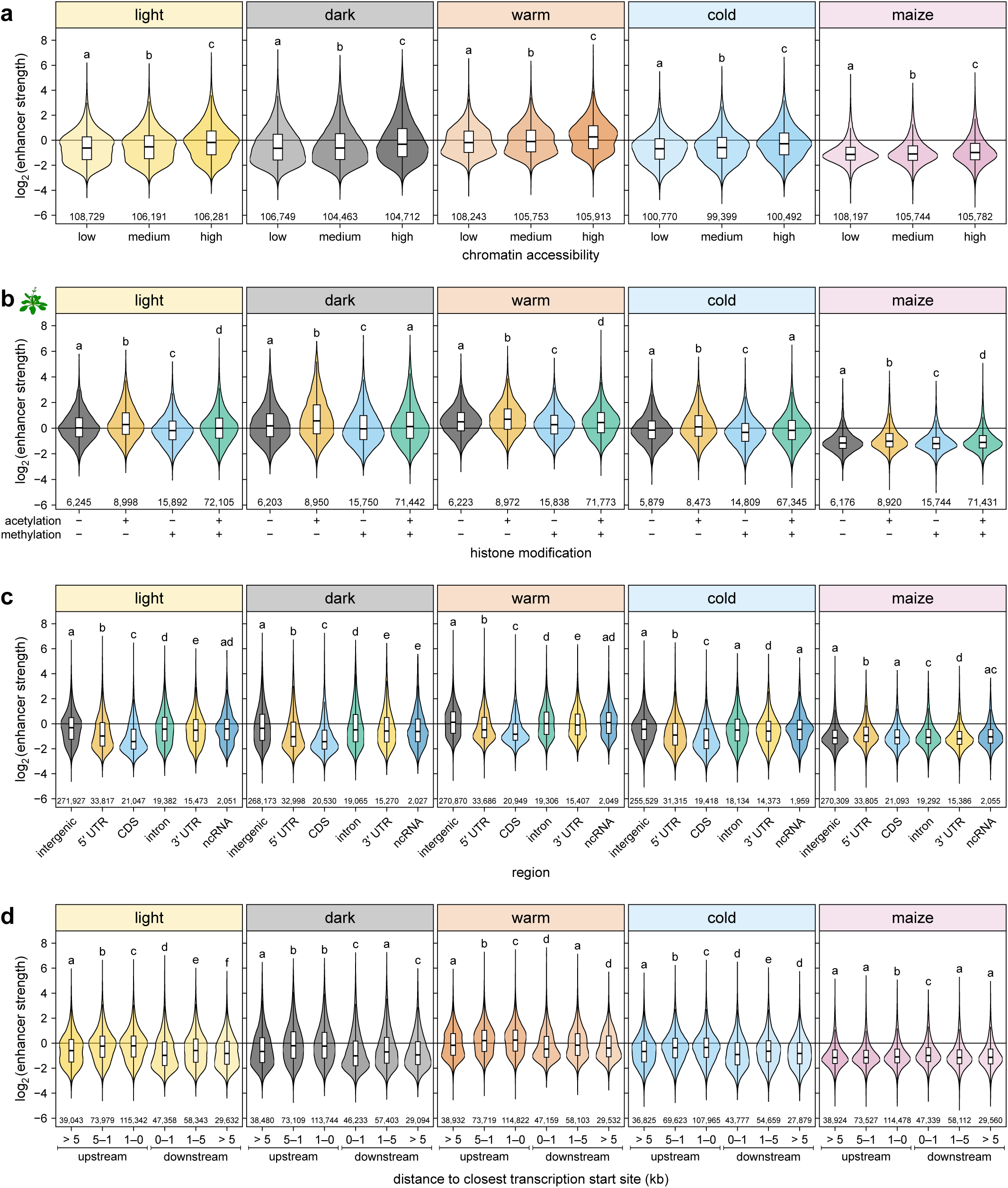
| Genomic signatures of enhancers. **a**–**d**, Enhancer strength of test sequences grouped by the relative accessibility of the corresponding region in the genome (**a**), by the occurrence of acetylated and/or methylated histones in a 1-kb window surrounding the corresponding region in the genome (**b**), by the genomic region from which they are derived (**c**), or by their distance to the closest transcription start site (**d**). Violin plots are as defined in Fig. 1 and letters above the plots indicate significance groups determined by post-hoc Tukey tests performed separately for each condition and species. Exact *p* values are listed in Supplementary Data 4. UTR, untranslated region; CDS, coding sequence; ncRNA, non-coding RNA.

**Extended Data Fig. 3.**
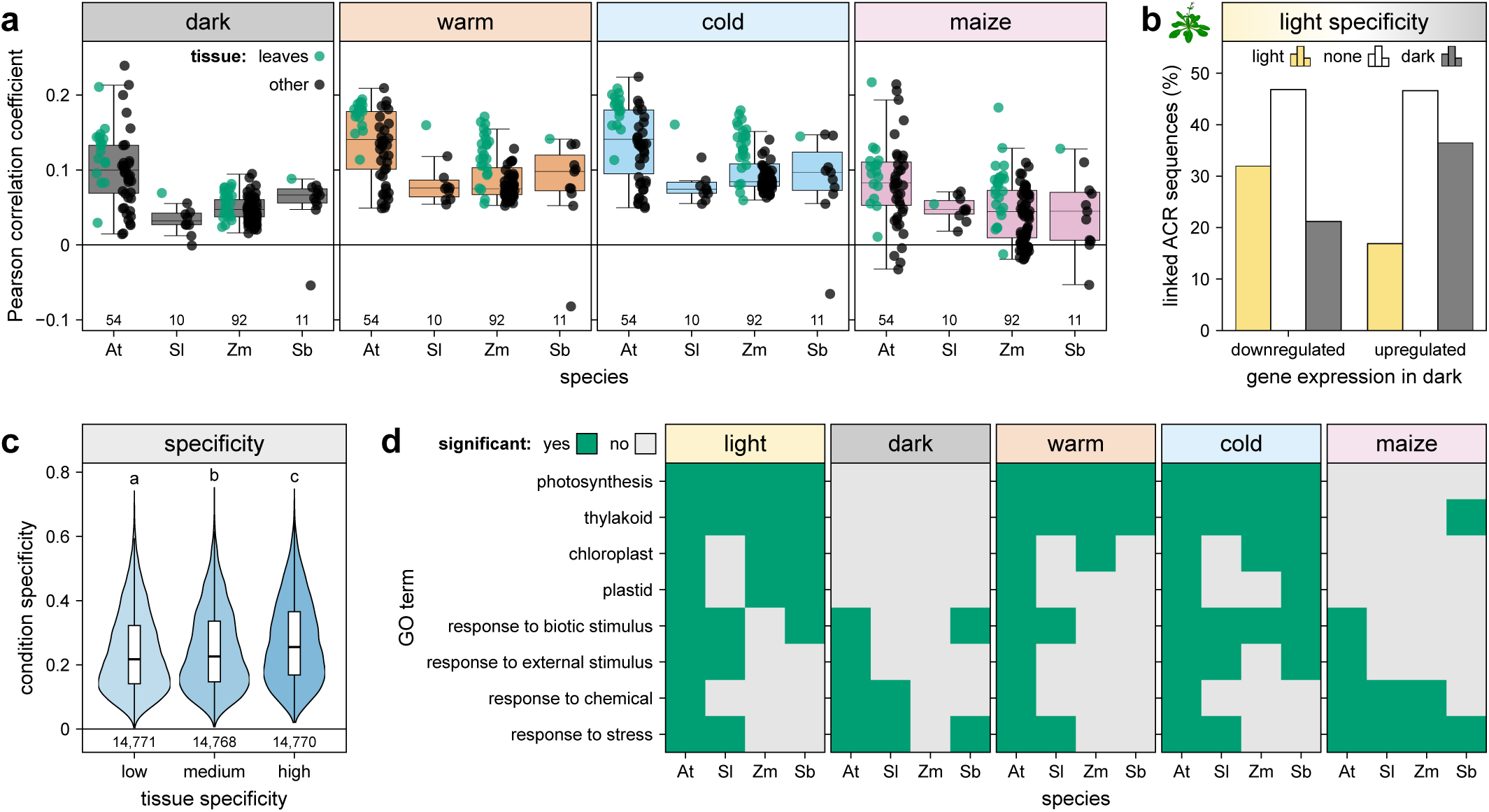
| Enhancer strength is correlated with gene expression *in planta*. **a**, Correlation between enhancer strength and gene expression in various tissues of the indicated species. At, *Arabidopsis*; Sl, tomato; Zm, maize; Sb, sorghum. ACR sequences were linked to the closest transcription start site. For each gene, only the ACR sequence with the highest enhancer strength was retained. **b**, For genes down- or upregulated after shifting *Arabidopsis* plants to the dark, the percentage of linked ACR sequences showing light-specific (1.5× stronger in the light than in the dark; light), dark-specific (1.5× stronger in the dark than in the light; dark), or constitutive (none) enhancer activity is indicated. **c**, Condition-specific enhancer activity of ACR sequences grouped by the relative tissue specificity of the linked genes. Condition and tissue specificity was determined using the tau index based on our Plant STARR-seq data and the gene expression data used in (**a**), respectively. For each gene, only the ACR sequence with the highest condition specificity was retained. Violin plots are as defined in Fig. 1 and letters above the plots indicate significance groups determined by a post-hoc Tukey test. Exact *p* values are listed in Supplementary Data 4. **d**, GO term enrichment for genes linked to the top 5% of test sequences with the highest enhancer strength in the indicated conditions and species. Significant (adjusted *p* value ≤ 0.05) enrichment is indicated in green; non-significant results are in gray. See Supplementary Data 2 for all GO terms and *p* values. Gene expression data was obtained from ref. ^33–36^.

**Extended Data Fig. 4.**
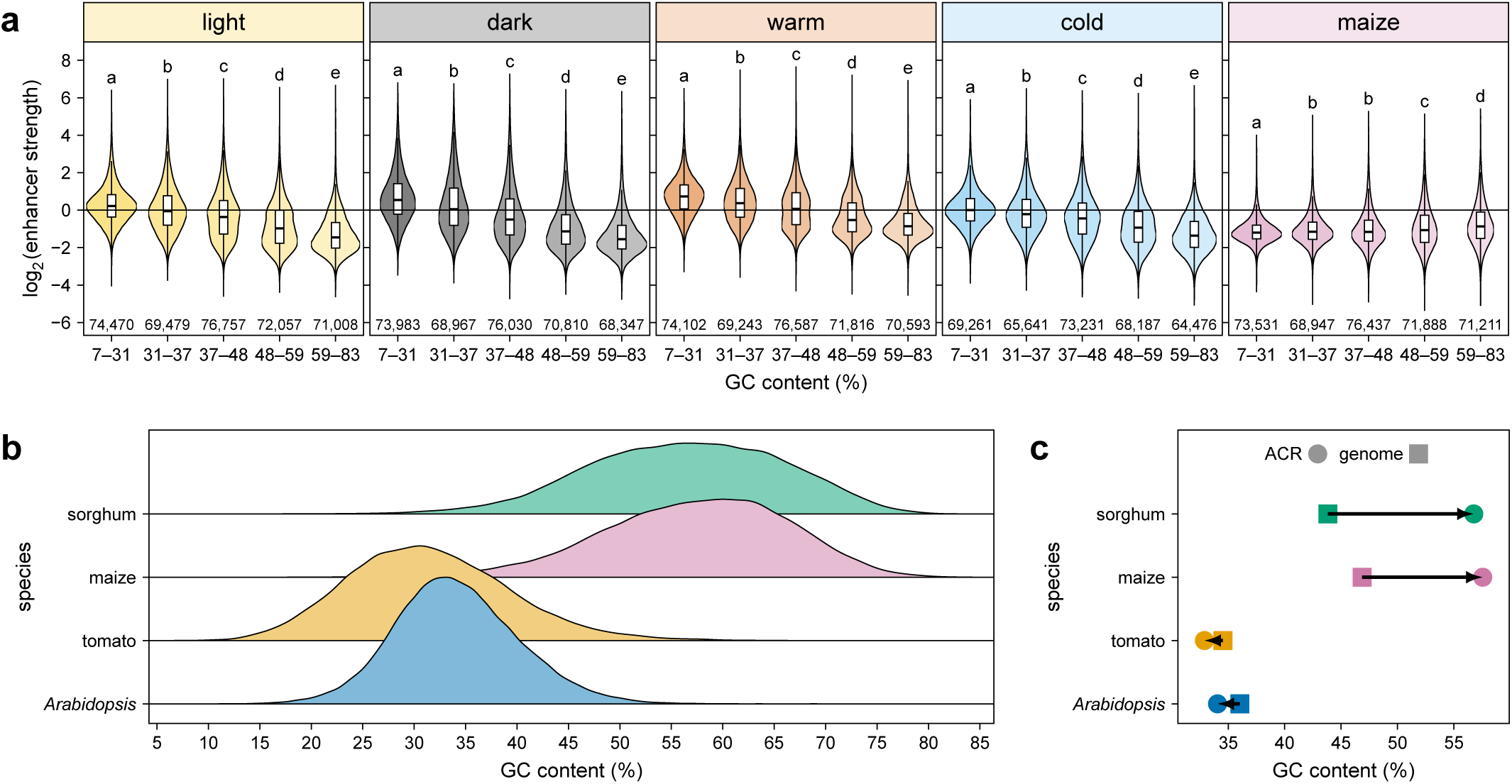
| GC content affects enhancer strength in tobacco. **a**, Violin plots (as defined in Fig. 1) of the enhancer strength of test sequences grouped by GC content. Letters above the plots indicate significance groups determined by post-hoc Tukey tests performed separately for each condition and species. Exact *p* values are listed in Supplementary Data 4. **b**, Distribution of GC content for the test sequences in the ACR library. **c**, Average GC content of the ACR sequences (circles) and the whole genome (squares) of the indicated species.

**Extended Data Fig. 5.**
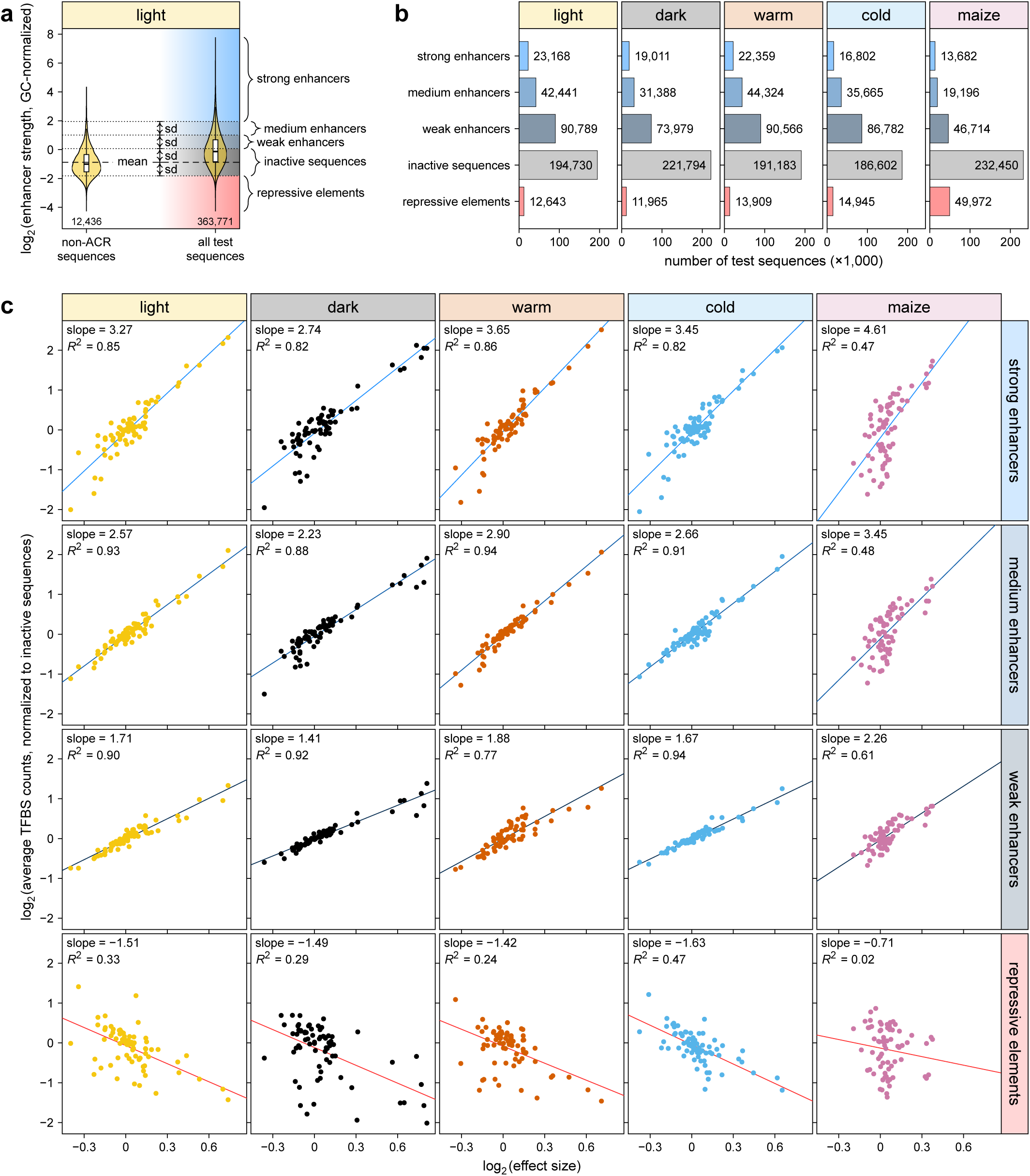
| Activating transcription factor binding sites are enriched in active enhancers. **a**,**b**, Test sequences in the ACR library were categorized based on the mean (mean) and standard deviation (sd) of the GC-normalized enhancer strength of non-ACR sequences. The categorization scheme is shown for the light condition as an example (**a**). The number of sequences in each category is indicated for all conditions and assay systems (**b**). **c**, The average count of putative transcription factor binding sites (TFBS) in the test sequences of each category was normalized to the average count in inactive sequences and compared to the effect size of the binding site as determined in Fig. 2a.

**Extended Data Fig. 6.**
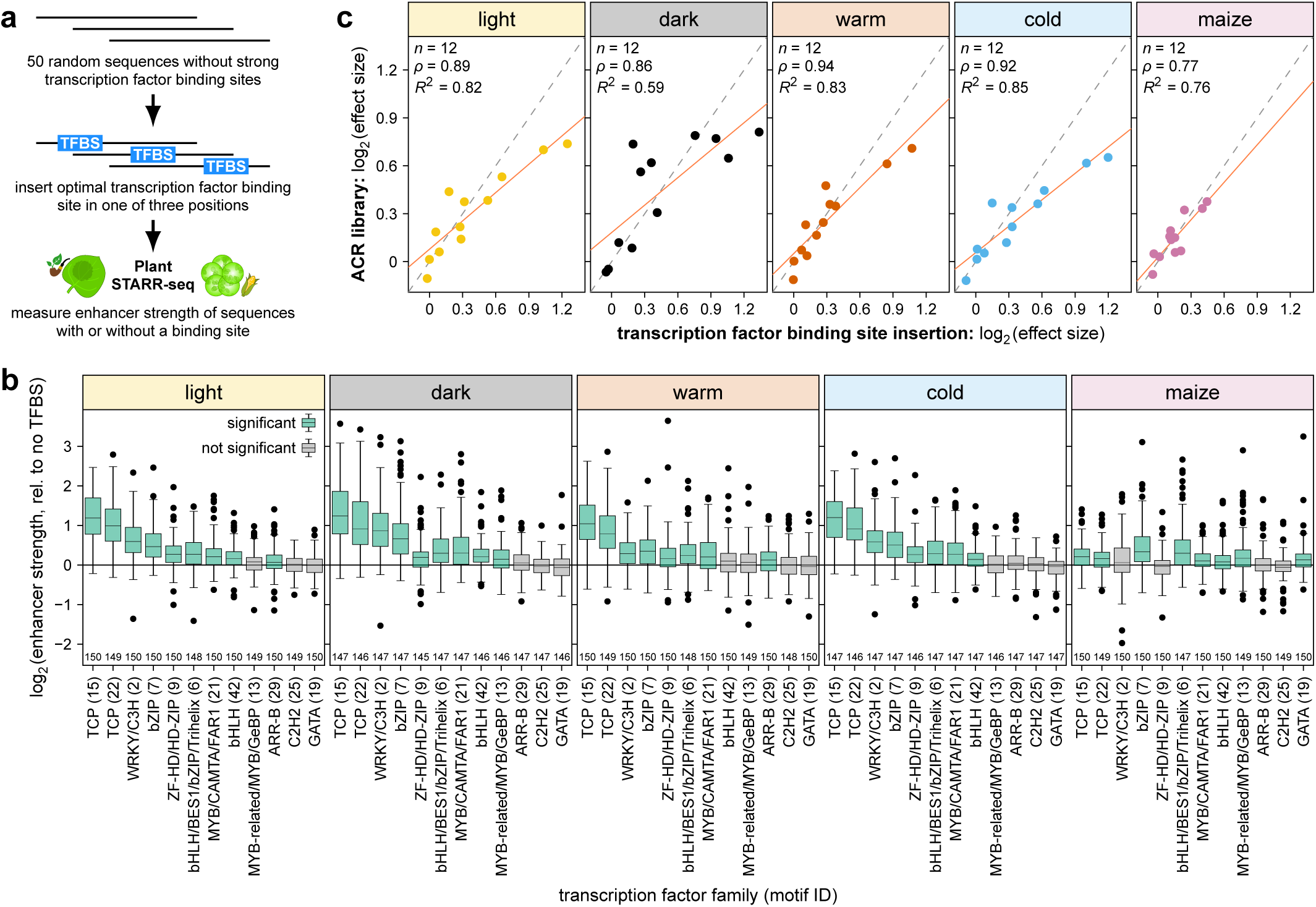
| Inserted transcription factor binding sites can increase enhancer strength. **a**–**c**, A transcription factor binding site (TFBS) was inserted into 50 random sequences and the sequences were subjected to Plant STARR-seq (**a**) to determine their enhancer strength relative to the corresponding random sequence without a binding site (no TFBS) (**b**). The effect size of each binding site was determined as the fold-change between the sequences with and without it, and was compared to the effect size in the ACR library (see Fig. 2b) (**c**). The solid and dashed lines represent a linear regression line and a *y* = *x* line, respectively. Pearson’s *R*^2^, Spearman’s *ρ*, and the number (*n*) of samples are indicated. Box plots in (**b**) are as defined in Fig. 1. Significant differences from a null distribution were determined using one-sample Wilcoxon rank-sum tests. Box plots for significant groups (Bonferroni-adjusted *p* value ≤ 0.05) are shown in green. Non-significant groups are represented by gray box plots. Exact *p* values are listed in Supplementary Data 4.

**Extended Data Fig. 7.**
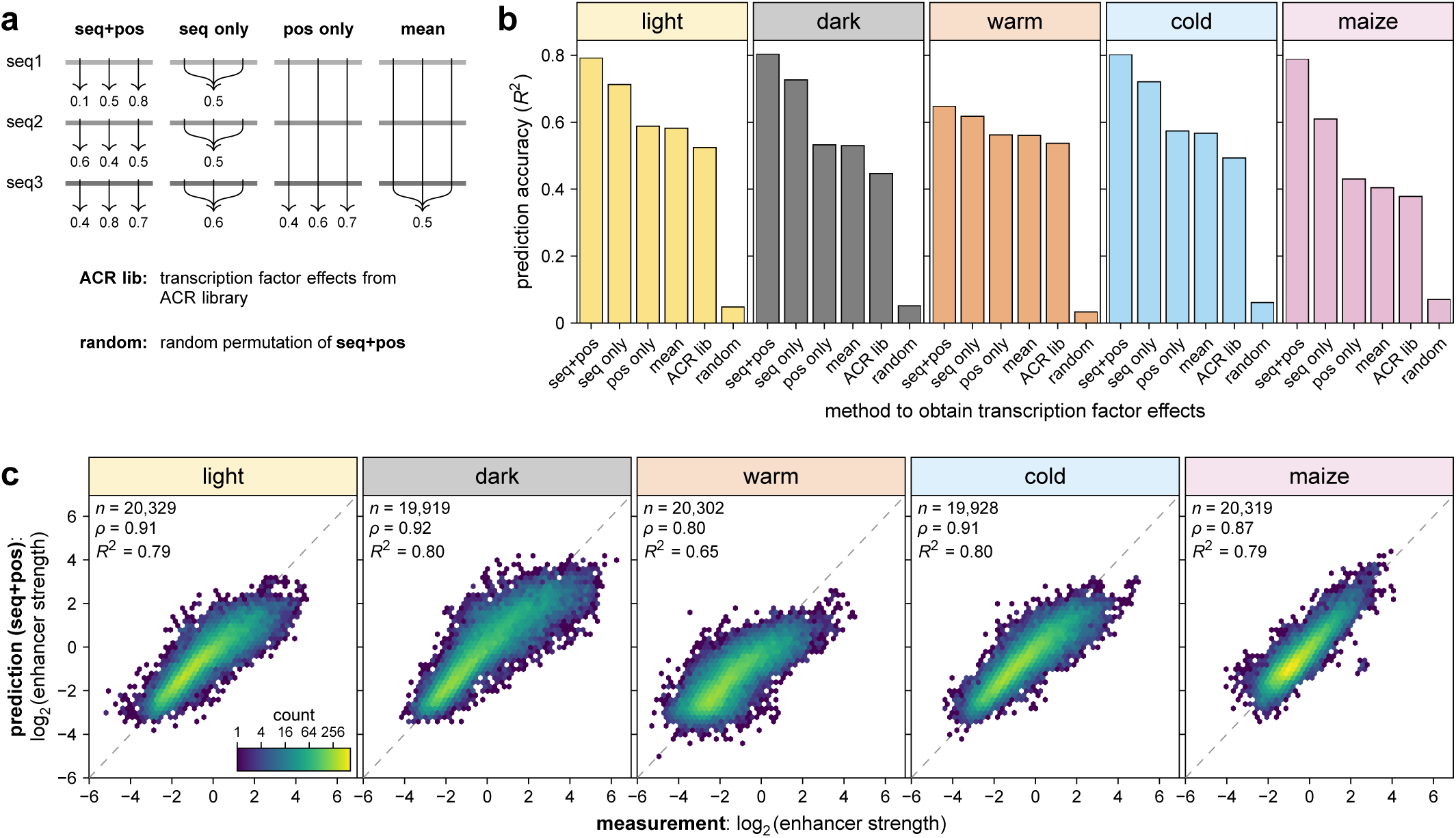
| The strength of synthetic enhancers can be predicted. **a**–**c**, The strength of synthetic enhancers with two or three transcription factor binding sites was predicted based on the strength of the random sequence without any binding sites and the effects observed for inserting individual binding sites. The transcription factor binding site effects were calculated individually for each random sequence and each position (seq+pos), as the mean across all positions for each sequence (seq only), as the mean across all sequences for each position (pos only), or as the mean across all sequences and positions (mean). Additionally, transcription factor binding site effects were obtained from the ACR library experiment (ACR lib; see Fig. 2a) or from a random permutation of the seq+pos method (**a**). For each method of obtaining transcription factor effects, the prediction accuracy was determined as the correlation between the predicted and experimentally determined synthetic enhancer strengths (**b**). Hexbin plots (as defined in Fig. 1) of this correlation are shown for the seq+pos method (**c**).

**Extended Data Fig. 8.**
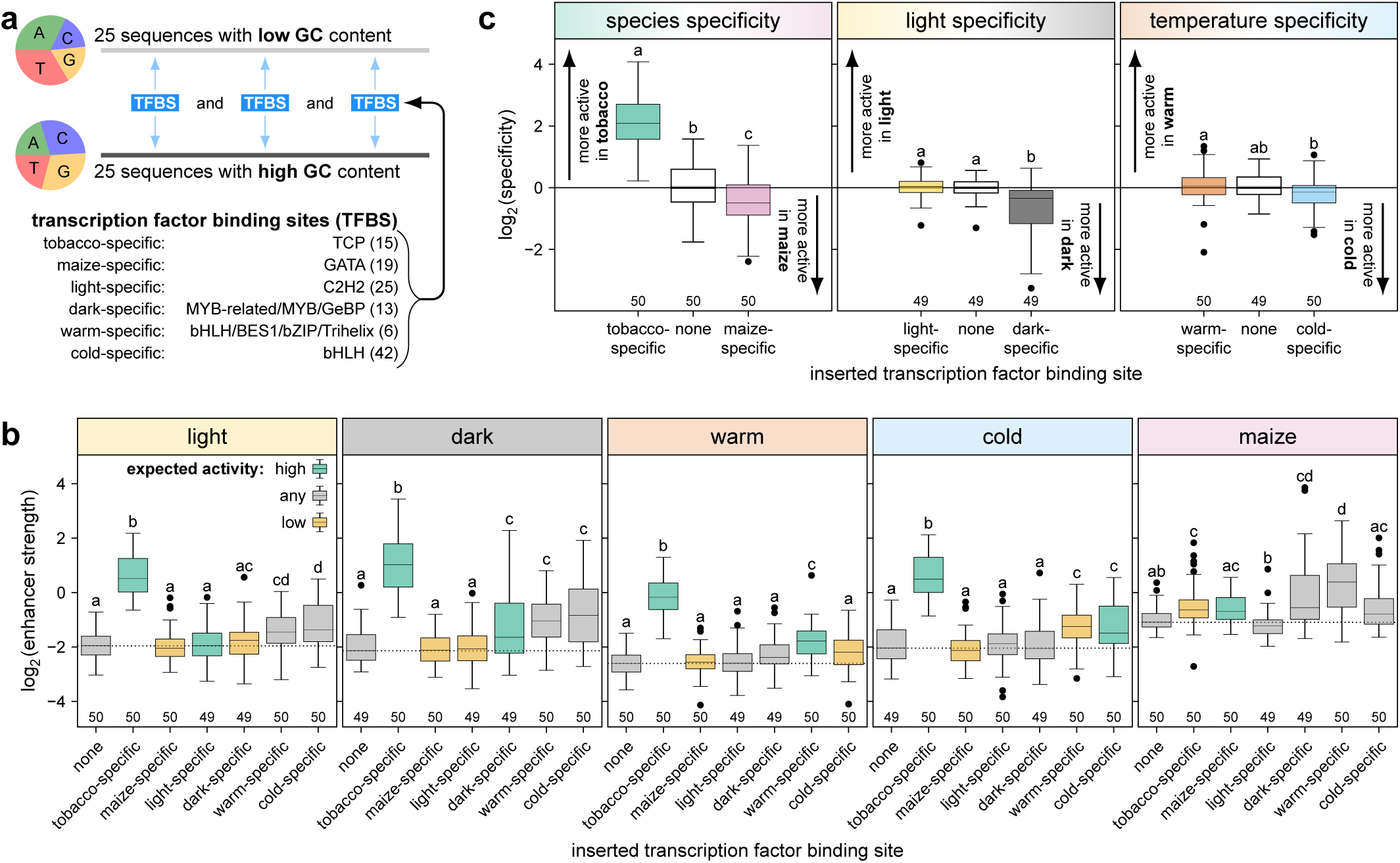
| Rationally designed synthetic enhancers show little condition specificity. **a**–**c**, Synthetic enhancers were designed by inserting three binding sites for transcription factors with species- or condition-specific effects into 25 random sequences each with a nucleotide composition similar to an average dicot (low GC) or monocot (high GC) ACR (**a**). The strength (**b**) and species or condition specificity (**c**) of these synthetic enhancers was determined by Plant STARR-seq in the indicated conditions. Species and condition specificity was calculated as the fold-change in enhancer strength between two species (tobacco [mean of light, dark, warm and cold] and maize) or conditions (light and dark for light specificity; warm and cold for temperature specificity). Box plots are as defined in Fig. 1 and letters above the box plots in indicate significance groups determined by post-hoc Tukey tests performed separately for each condition, species, or specificity. Exact *p* values are listed in Supplementary Data 4. In (**b**), box plots are colored according to the expected activity in the indicated condition.

**Extended Data Fig. 9.**
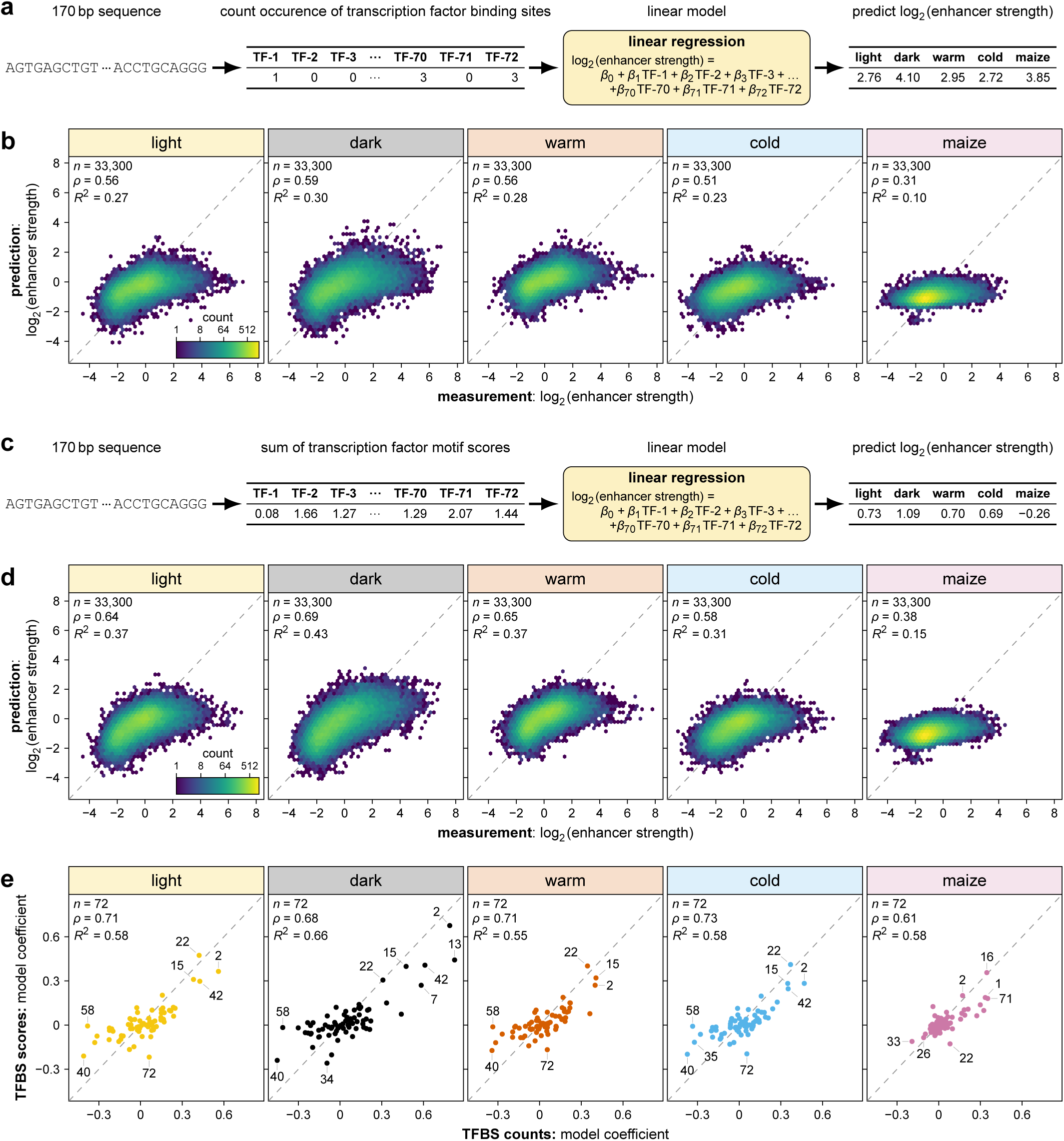
| Linear models based on transcription factor binding sites predict enhancer strength with low accuracy. **a**–**d**, Linear models based on the counts of putative transcription factor binding sites (**a**,**b**) or on the cumulative scores of binding motif matches within each sequence (**c**,**d**) were trained to predict enhancer strength. The predictions of the models were compared with the experimentally determined enhancer strengths in a held-out test set comprising 10% of the test sequences (**b**,**d**). **e**, Correlation between the coefficients assigned by the linear models in (**a**–**d**) to the 72 different transcription factor binding motifs. Selected motifs associated with increased or decreased enhancer strength are labeled. Hexbin plots in (**b**) and (**d**) are as defined in Fig. 1. In (**b**), (**c**), and (**e**), the dashed lines represents a *y* = *x* line. Pearson’s *R*^2^, Spearman’s *ρ*, and the number (*n*) of samples are indicated.

**Extended Data Fig. 10.**
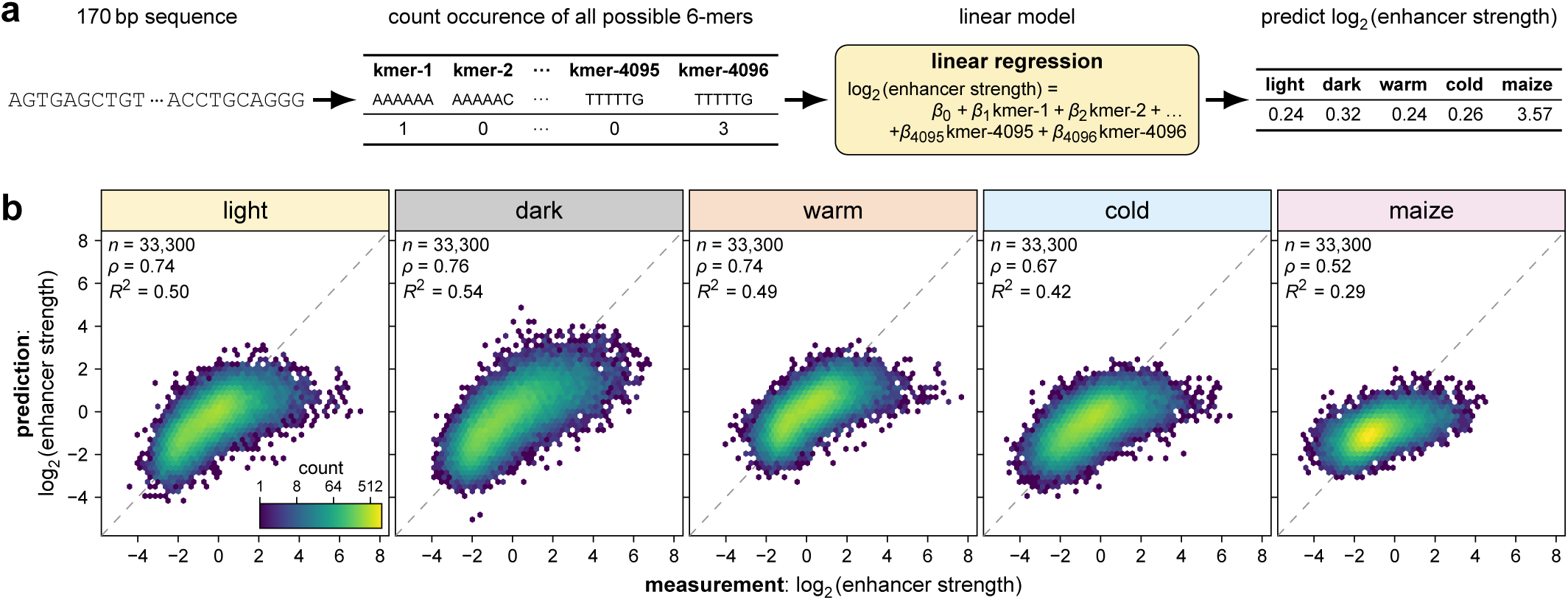
| A linear model based on 6-mer counts predicts enhancer strength with limited accuracy. **a**,**b**, A linear model based on the counts of all possible 6-mers in a given sequence was trained to predict enhancer strength (**a**). The model predictions were compared with the experimentally determined enhancer strengths in a held-out test set comprising 10% of the test sequences (**b**). Hexbin plots are as defined in Fig. 1.

## Notes

https://github.com/tobjores/plantGREP

### Competing Interest Statement

The authors have declared no competing interest.

## References

1. Marand A.P., Eveland A.L., Kaufmann K., Springer N.M. cis-Regulatory Elements in Plant Development, Adaptation, and Evolution. Annu. Rev. Plant Biol. 74, 111–137 (2023).

2. Weber B., Zicola J., Oka R., Stam M. Plant Enhancers: A Call for Discovery. Trends Plant Sci. 21, 974–987 (2016).

3. Andersson R., Sandelin A. Determinants of enhancer and promoter activities of regulatory elements. Nat. Rev. Genet. 21, 71–87 (2020).

4. Schmitz R.J., Grotewold E., Stam M. Cis-regulatory sequences in plants: Their importance, discovery, and future challenges. Plant Cell 34, 718–741 (2022).

5. Banerji J., Olson L., Schaffner W. A lymphocyte-specific cellular enhancer is located downstream of the joining region in immunoglobulin heavy chain genes. Cell 33, 729–740 (1983).

6. Banerji J., Rusconi S., Schaffner W. Expression of a β-globin gene is enhanced by remote SV40 DNA sequences. Cell 27, 299–308 (1981).

7. Chandrasekharappa S.C., Subramanian K.N. Effects of position and orientation of the 72-base-pair-repeat transcriptional enhancer on replication from the simian virus 40 core origin. J. Virol. 61, 2973–2980 (1987).

8. Gisselbrecht S.S. et al. Transcriptional Silencers in *Drosophila* Serve a Dual Role as Transcriptional Enhancers in Alternate Cellular Contexts. Mol. Cell 77, 324–337.e8 (2020).

9. Erhard K.F. Jr., Talbot J.-E.R.B., Deans N.C., McClish A.E., Hollick J.B. Nascent Transcription Affected by RNA Polymerase IV in Zea mays. Genetics 199, 1107–1125 (2015).

10. Hetzel J., Duttke S.H., Benner C., Chory J. Nascent RNA sequencing reveals distinct features in plant transcription. Proc. Natl. Acad. Sci. 113, 12316–12321 (2016).

11. Ricci W.A. et al. Widespread long-range cis-regulatory elements in the maize genome. Nat. Plants 5, 1237–1249 (2019).

12. Thieffry A. et al. Characterization of Arabidopsis thaliana promoter Bidirectionality and Antisense RNAs by Depletion of Nuclear RNA Decay Pathways. Plant Cell (2020).

13. Silver B.D., Willett C.G., Maher K.A., Wang D., Deal R.B. Differences in transcription initiation directionality underlie distinctions between plants and animals in chromatin modification patterns at genes and cis-regulatory elements. G3 GenesGenomesGenetics 14, jkae016 (2024).

14. McDonald B.R. et al. Enhancers associated with unstable RNAs are rare in plants. Nat. Plants 10, 1246–1257 (2024).

15. Paterson A.H., Queitsch C. Genome organization and botanical diversity. Plant Cell 36, 1186–1204 (2024).

16. Beernink B.M., Vogel J.P., Lei L. Enhancers in Plant Development, Adaptation and Evolution. Plant Cell Physiol. 66, 461–476 (2025).

17. Hendelman A. et al. Conserved pleiotropy of an ancient plant homeobox gene uncovered by cis-regulatory dissection. Cell 184, 1724–1739.e16 (2021).

18. Wang X. et al. Dissecting cis-regulatory control of quantitative trait variation in a plant stem cell circuit. Nat. Plants 7, 419–427 (2021).

19. Liu L. et al. Enhancing grain-yield-related traits by CRISPR–Cas9 promoter editing of maize CLE genes. Nat. Plants 7, 287–294 (2021).

20. Rodríguez-Leal D., Lemmon Z.H., Man J., Bartlett M.E., Lippman Z.B. Engineering Quantitative Trait Variation for Crop Improvement by Genome Editing. Cell 171, 470–480.e8 (2017).

21. Song X. et al. Targeting a gene regulatory element enhances rice grain yield by decoupling panicle number and size. Nat. Biotechnol. 40, 1403–1411 (2022).

22. Xiang H. et al. A molecular framework for lc controlled locule development of the floral meristem in tomato. Front. Plant Sci. 14, (2023).

23. Alexandre C.M. et al. Complex Relationships between Chromatin Accessibility, Sequence Divergence, and Gene Expression in Arabidopsis thaliana. Mol. Biol. Evol. 35, 837–854 (2018).

24. Marand A.P., Chen Z., Gallavotti A., Schmitz R.J. A *cis*-regulatory atlas in maize at single-cell resolution. Cell 184, 3041–3055.e21 (2021).

25. Parvathaneni R.K. et al. Regulatory signatures of drought response in stress resilient Sorghum bicolor. Preprint at bioRxiv https://www.biorxiv.org/content/10.1101/2020.08.07.240580v3 (2021).

26. Jores T. et al. Plant enhancers exhibit both cooperative and additive interactions among their functional elements. Plant Cell 36, 2570–2586 (2024).

27. Jores T. et al. Small DNA elements can act as both insulators and silencers in plants. Plant Cell 37, koaf084 (2025).

28. Jores T. et al. Identification of Plant Enhancers and Their Constituent Elements by STARR-seq in Tobacco Leaves. Plant Cell 32, 2120–2131 (2020).

29. Fluhr R., Kuhlemeier C., Nagy F., Chua N.-H. Organ-Specific and Light-Induced Expression of Plant Genes. Science 232, 1106–1112 (1986).

30. Simpson J., Schell J., Montagu M.V., Herrera-Estrella L. Light-inducible and tissue-specific pea lhcp gene expression involves an upstream element combining enhancer- and silencer-like properties. Nature 323, 551–554 (1986).

31. Nagy F., Boutry M., Hsu M.Y., Wong M., Chua N.H. The 5′-proximal region of the wheat Cab-1 gene contains a 268-bp enhancer-like sequence for phytochrome response. EMBO J. 6, 2537–2542 (1987).

32. Fang R.X., Nagy F., Sivasubramaniam S., Chua N.H. Multiple cis regulatory elements for maximal expression of the cauliflower mosaic virus 35S promoter in transgenic plants. Plant Cell 1, 141–150 (1989).

33. Sato S. et al. The tomato genome sequence provides insights into fleshy fruit evolution. Nature 485, 635–641 (2012).

34. Stelpflug S.C. et al. An Expanded Maize Gene Expression Atlas based on RNA Sequencing and its Use to Explore Root Development. Plant Genome 9, plantgenome2015.04.0025 (2016).

35. Wang B. et al. A comparative transcriptional landscape of maize and sorghum obtained by single-molecule sequencing. Genome Res. 28, 921–932 (2018).

36. Mergner J. et al. Mass-spectrometry-based draft of the Arabidopsis proteome. Nature 579, 409–414 (2020).

37. Moreno P. et al. Expression Atlas update: gene and protein expression in multiple species. Nucleic Acids Res. 50, D129–D140 (2022).

38. Yanai I. et al. Genome-wide midrange transcription profiles reveal expression level relationships in human tissue specification. Bioinformatics 21, 650–659 (2005).

39. Jores T. et al. Synthetic promoter designs enabled by a comprehensive analysis of plant core promoters. Nat. Plants 7, 842–855 (2021).

40. O’Malley R.C. et al. Cistrome and Epicistrome Features Shape the Regulatory DNA Landscape. Cell 165, 1280–1292 (2016).

41. Lehti-Shiu M.D., Panchy N., Wang P., Uygun S., Shiu S.-H. Diversity, expansion, and evolutionary novelty of plant DNA-binding transcription factor families. Biochim. Biophys. Acta BBA - Gene Regul. Mech. 1860, 3–20 (2017).

42. Zenker S. et al. Many transcription factor families have evolutionarily conserved binding motifs in plants. Plant Physiol. 198, kiaf205 (2025).

43. Huang G., Liu Z., Van Der Maaten L., Weinberger K.Q. Densely Connected Convolutional Networks. Preprint at https://ieeexplore.ieee.org/document/8099726 (2017).

44. Gorjifard S. et al. Arabidopsis and maize terminator strength is determined by GC content, polyadenylation motifs and cleavage probability. Nat. Commun. 15, 5868 (2024).

45. Deng K., Zhang Q., Hong Y., Yan J., Hu X. iCREPCP: A deep learning-based web server for identifying base-resolution cis-regulatory elements within plant core promoters. Plant Commun. 4, 100455 (2023).

46. Shrikumar A., Greenside P., Kundaje A. Learning Important Features Through Propagating Activation Differences. Preprint at arXiv http://arxiv.org/abs/1704.02685 (2019).

47. Shrikumar A. et al. Technical Note on Transcription Factor Motif Discovery from Importance Scores (TF-MoDISco) version 0.5.6.5. Preprint at arXiv http://arxiv.org/abs/1811.00416 (2020).

48. Clark R.M., Wagler T.N., Quijada P., Doebley J. A distant upstream enhancer at the maize domestication gene tb1 has pleiotropic effects on plant and inflorescent architecture. Nat. Genet. 38, 594–597 (2006).

49. Studer A., Zhao Q., Ross-Ibarra J., Doebley J. Identification of a functional transposon insertion in the maize domestication gene tb1. Nat. Genet. 43, 1160–1163 (2011).

50. Sullivan A.M. et al. Mapping and Dynamics of Regulatory DNA and Transcription Factor Networks in A. thaliana. Cell Rep. 8, 2015–2030 (2014).

51. de Boer C.G. et al. Deciphering eukaryotic gene-regulatory logic with 100 million random promoters. Nat. Biotechnol. 38, 56–65 (2020).

52. Kulkarni M.M., Arnosti D.N. Information display by transcriptional enhancers. Development 130, 6569–6575 (2003).

53. Thanos D., Maniatis T. Virus induction of human IFNβ gene expression requires the assembly of an enhanceosome. Cell 83, 1091–1100 (1995).

54. Panne D. The enhanceosome. Curr. Opin. Struct. Biol. 18, 236–242 (2008).

55. Simicevic J., Deplancke B. Transcription factor proteomics—Tools, applications, and challenges. PROTEOMICS 17, 1600317 (2017).

56. Dorrity M.W. et al. The regulatory landscape of Arabidopsis thaliana roots at single-cell resolution. Nat. Commun. 12, 3334 (2021).

57. Zhang T.-Q., Chen Y., Liu Y., Lin W.-H., Wang J.-W. Single-cell transcriptome atlas and chromatin accessibility landscape reveal differentiation trajectories in the rice root. Nat. Commun. 12, 2053 (2021).

58. Demesa-Arevalo E. et al. Imputation integrates single-cell and spatial gene expression data to resolve transcriptional networks in barley shoot meristem development. Preprint at bioRxiv https://www.biorxiv.org/content/10.1101/2025.05.09.653223v1 (2025).

59. Stebbins G.L. A brief summary of my ideas on evolution. Am. J. Bot. 86, 1207–1208 (1999).

60. Sullivan A.M. et al. Mapping and Dynamics of Regulatory DNA in Maturing Arabidopsis thaliana Siliques. Front. Plant Sci. 10, (2019).

61. Maurano M.T. et al. Systematic Localization of Common Disease-Associated Variation in Regulatory DNA. Science 337, 1190–1195 (2012).

62. Boyle E.A., Li Y.I., Pritchard J.K. An Expanded View of Complex Traits: From Polygenic to Omnigenic. Cell 169, 1177–1186 (2017).

63. Sinnott-Armstrong N. et al. Understanding genetic variants in context. eLife 13, e88231 (2024).

64. Sinnott-Armstrong N., Naqvi S., Rivas M., Pritchard J.K. GWAS of three molecular traits highlights core genes and pathways alongside a highly polygenic background. eLife 10, e58615 (2021).

65. Connally N.J. et al. The missing link between genetic association and regulatory function. eLife 11, e74970 (2022).

66. Claussnitzer M. et al. FTO Obesity Variant Circuitry and Adipocyte Browning in Humans. N. Engl. J. Med. 373, 895–907 (2015).

67. Stergachis A.B., Debo B.M., Haugen E., Churchman L.S., Stamatoyannopoulos J.A. Single-molecule regulatory architectures captured by chromatin fiber sequencing. Science 368, 1449–1454 (2020).

68. Jores T., Hamm M., Cuperus J.T., Queitsch C. Frontiers and techniques in plant gene regulation. Curr. Opin. Plant Biol. 75, 102403 (2023).

69. Bubb K.L. et al. The regulatory potential of transposable elements in maize. Nat. Plants 11, 1181–1192 (2025).

70. Cheng C.-Y. et al. Araport11: a complete reannotation of the Arabidopsis thaliana reference genome. Plant J. 89, 789–804 (2017).

71. Jiao Y. et al. Improved maize reference genome with single-molecule technologies. Nature 546, 524–527 (2017).

72. McCormick R.F. et al. The Sorghum bicolor reference genome: improved assembly, gene annotations, a transcriptome atlas, and signatures of genome organization. Plant J. 93, 338–354 (2018).

73. Alonge M. et al. Automated assembly scaffolding using RagTag elevates a new tomato system for high-throughput genome editing. Genome Biol. 23, 258 (2022).

74. Mejía-Guerra M.K. et al. Core Promoter Plasticity Between Maize Tissues and Genotypes Contrasts with Predominance of Sharp Transcription Initiation Sites. Plant Cell 27, 3309–3320 (2015).

75. Engler C., Kandzia R., Marillonnet S. A One Pot, One Step, Precision Cloning Method with High Throughput Capability. PLOS ONE 3, e3647 (2008).

76. Tonnies J., Mueth N.A., Gorjifard S., Chu J., Queitsch C. Scalable Transfection of Maize Mesophyll Protoplasts. JoVE J. Vis. Exp. e64991 (2023).

77. Clough S.J., Bent A.F. Floral dip: a simplified method for Agrobacterium -mediated transformation of Arabidopsis thaliana. Plant J. 16, 735–743 (1998).

78. Masella A.P., Bartram A.K., Truszkowski J.M., Brown D.G., Neufeld J.D. PANDAseq: paired-end assembler for illumina sequences. BMC Bioinformatics 13, 1–7 (2012).

79. Quinlan A.R., Hall I.M. BEDTools: a flexible suite of utilities for comparing genomic features. Bioinformatics 26, 841–842 (2010).

80. Neph S. et al. BEDOPS: high-performance genomic feature operations. Bioinformatics 28, 1919–1920 (2012).

81. Hammal F., de Langen P., Bergon A., Lopez F., Ballester B. ReMap 2022: a database of Human, Mouse, Drosophila and Arabidopsis regulatory regions from an integrative analysis of DNA-binding sequencing experiments. Nucleic Acids Res. 50, D316–D325 (2022).

82. Liu Y. et al. Genome-wide mapping of DNase I hypersensitive sites reveals chromatin accessibility changes in Arabidopsis euchromatin and heterochromatin regions under extended darkness. Sci. Rep. 7, 4093 (2017).

83. Raudvere U. et al. g:Profiler: a web server for functional enrichment analysis and conversions of gene lists (2019 update). Nucleic Acids Res. 47, W191–W198 (2019).

84. Tremblay B.J.-M. universalmotif: An R package for biological motif analysis. J. Open Source Softw. 9, 7012 (2024).

85. Pedregosa F. et al. Scikit-learn: Machine Learning in Python. J. Mach. Learn. Res. 12, 2825–2830 (2011).

86. Klie A. et al. Predictive analyses of regulatory sequences with EUGENe. Nat. Comput. Sci. 3, 946–956 (2023).

87. Paszke A. et al. PyTorch: An Imperative Style, High-Performance Deep Learning Library. Preprint at *Curran Associates, Inc*. https://proceedings.neurips.cc/paper_files/paper/2019/hash/bdbca288fee7f92f2bfa9f7012727740-Abstract.html (2019).

88. Kingma D.P., Ba J. Adam: A Method for Stochastic Optimization. Preprint at arXiv http://arxiv.org/abs/1412.6980 (2017).

89. Kokhlikyan N. et al. Captum: A unified and generic model interpretability library for PyTorch. Preprint at arXiv http://arxiv.org/abs/2009.07896 (2020).

