## Supplementary Figures for "Massively parallel characterization and deep learning of enhancers in plant genomes"

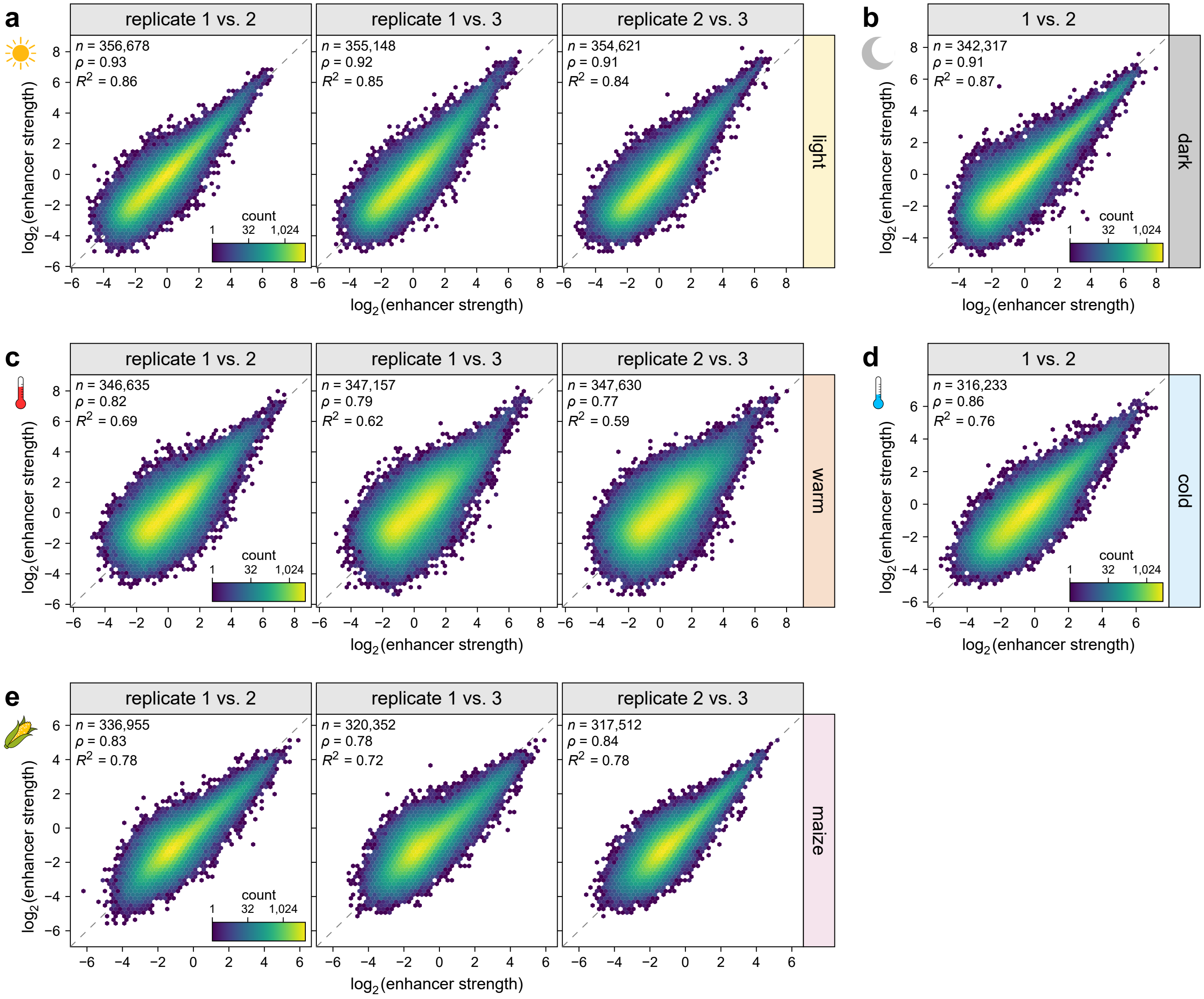


**Supplementary Fig. 1 | Plant STARR-seq is reproducible. a**–**e**, Hexbin plots (color represents the count of points in each hexagon) of the correlation between enhancer strength in independent Plant STARR-seq replicate experiments in the indicated conditions. The dashed lines represents a *y* = *x* line. Pearson's $R^2$, Spearman's $\rho$, and the number ($n$) of samples are indicated.

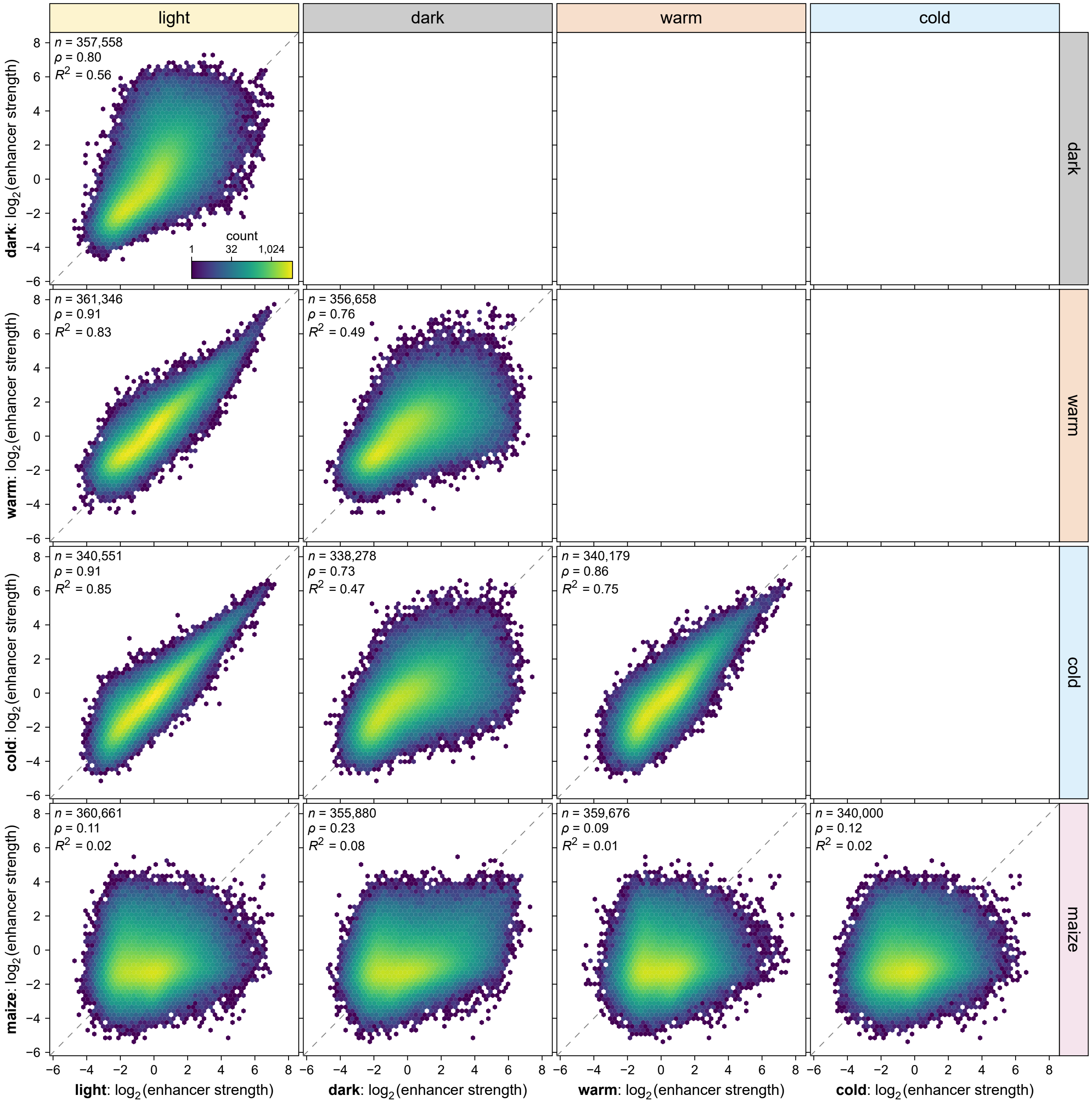


**Supplementary Fig. 2 | Plant STARR-seq detects species- and condition-specific differences in enhancer strength.** Hexbin plots (as defined in Supplementary Fig. 1) of the correlation between enhancer strength in different conditions.

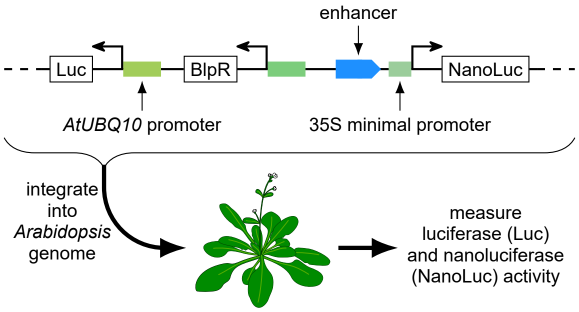


**Supplementary Fig. 3 | A dual-luciferase assay to measure enhancer strength in stable *Arabidopsis* lines.** Transgenic Arabidopsis lines were generated with T-DNAs harboring a constitutively expressed luciferase (Luc) gene and a nanoluciferase (NanoLuc) gene under control of a 35S minimal promoter coupled to a candidate enhancer. Luciferase and nanoluciferase activity was measured in the T2 generation.

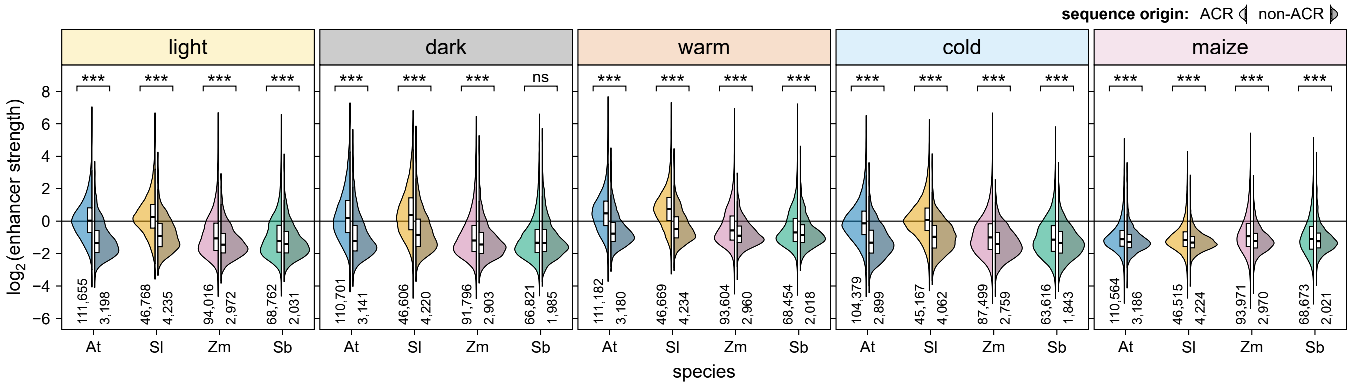


**Supplementary Fig. 4 | Accessible chromatin regions are enriched for active enhancers.** Enhancer strength of sequences derived from accessible chromatin regions (ACR, left) and non-ACR sequences (right) randomly sampled from inaccessible regions in the genomes of the indicated species. At, *Arabidopsis*; Sl, tomato; Zm, maize; Sb, sorghum. Violin plots represent the kernel density distribution and the box plots inside represent the median (center line), upper and lower quartiles (box limits), and 1.5× interquartile range (whiskers) for all corresponding sequences. Significant differences between ACR and non-ACR sequences were determined by two-sided Wilcoxon rank-sum tests. The obtained *p* values were Bonferroni-adjusted separately for each condition and species and are indicated: *, $p \leq 0.05$; **, $p \leq 0.01$; ***, $p \leq 0.001$; ns, not significant. Exact *p* values are listed in Supplementary Data 4.

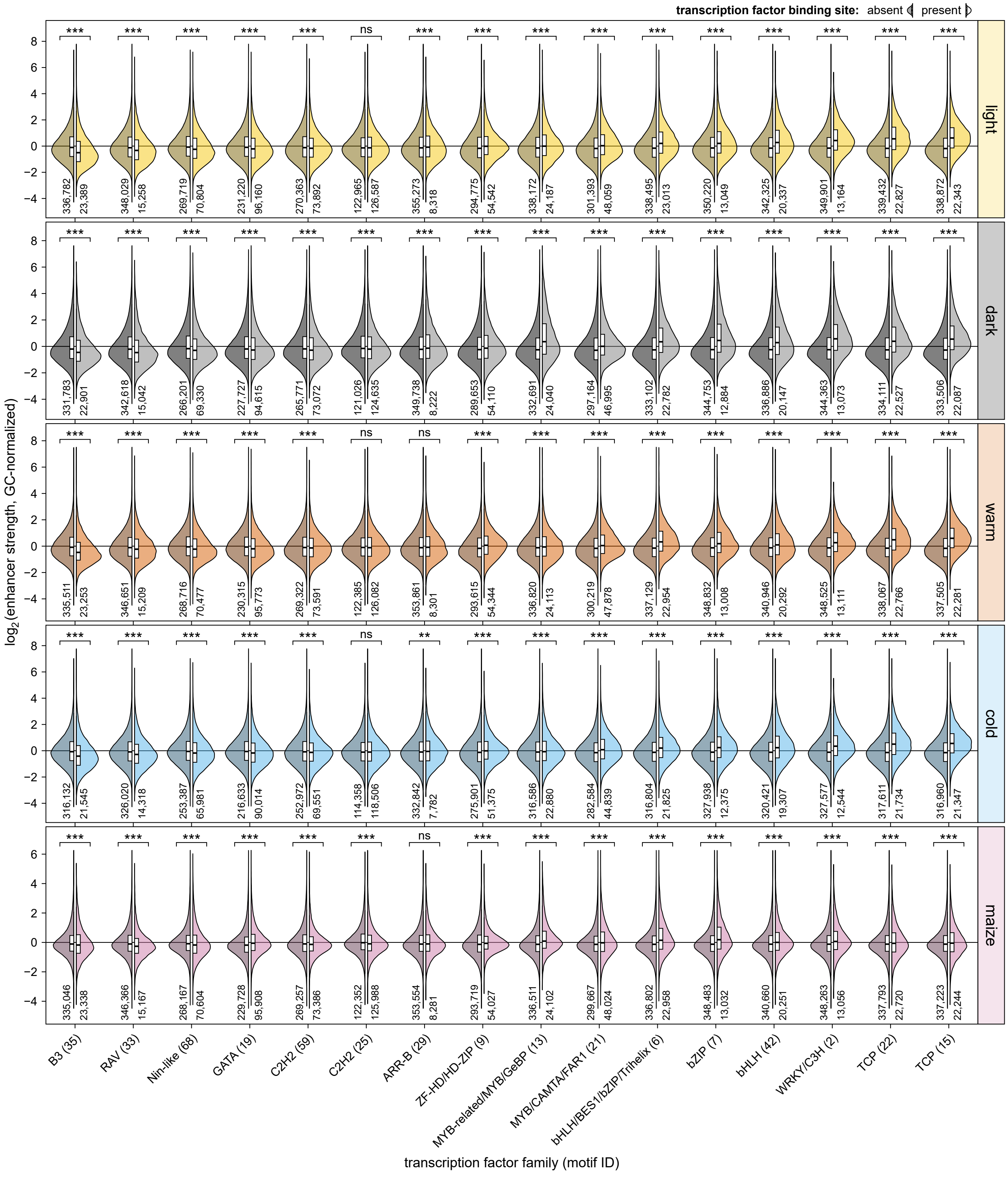


**Supplementary Fig. 5 | The presence of transcription factor binding sites affects enhancer strength.** Putative transcription factor binding sites were identified by scanning the test sequences with known binding motifs for transcription factors of the indicated families. The enhancer strength of test sequences without or with one putative binding site is shown in violin plots as defined in Supplementary Fig. 4. Significant differences between sequences with and without a binding site were determined by two-sided Wilcoxon rank-sum tests. The obtained $p$ values were Bonferroni-adjusted separately for each condition and species and are indicated: *, $p \leq 0.05$; **, $p \leq 0.01$; ***, $p \leq 0.001$; ns, not significant. Exact $p$ values are listed in Supplementary Data 4.

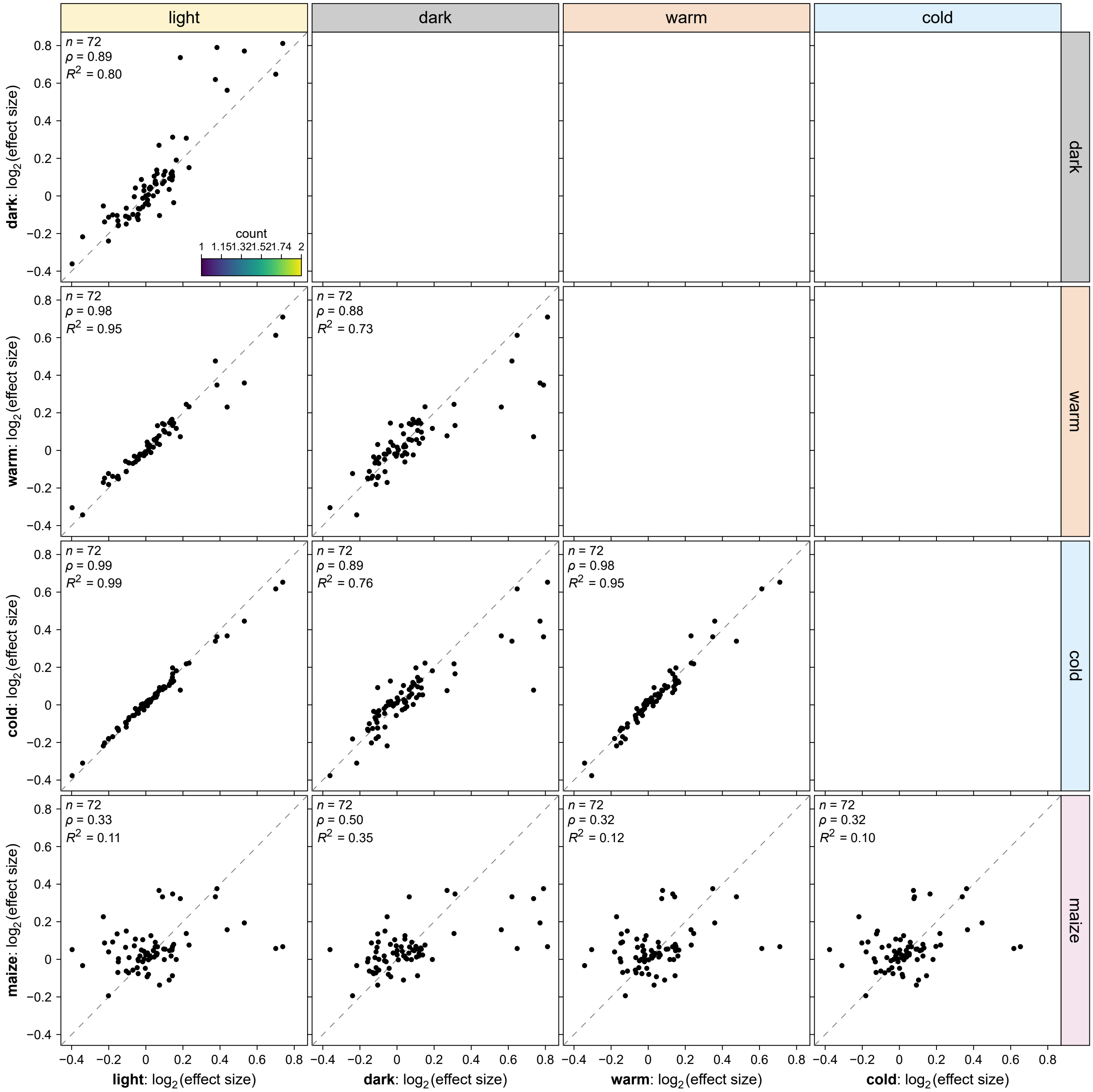


**Supplementary Fig. 6 | Transcription factor binding sites show species- and condition-specific effects on enhancer strength.** Correlation between the effects of transcription factor binding sites (as determined in Fig. 2) in different conditions. Each dot represents a different transcription factor binding motif. Pearson's $R^2$, Spearman's $\rho$, and the number ($n$) of samples are indicated.

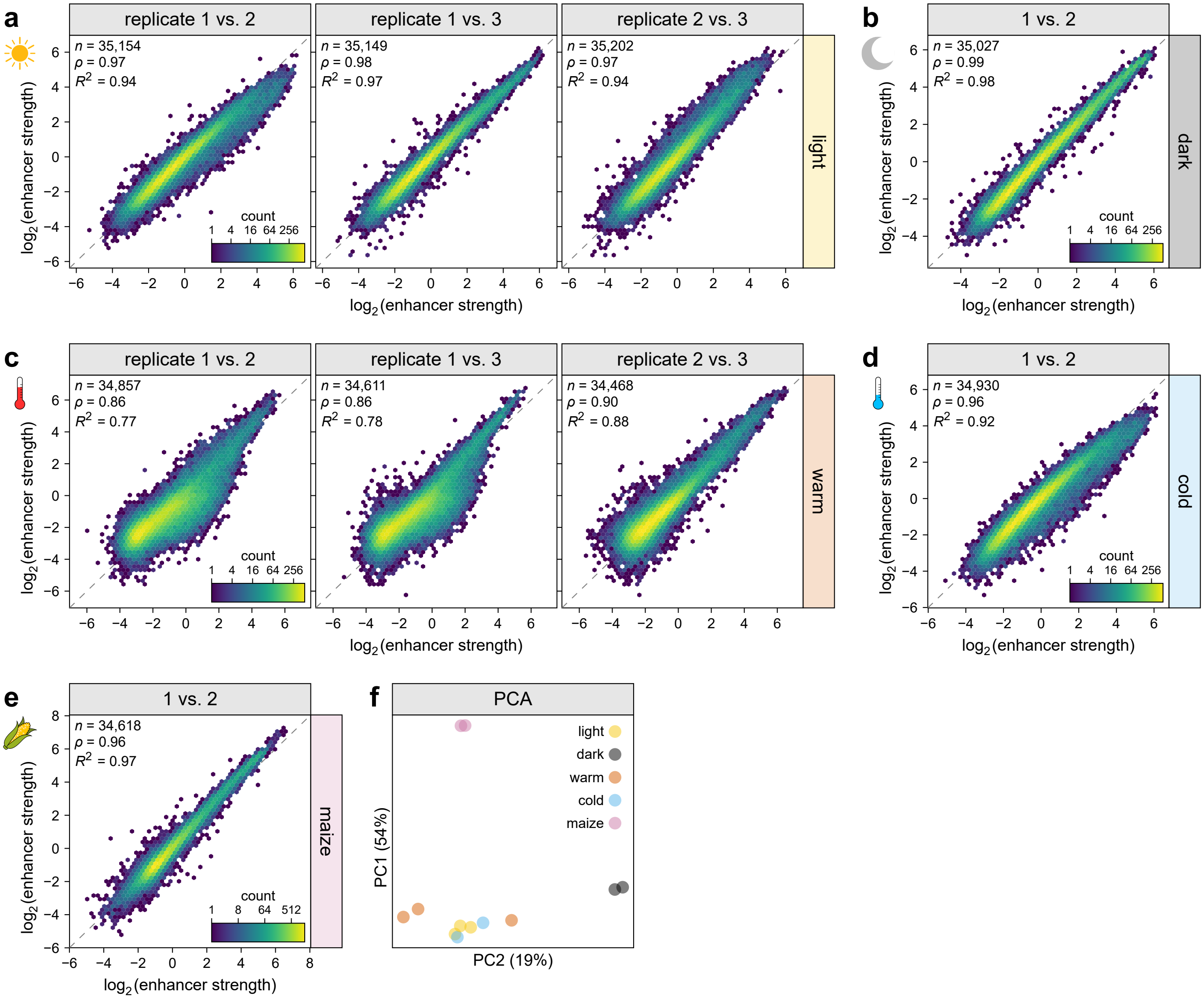


**Supplementary Fig. 7 | The validation library yielded reproducible results. a**–**e**, Hexbin plots (as defined in Supplementary Fig. 1) of the correlation between enhancer strength in independent Plant STARR-seq replicate experiments with the validation library in the indicated conditions. **f**, Principal component analysis of enhancer strength determined in Plant STARR-seq replicate experiments with the validation library.

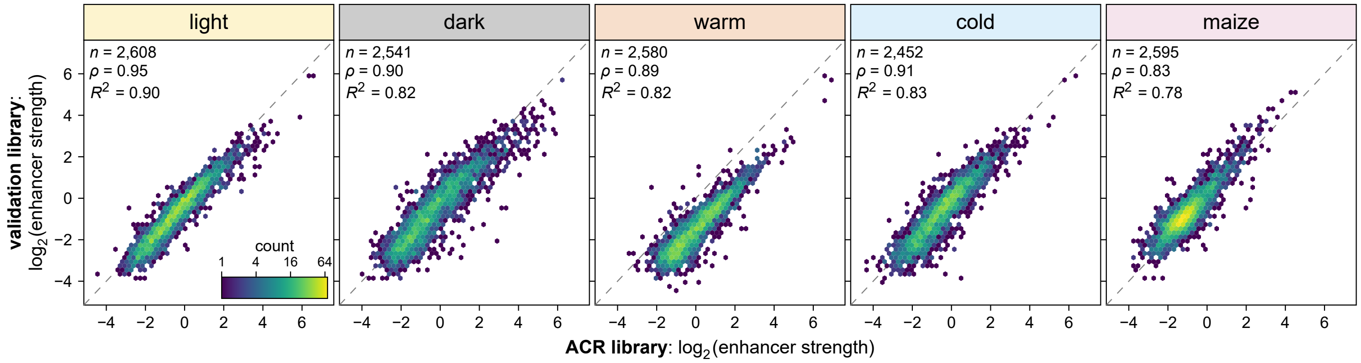


**Supplementary Fig. 8 | The validation library results reproduce the results obtained with the ACR library.** Hexbin plots (as defined in Supplementary Fig. 1) of the correlation between the strength of enhancers included in the ACR library and the validation library.

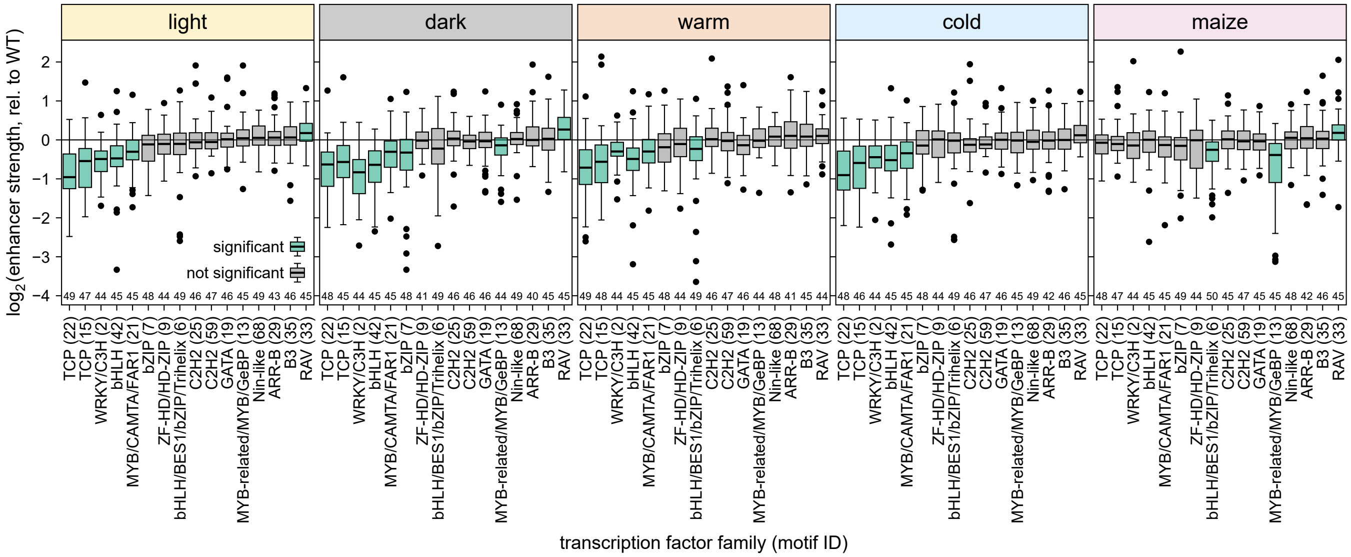


**Supplementary Fig. 9 | Removing transcription factor binding sites affects enhancer strength.** Relative enhancer strength of sequences with mutations destroying a single strong ($p$ value ≤ 0.0001) binding site for transcription factors of the indicated family. For each sequence, the enhancer strength was normalized to the strength of the corresponding wild-type sequence. Box plots represent the median (center line), upper and lower quartiles (box limits), 1.5× interquartile range (whiskers), and outliers (points) for all corresponding sequences. Numbers at the bottom of the plots indicate the number of samples in each group. Significant differences from a null distribution were determined using one-sample Wilcoxon rank-sum tests. Box plots for significant groups (Bonferroni-adjusted $p$ value ≤ 0.05) are shown in green. Non-significant groups are represented by gray box plots. Exact $p$ values are listed in Supplementary Data 4.

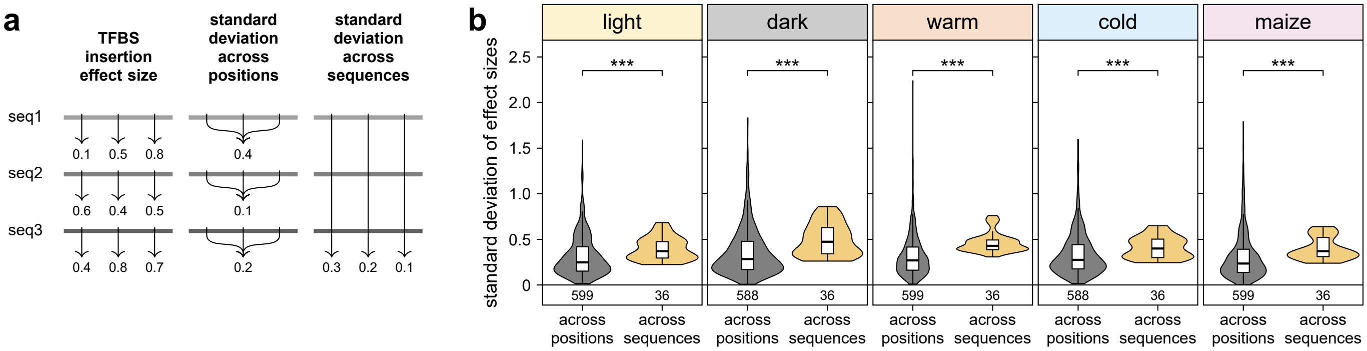


**Supplementary Fig. 10 | Sequence context has a stronger impact on enhancer strength than position of a transcription factor binding site.** A transcription factor binding site (TFBS) was inserted into 50 random sequences in one of three different positions, and the effect size of each binding site was determined as the fold-change in enhancer strength between the sequence with and without this binding site. For each binding site, the standard deviation of its effects sizes was calculated across the three positions within each sequence (across positions) or across all random sequences for each position (across sequences). Violin plots are as defined in Supplementary Fig. 4. Significant differences were determined by two-sided Wilcoxon rank-sum tests: *, $p \leq 0.05$; **, $p \leq 0.01$; ***, $p \leq 0.001$; ns, not significant. Exact $p$ values are listed in Supplementary Data 4.

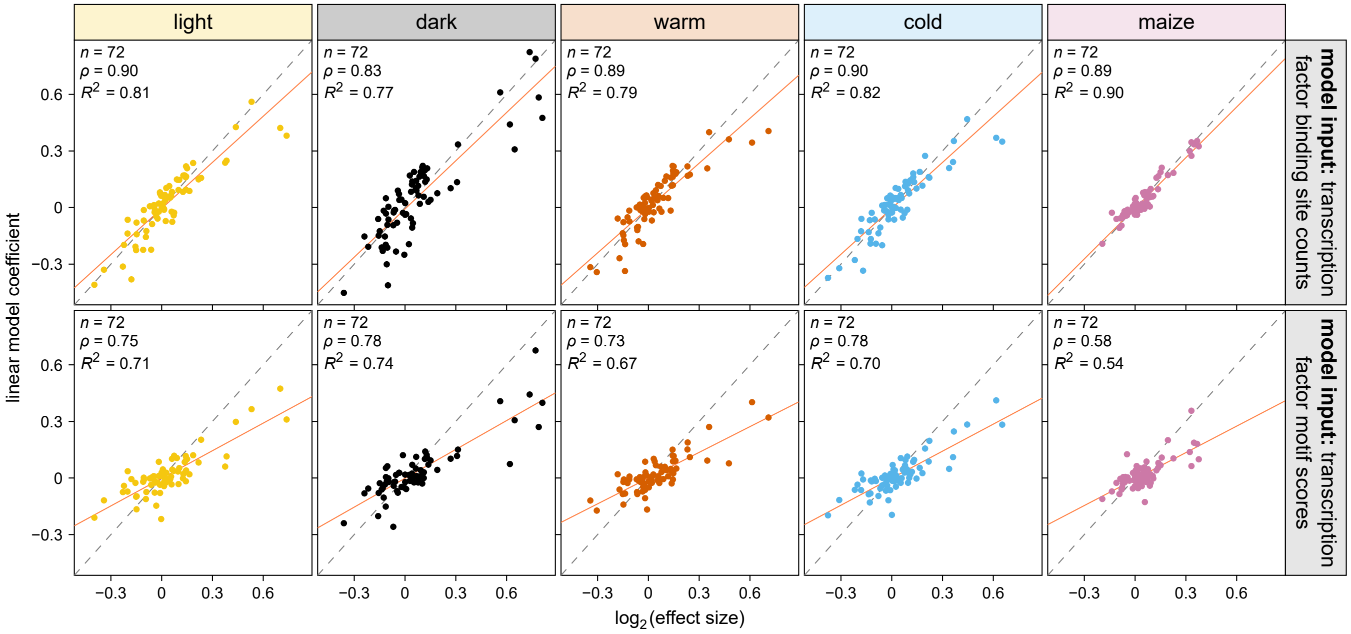


**Supplementary Fig. 11 | Linear model coefficients are correlated with the effects of transcription factor binding sites in the ACR library.** Correlation between the coefficients of the linear models from Extended Data Fig. 9 and the transcription factor binding site effects in the ACR library (see Fig. 2a). The solid and dashed lines represents a $y = x$ line and a linear regression line, respectively. Pearson's $R^2$, Spearman's $\rho$, and the number ($n$) of samples are indicated.

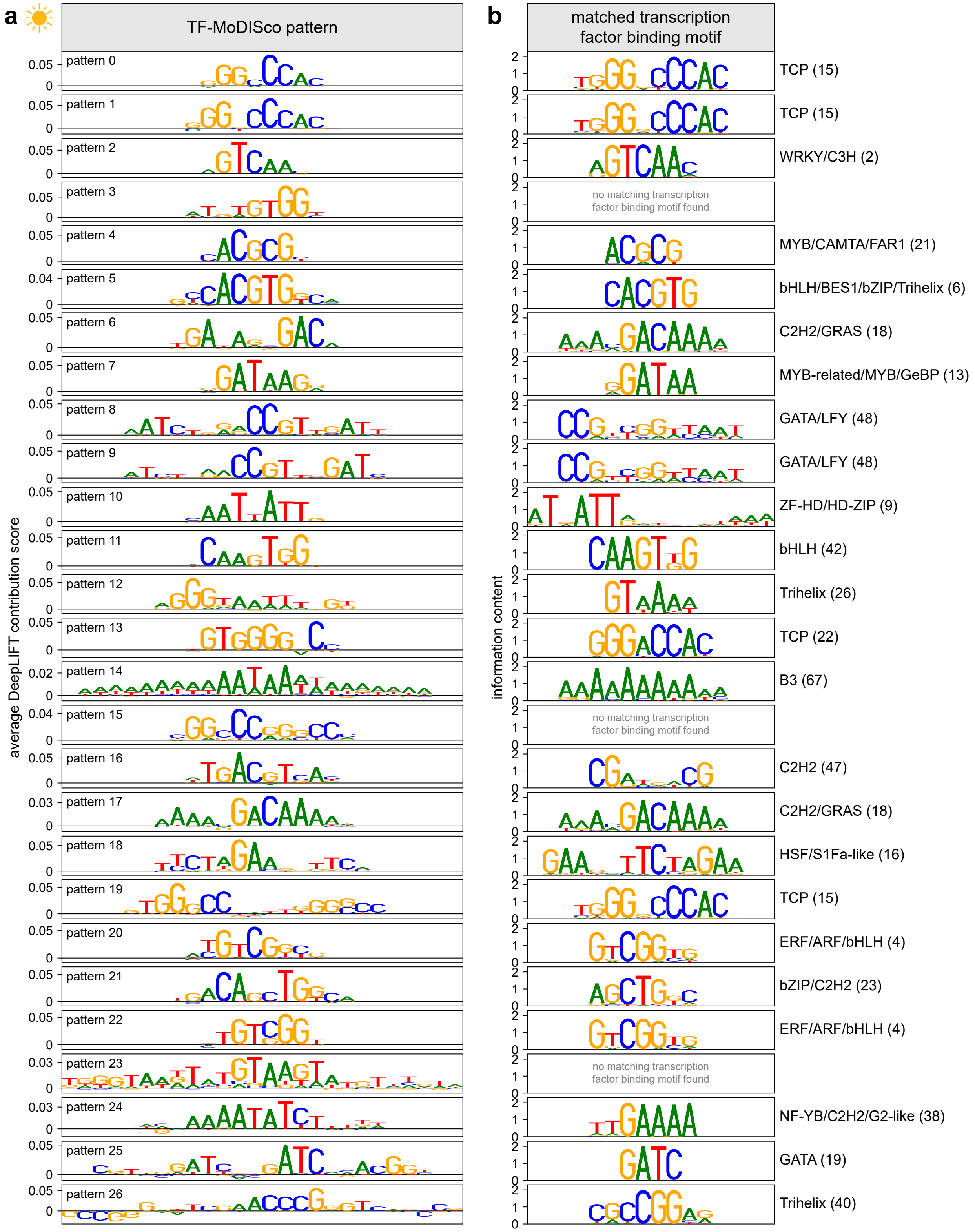


**Supplementary Fig. 12 | TF-MoDISco patterns match known transcription factor binding motifs. a**, DNA Patterns identified by TF-MoDISco that positively contribute to plantGREP predictions of enhancer strength in tobacco leaves in the light. **b**, Transcription factor binding motifs that match the patterns in (**a**).

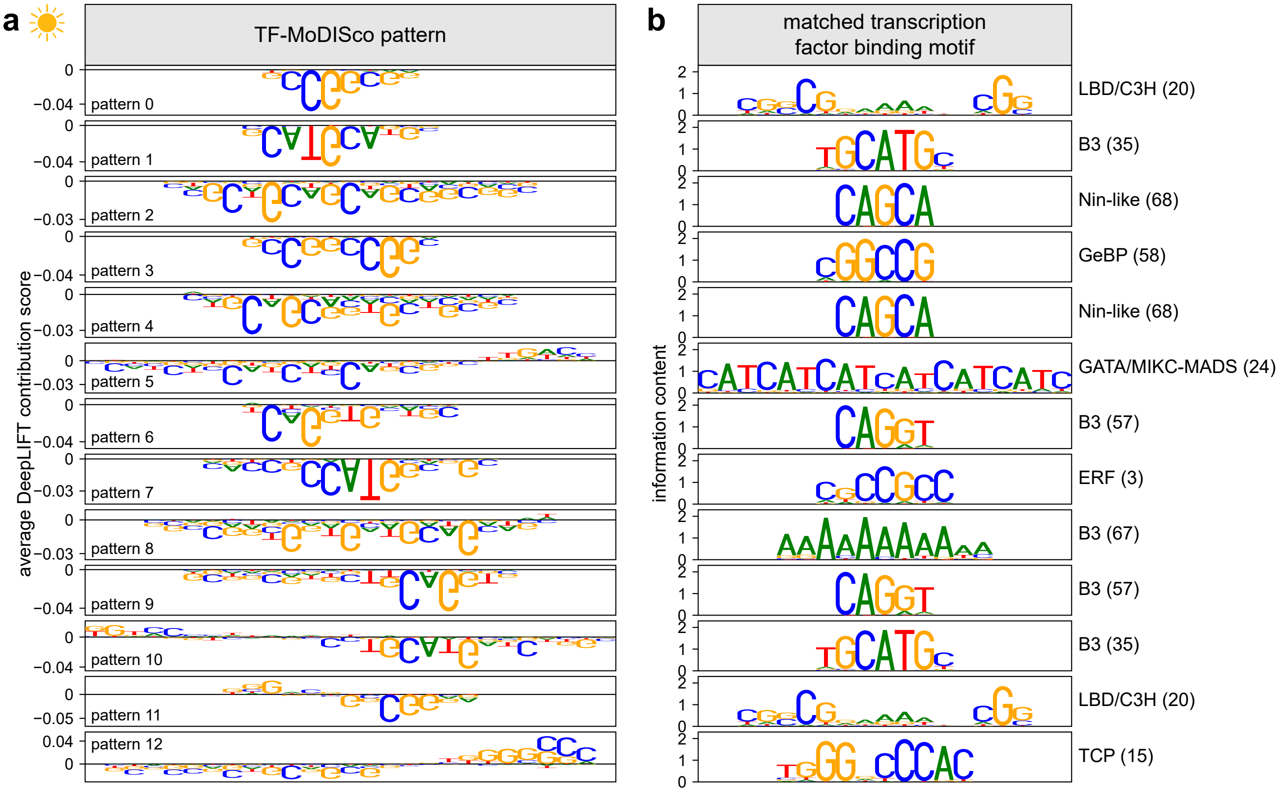


**Supplementary Fig. 13 | TF-MoDISco patterns match known transcription factor binding motifs.** **a**, DNA Patterns identified by TF-MoDISco that negatively contribute to plantGREP predictions of enhancer strength in tobacco leaves in the light. **b**, Transcription factor binding motifs that match the patterns in (**a**).

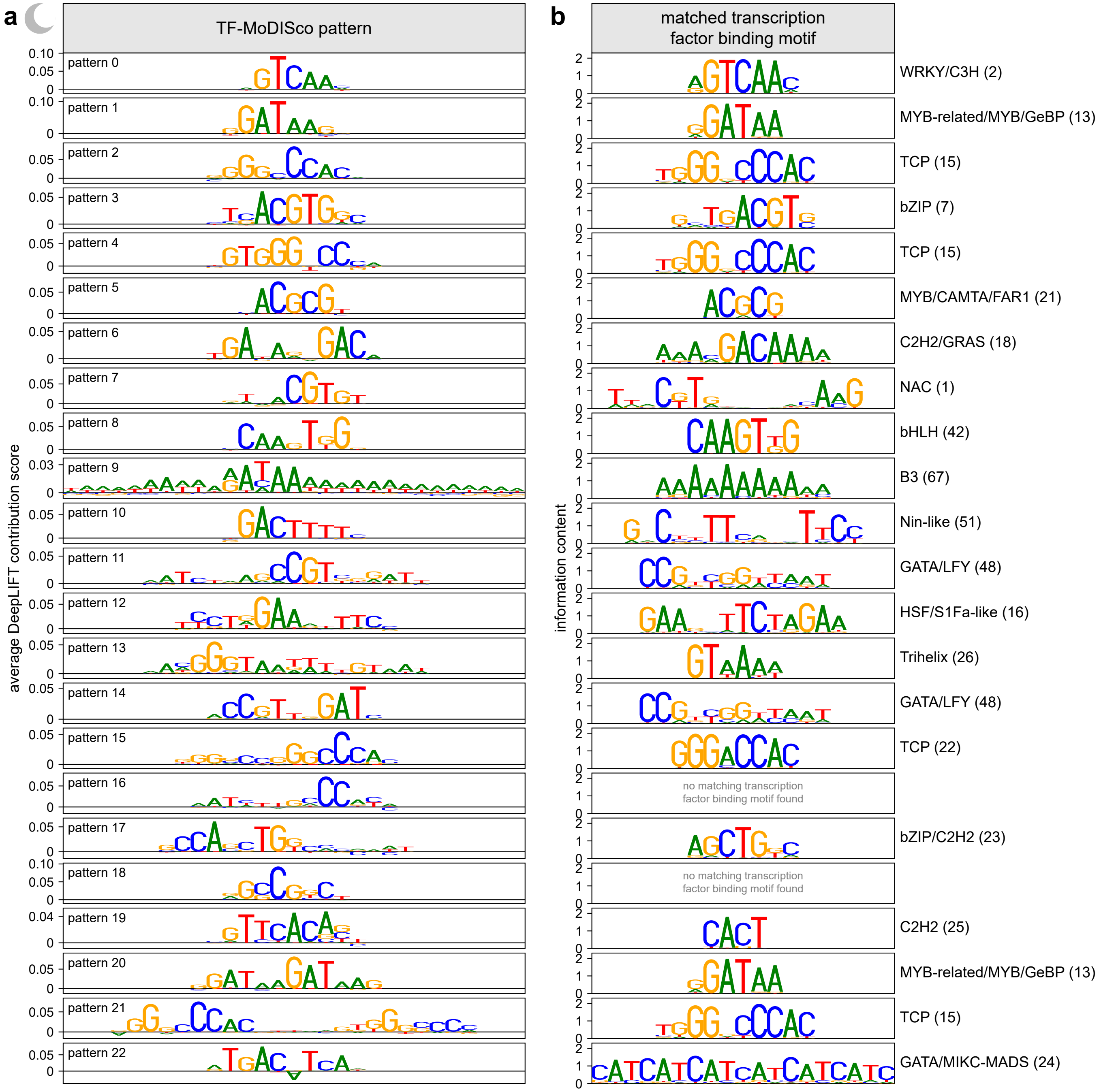


**Supplementary Fig. 14 | TF-MoDISco patterns match known transcription factor binding motifs.** **a**, DNA Patterns identified by TF-MoDISco that positively contribute to plantGREP predictions of enhancer strength in tobacco leaves in the dark. **b**, Transcription factor binding motifs that match the patterns in (**a**).

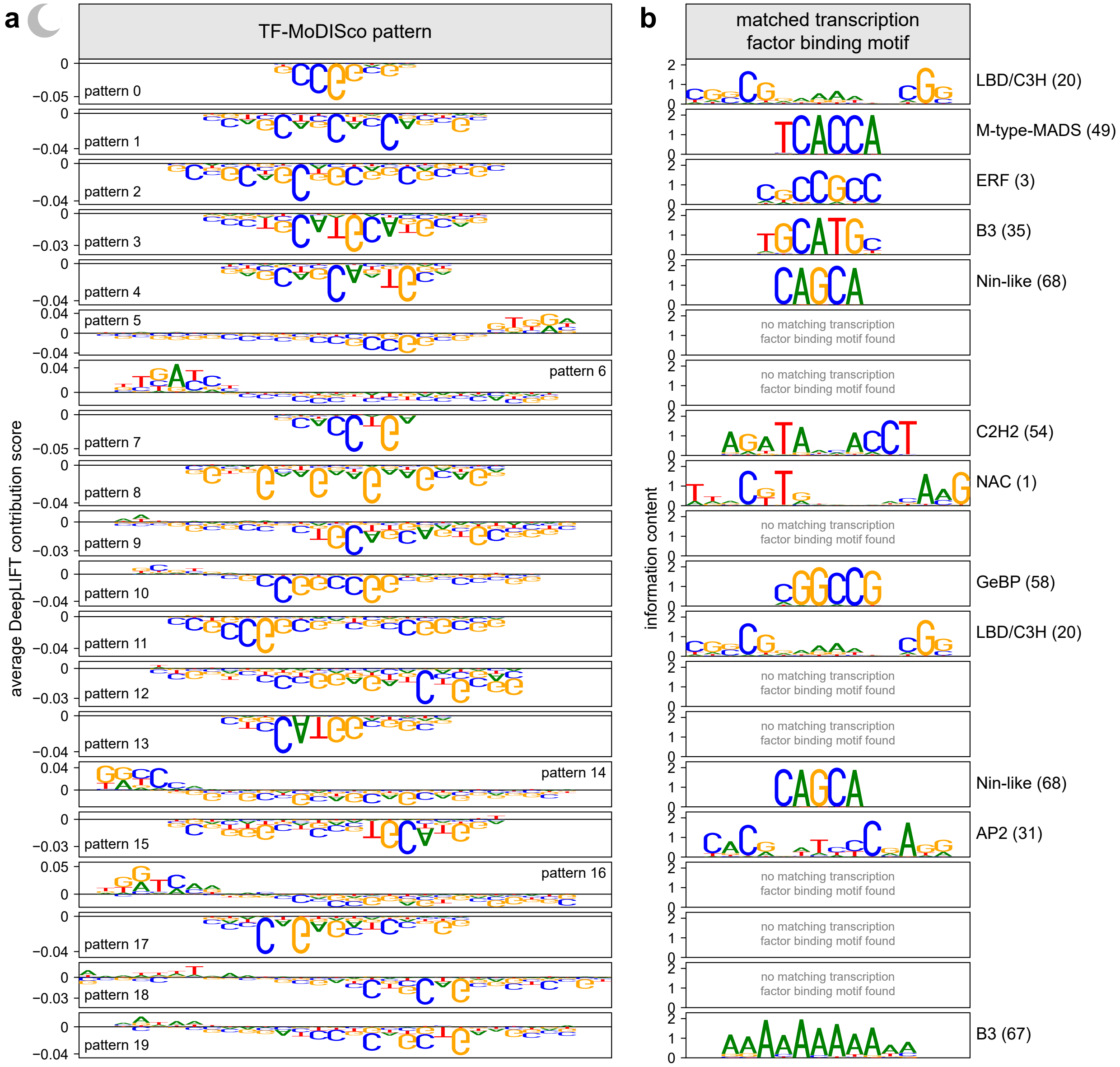


**Supplementary Fig. 15 | TF-MoDISco patterns match known transcription factor binding motifs. a**, DNA Patterns identified by TF-MoDISco that negatively contribute to plantGREP predictions of enhancer strength in tobacco leaves in the dark. **b**, Transcription factor binding motifs that match the patterns in (**a**).

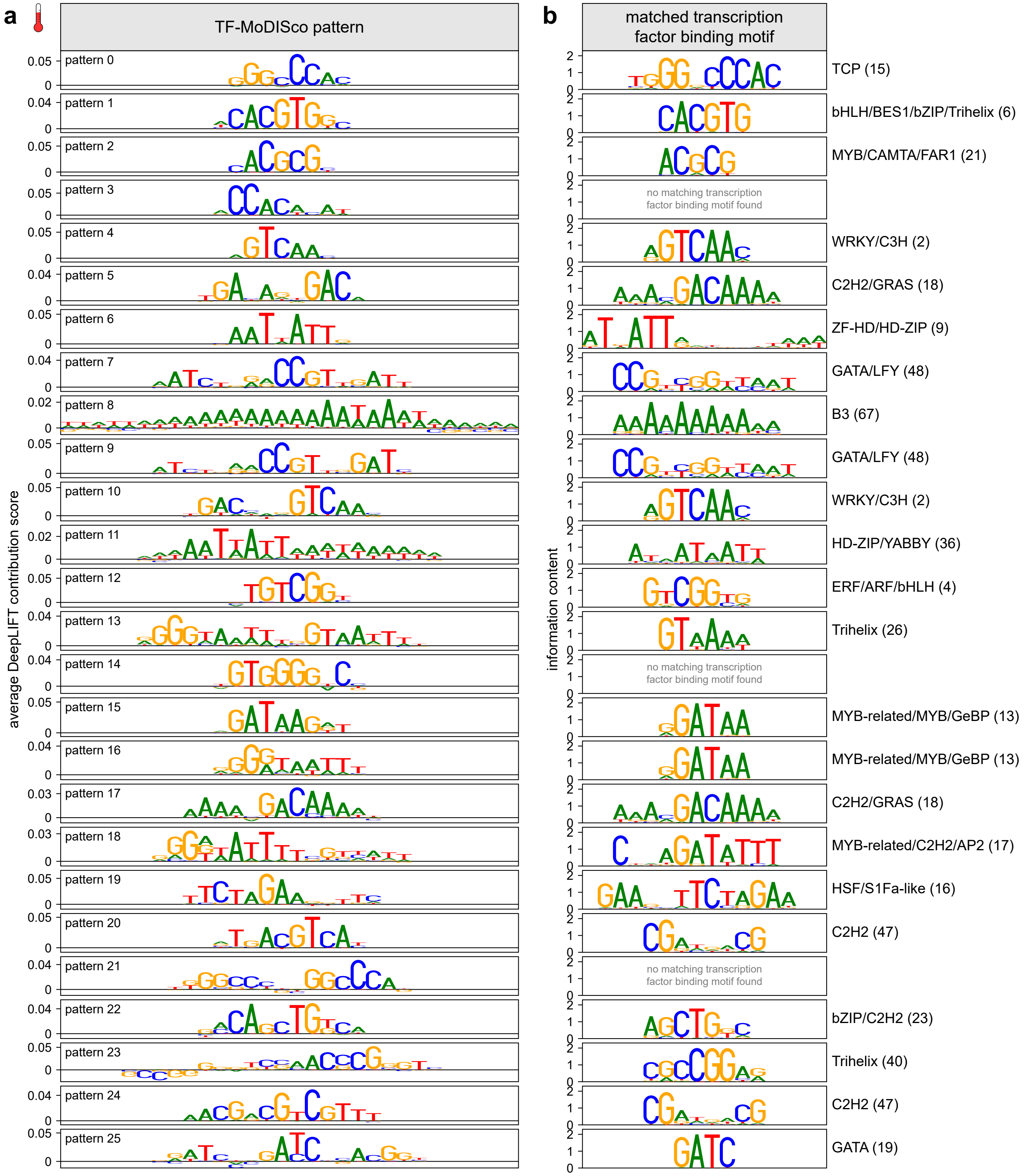


**Supplementary Fig. 16 | TF-MoDISco patterns match known transcription factor binding motifs. a**, DNA Patterns identified by TF-MoDISco that positively contribute to plantGREP predictions of enhancer strength in tobacco leaves at elevated ambient temperature. **b**, Transcription factor binding motifs that match the patterns in (**a**).

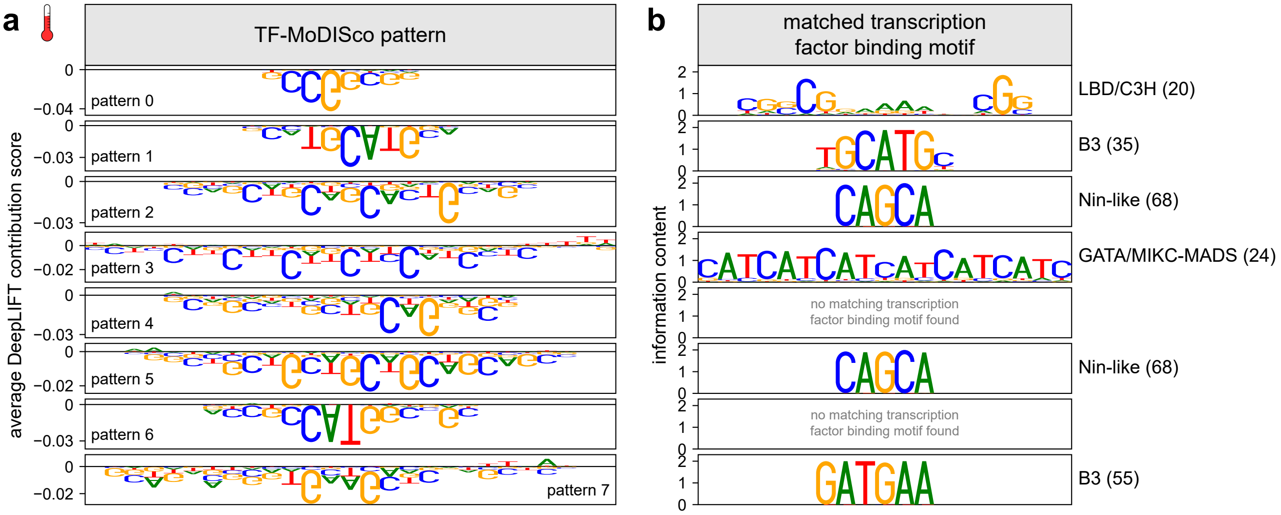


**Supplementary Fig. 17 | TF-MoDISco patterns match known transcription factor binding motifs. a**, DNA Patterns identified by TF-MoDISco that negatively contribute to plantGREP predictions of enhancer strength in tobacco leaves at elevated ambient temperature. **b**, Transcription factor binding motifs that match the patterns in (**a**).

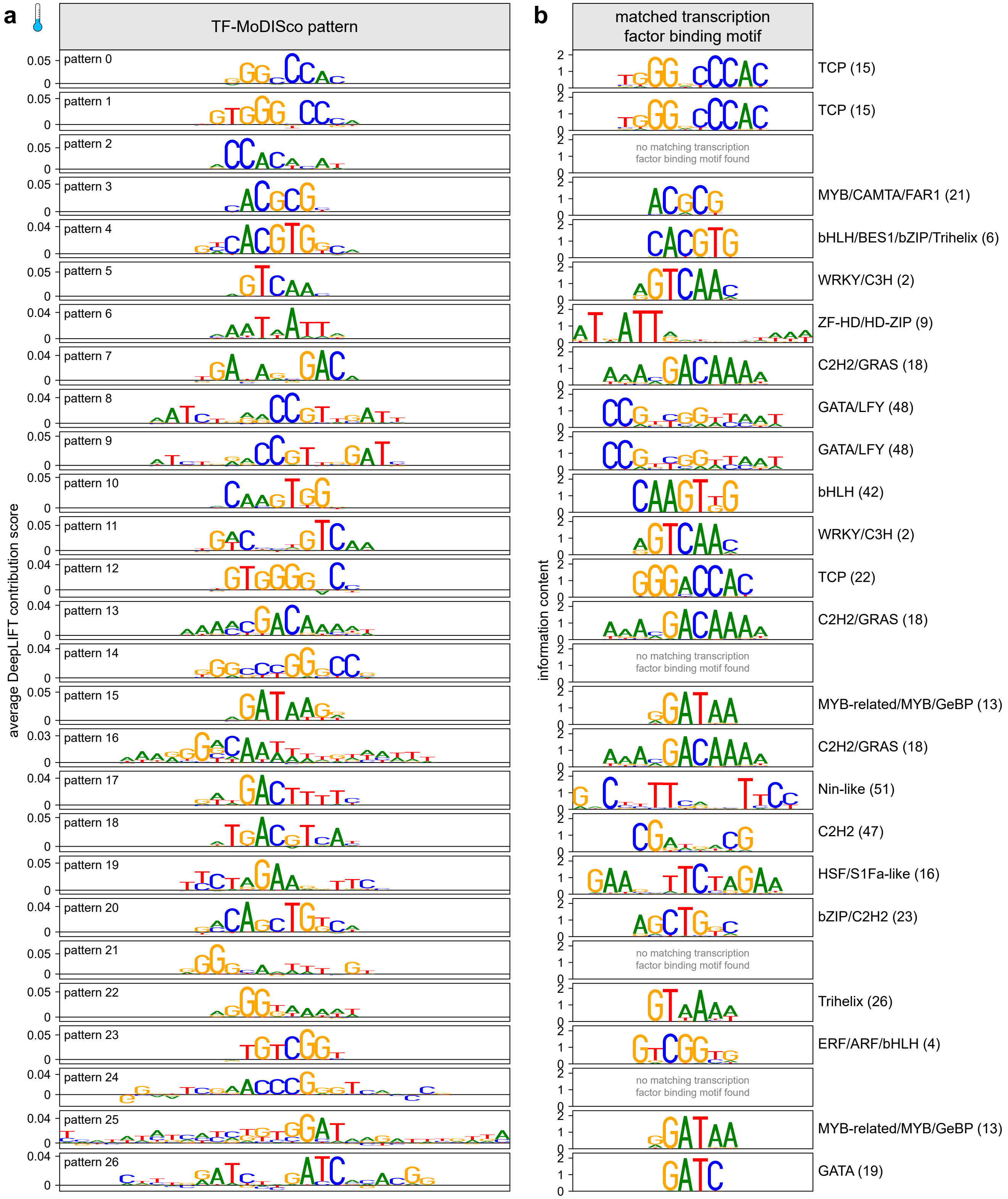


**Supplementary Fig. 18 | TF-MoDISco patterns match known transcription factor binding motifs. a**, DNA Patterns identified by TF-MoDISco that positively contribute to plantGREP predictions of enhancer strength in tobacco leaves at reduced ambient temperature. **b**, Transcription factor binding motifs that match the patterns in (**a**).

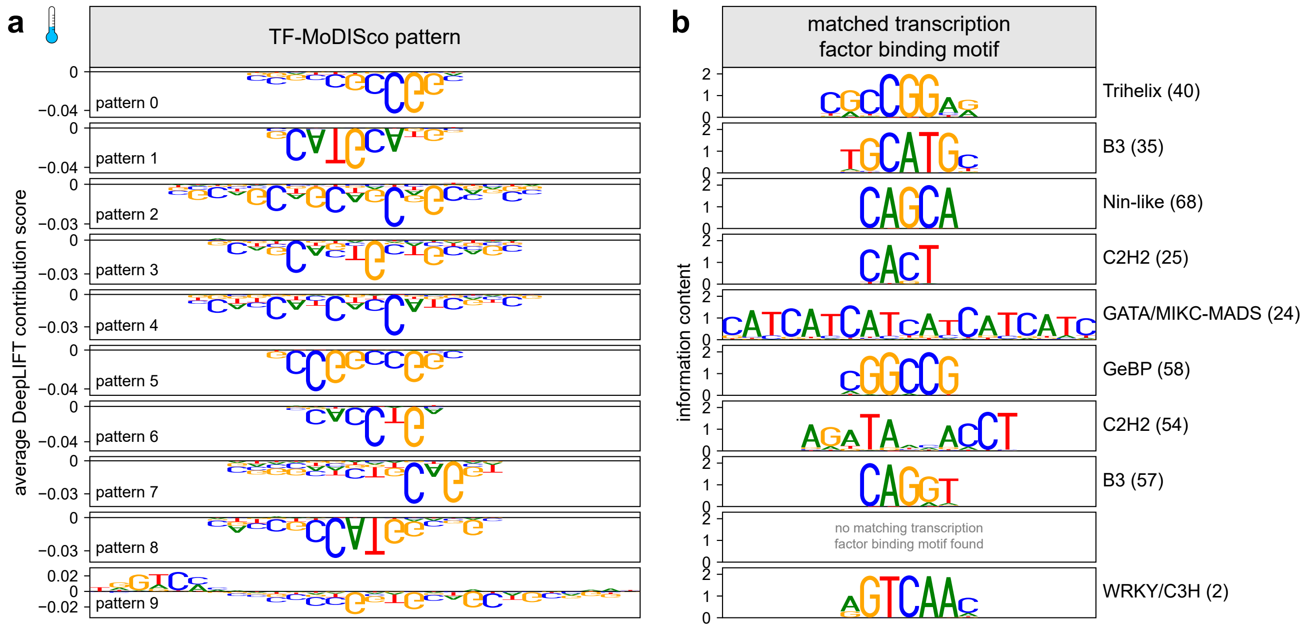


**Supplementary Fig. 19 | TF-MoDISco patterns match known transcription factor binding motifs. a**, DNA Patterns identified by TF-MoDISco that negatively contribute to plantGREP predictions of enhancer strength in tobacco leaves at reduced ambient temperature. **b**, Transcription factor binding motifs that match the patterns in (**a**).

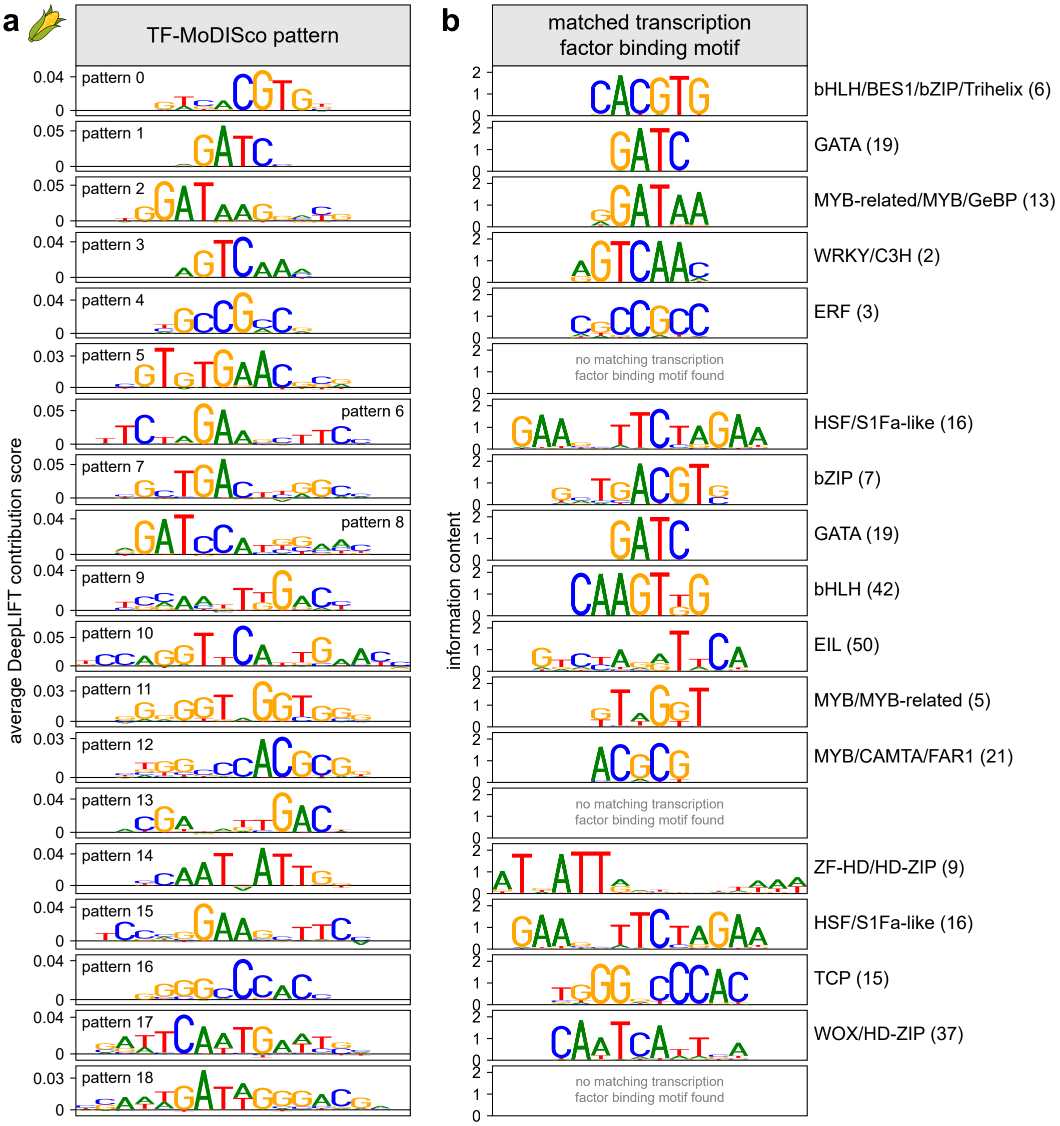


**Supplementary Fig. 20 | TF-MoDISco patterns match known transcription factor binding motifs. a**, DNA Patterns identified by TF-MoDISco that positively contribute to plantGREP predictions of enhancer strength in maize protoplasts. **b**, Transcription factor binding motifs that match the patterns in (**a**).

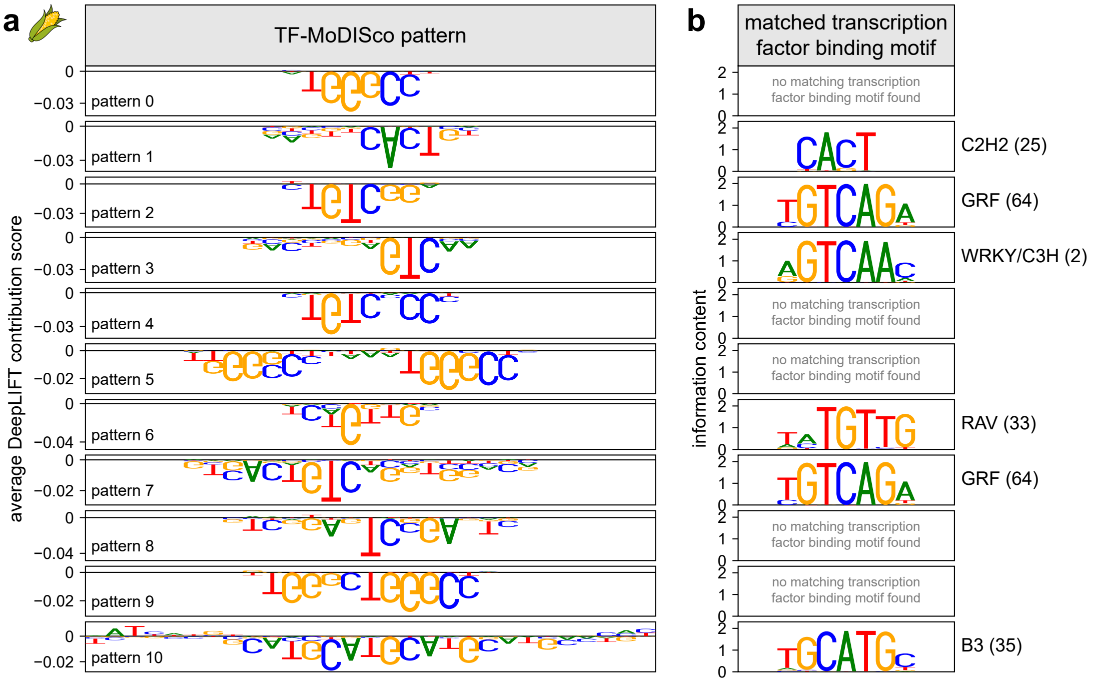


**Supplementary Fig. 21 | TF-MoDISco patterns match known transcription factor binding motifs. a**, DNA Patterns identified by TF-MoDISco that negatively contribute to plantGREP predictions of enhancer strength in maize protoplasts. **b**, Transcription factor binding motifs that match the patterns in (**a**).

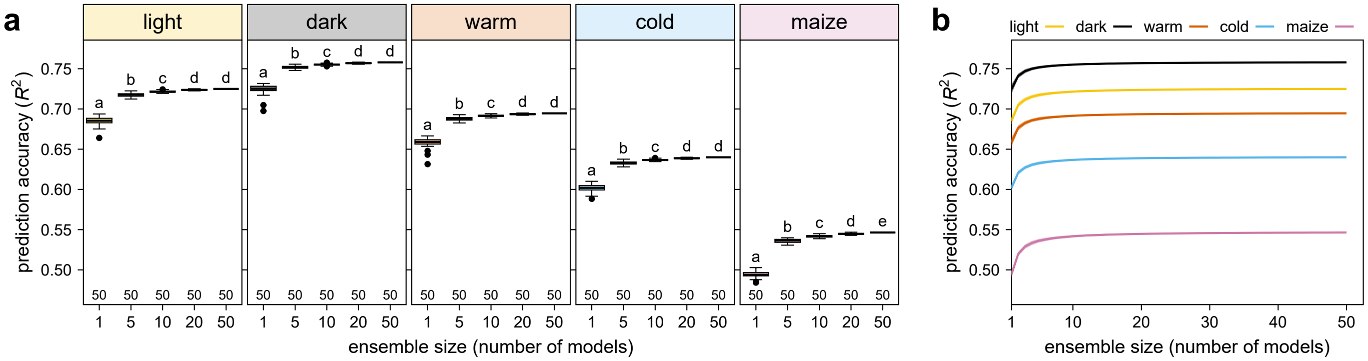


**Supplementary Fig. 22 | A model ensemble increases the prediction accuracy of plantGREP. a**,**b**, To build a plantGREP model ensemble, 50 models were trained with different, random initializations of the starting model weights. Box plots (as defined in Supplementary Fig. 9) in (**a**) show the prediction accuracy of a plantGREP model ensemble consisting of 1, 5, 10, 20, or 50 randomly sampled models. Lines in (**b**) indicate the mean prediction accuracy and the shaded area indicates the mean ± standard deviation. For each ensemble size, models were sampled independently for a total of 50 times. Letters above the plots in (**a**) indicate significance groups determined by post-hoc Tukey tests performed separately for each condition and species. Exact $p$ values are listed in Supplementary Data 4.

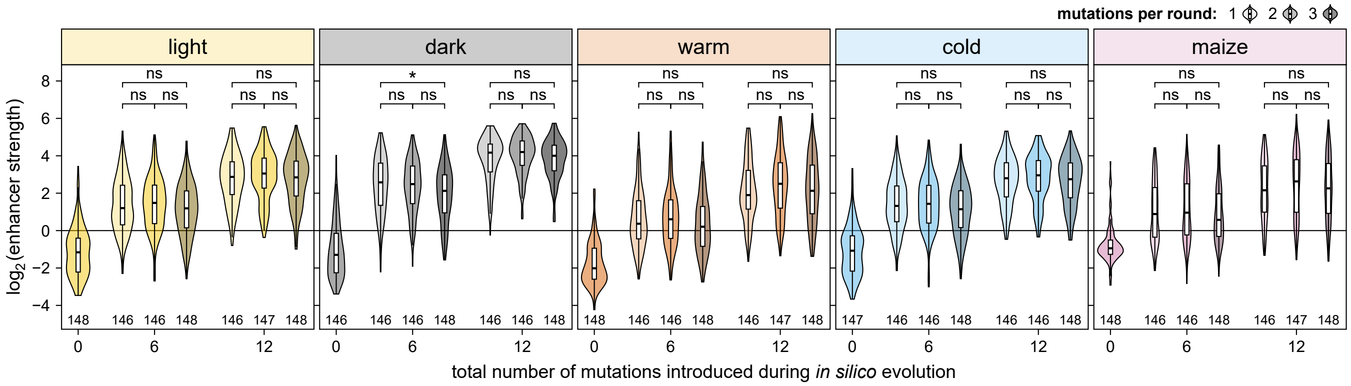


**Supplementary Fig. 23 | Increasing the number of mutations per round during *in silico* evolution did not increase performance.** *In silico* evolution was performed as in Fig. 4a (*i.e.*, with one mutation per round) or by inserting two or three mutations within an 8-bp window in each round. Sequences before (0 mutations) or after inserting six or twelve mutations during *in silico* evolution were subjected to Plant STARR-seq in the indicated conditions and species. Violin plots are as defined in Supplementary Fig. 4. Significant differences between sequences generated with one, two, or three mutations per round were determined by two-sided Wilcoxon rank-sum tests. The obtained *p* values were Bonferroni-adjusted separately for each condition and species and are indicated: *, $p \leq 0.05$; **, $p \leq 0.01$; ***, $p \leq 0.001$; ns, not significant. Exact *p* values are listed in Supplementary Data 4.

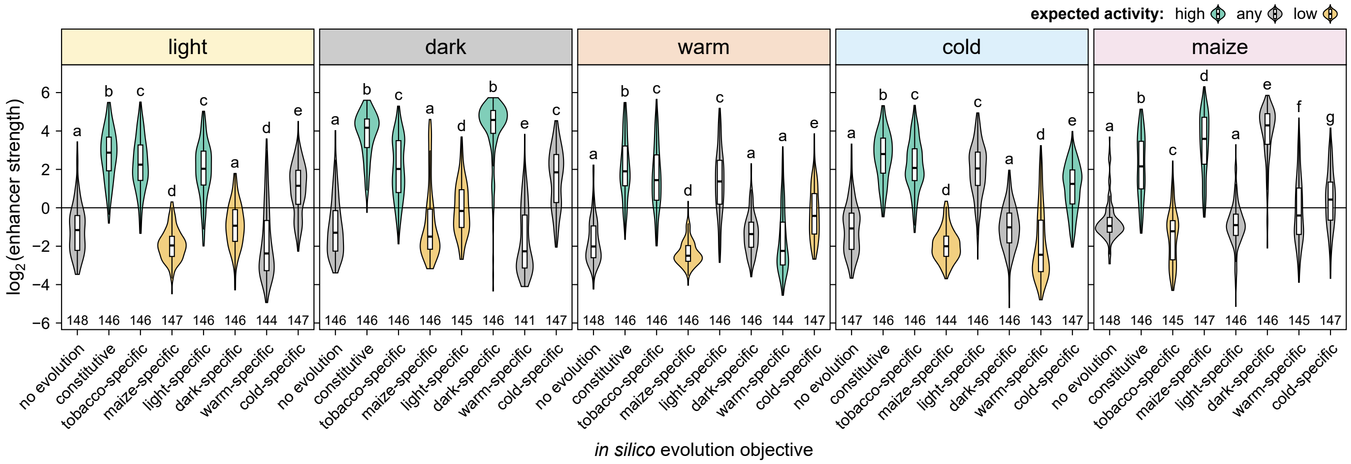


**Supplementary Fig. 24 | *In silico* evolved enhancers show species- and condition-specific activity.** Violin plots (as defined in Supplementary Fig. 4 and colored according to the expected activity in the indicated condition) of the strength of enhancers before (no evolution) or after twelve rounds of *in silico* evolution using plantGREP with different objectives to generate constitutive, species-, or condition-specific enhancers. Letters above the plots indicate significance groups determined by post-hoc Tukey tests performed separately for each condition and species. Exact *p* values are listed in Supplementary Data 4.
