## Supplementary Tables for "Massively parallel characterization and deep learning of enhancers in plant genomes"

### Supplementary Table 1 | Composition of the ACR library.

| category | *Arabidopsis* | tomato | maize | sorghum | total |
|---|---|---|---|---|---|
| ACR sequences | 58,068 | 24,166 | 52,282 | 35,314 | 169,830 |
| non-ACR sequences | 1,673 | 2,164 | 1,561 | 1,066 | 6,464 |
| genes of interest | | 15,670 | | | 15,670 |
| known regulatory elements | | | | | 36 |
| | | | | | **192,000** |

### Supplementary Table 2 | Tomato genes of interest with sequences tiled across their upstream regions.

| gene ID | description | # sequences |
|---|---|---|
| Solyc01g009560 | Terminal flower 1 | 79 |
| Solyc01g009580 | Terminal flower 1 | 76 |
| Solyc01g014100 | Unknown Protein | 177 |
| Solyc01g066950 | Flowering promoting factor-like 1 | 177 |
| Solyc01g066970 | Flowering promoting factor-like 1 | 83 |
| Solyc01g066980 | Flowering promoting factor-like 1 | 177 |
| Solyc01g080770 | Receptor like kinase, RLK | 177 |
| Solyc01g087990 | MADS-box transcription factor 3 | 68 |
| Solyc01g092950 | MADS-box transcription factor 2 | 92 |
| Solyc01g093960 | MADS box transcription factor | 87 |
| Solyc01g098390 | GID1-like gibberellin receptor | 82 |
| Solyc01g098890 | Unknown Protein | 28 |
| Solyc01g103530 | Receptor like kinase, RLK | 177 |
| Solyc02g032180 | Nuclear transcription factor Y subunit B-9 | 177 |
| Solyc02g065290 | Dof zinc finger protein 6 | 177 |
| Solyc02g065730 | MADS box transcription factor | 59 |
| Solyc02g067550 | Unknown Protein | 21 |
| Solyc02g069510 | Light-dependent short hypocotyls 1 | 71 |
| Solyc02g071730 | MADS-box transcription factor AGAMOUS | 114 |
| Solyc02g076820 | Light-dependent short hypocotyls 1 | 177 |
| Solyc02g077390 | WUSCHEL-related homeobox 11 | 137 |
| Solyc02g079020 | AP2 domain-containing transcription factor | 46 |
| Solyc02g079290 | SELF PRUNING 2G | 168 |
| Solyc02g081670 | Fimbriata | 78 |
| Solyc02g082670 | WUSCHEL-related homeobox 14 | 80 |
| Solyc02g083520 | BZIP transcription factor | 83 |
| Solyc02g083950 | WUSCHEL-related homeobox-containing protein 4 | 154 |
| Solyc02g084630 | MADS-box transcription factor | 177 |
| Solyc02g087470 | Unknown Protein | 14 |
| Solyc02g089200 | MADS-box transcription factor | 80 |
| Solyc02g089210 | MADS box transcription factor | 114 |
| Solyc02g090220 | Dof zinc finger protein | 88 |
| Solyc02g091840 | Receptor like kinase, RLK | 76 |
| Solyc02g092050 | AP2-like ethylene-responsive transcription factor At1g16060 | 24 |
| Solyc02g094460 | AP2 domain-containing transcription factor | 22 |
| Solyc03g006830 | MADS-box transcription factor 2 | 67 |
| Solyc03g007050 | Receptor like kinase, RLK | 177 |
| Solyc03g019710 | MADS-box transcription factor | 90 |
| Solyc03g025960 | Unknown Protein | 14 |
| Solyc03g026050 | Terminal flower 1 | 142 |
| Solyc03g043770 | Receptor like kinase, RLK | 177 |
| Solyc03g063100 | SELF PRUNING 3D | 177 |
| Solyc03g096300 | WUSCHEL-related homeobox 5 | 177 |
| Solyc03g114830 | MADS box transcription factor | 117 |
| Solyc03g114840 | MADS box transcription factor 11 | 177 |
| Solyc03g115850 | NAC domain protein IPR003441 | 117 |
| Solyc03g117230 | Ethylene responsive transcription factor 12 | 82 |
| Solyc03g118160 | FLORICAULA/LEAFY-like protein | 54 |
| Solyc03g118770 | WUSCHEL-related homeobox-containing protein 4 | 133 |
| Solyc03g119100 | Terminal flower 1 | 30 |
| Solyc03g123430 | AP2-like ethylene-responsive transcription factor At1g16060 | 26 |
| Solyc04g005320 | MADS-box transcription factor | 105 |
| Solyc04g009980 | Light-dependent short hypocotyls 1 | 61 |
| Solyc04g015060 | Nuclear transcription factor Y subunit B-6 | 36 |
| Solyc04g040220 | NPR1-like protein | 177 |
| Solyc04g056640 | LRR receptor-like serine/threonine-protein kinase, RLP | 158 |
| Solyc04g072570 | Receptor like kinase, RLK | 92 |
| Solyc04g076280 | MPF2-like | 71 |
| Solyc04g076680 | MADS-box transcription factor-like protein | 177 |
| Solyc04g077490 | AP2-like ethylene-responsive transcription factor At1g16060 | 127 |
| Solyc04g078390 | F-box protein | 177 |

| gene ID | description | # sequences |
|---|---|---|
| Solyc04g078650 | WUSCHEL-related homeobox-containing protein 4 | 154 |
| Solyc04g080080 | Glycosyltransferase family 77 protein | 29 |
| Solyc04g081000 | MADS box transcription factor | 66 |
| Solyc04g081590 | Receptor like kinase, RLK | 107 |
| Solyc05g006610 | Unknown Protein | 177 |
| Solyc05g007650 | Unknown Protein | 55 |
| Solyc05g056620 | Macrocalyx | 33 |
| Solyc05g012020 | Ripening Inhibitor | 177 |
| Solyc05g015720 | MADS box transcription factor-like protein | 148 |
| Solyc05g015730 | MADS-box transcription factor 1 | 120 |
| Solyc05g015750 | Transcription factor MADS-box 2 | 177 |
| Solyc05g023760 | Receptor-like protein kinase 5 | 53 |
| Solyc05g053630 | Unknown Protein | 79 |
| Solyc05g053850 | SELF PRUNING 5G | 117 |
| Solyc05g055020 | Light-dependent short hypocotyls 1 | 55 |
| Solyc05g055660 | SELF PRUNING 6A | 68 |
| Solyc06g008870 | GID1-like gibberellin receptor | 95 |
| Solyc06g035570 | Transcription factor MADS-box | 177 |
| Solyc06g059970 | MADS box transcription factor | 118 |
| Solyc06g064840 | Agamous MADS-box transcription factor | 168 |
| Solyc06g069430 | MADS box transcription factor | 95 |
| Solyc06g069710 | NAC domain protein IPR003441 | 86 |
| Solyc06g072890 | WUSCHEL-related homeobox 11 | 39 |
| Solyc06g074060 | Unknown Protein | 49 |
| Solyc06g074350 | self-pruning | 77 |
| Solyc06g076000 | WUSCHEL-related homeobox-containing protein 4 | 62 |
| Solyc06g082210 | Light-dependent short hypocotyls 1 | 115 |
| Solyc06g082520 | AP2 domain-containing transcription factor | 70 |
| Solyc06g083590 | B3 domain-containing transcription factor ABI3 | 53 |
| Solyc06g083600 | B3 domain-containing transcription factor ABI3 | 53 |
| Solyc06g083860 | Light-dependent short hypocotyls 1 | 64 |
| Solyc07g021170 | Os04g0226400 protein | 177 |
| Solyc07g053370 | Unknown Protein | 93 |
| Solyc07g055920 | Agamous MADS-box transcription factor | 177 |
| Solyc07g062470 | Light-dependent short hypocotyls 1 | 89 |
| Solyc07g062670 | Unknown Protein | 177 |
| Solyc07g062840 | NAC domain protein IPR003441 | 108 |
| Solyc08g041770 | Os04g0226400 protein | 177 |
| Solyc08g061560 | Receptor like kinase, RLK | 177 |
| Solyc08g067230 | MADS box transcription factor | 26 |
| Solyc08g075340 | Glycosyltransferase-like protein | 68 |
| Solyc08g078800 | GRAS family transcription factor | 5 |
| Solyc08g080100 | MADS box transcription factor | 141 |
| Solyc09g007260 | AP2-like ethylene-responsive transcription factor At1g79700 | 71 |
| Solyc09g009560 | SELF PRUNING 9D | 66 |
| Solyc09g025280 | Light-dependent short hypocotyls 1 | 54 |
| Solyc09g061410 | Unknown Protein | 177 |
| Solyc09g074270 | Acetyl esterase | 177 |
| Solyc09g090180 | Light-dependent short hypocotyls 1 | 177 |
| Solyc09g091810 | Unknown Protein | 90 |
| Solyc10g007310 | Light-dependent short hypocotyls 1 | 136 |
| Solyc10g008000 | Light-dependent short hypocotyls 1 | 177 |
| Solyc10g017630 | MADS-box transcription factor | 43 |
| Solyc10g017640 | MADS-box transcription factor | 177 |
| Solyc10g075030 | AP2 domain-containing transcription factor | 122 |
| Solyc10g079460 | NPR1-like protein | 177 |
| Solyc10g079750 | NPR1-like protein | 177 |
| Solyc11g005120 | MADS-box transcription factor 29 | 50 |
| Solyc11g008640 | Flowering locus T | 115 |
| Solyc11g008650 | Flowering locus T1 | 27 |
| Solyc11g008660 | Flowering locus T | 114 |
| Solyc11g010570 | MADS box transcription factor | 130 |
| Solyc11g010710 | AP2-like ethylene-responsive transcription factor At1g16060 | 177 |

| gene ID | description | # sequences |
|---|---|---|
| Solyc11g011260 | GAI-like protein 1 | 177 |
| Solyc11g028020 | Agamous MADS-box transcription factor | 177 |
| Solyc11g032100 | MADS box transcription factor | 177 |
| Solyc11g064850 | Os04g0226400 protein | 149 |
| Solyc11g066120 | Unknown Protein | 77 |
| Solyc12g038510 | MADS box transcription factor 11 | 177 |
| Solyc11g071380 | Unknown Protein | 65 |
| Solyc11g072770 | WUSCHEL-related homeobox 3B | 110 |
| Solyc11g072790 | WUSCHEL-related homeobox 3B | 106 |
| Solyc12g006340 | Auxin response factor 6 | 146 |
| Solyc12g014260 | Light-dependent short hypocotyls 1 | 69 |
| Solyc12g044760 | Os04g0226400 protein | 45 |
| Solyc12g056460 | MADS box transcription factor | 127 |
| Solyc12g087810 | Flowering locus C-like MADS-box protein | 94 |
| Solyc12g087820 | MADS-box transcription factor 13 | 42 |
| Solyc12g087830 | MADS box transcription factor | 42 |
| Solyc12g088090 | MADS-box transcription factor-like protein | 163 |
| Solyc12g099610 | Flowering promoting factor-like 1 | 34 |

### Supplementary Table 3 | Regulatory activity of test sequences.

| category[a] | light | dark | warm | cold | maize | at least one condition or assay system | all conditions and assay systems |
|---:|---:|---:|---:|---:|---:|---:|---:|
| enhancing sequences | 13.6% | 19.3% | 22.1% | 11.1% | 4.7% | 33.0% | 0.9% |
| inactive sequences | 75.5% | 69.2% | 76.1% | 77.9% | 84.3% | 98.7% | 42.9% |
| repressive sequences | 10.9% | 11.5% | 1.8% | 11.0% | 11% | 26.7% | 0.3% |

[a] Categories: enhancing sequences, $\log_2$(enhancer strength) > 1; inactive sequences, $\log_2$(enhancer strength) between −2 and 1; repressive sequences, $\log_2$(enhancer strength) < −2

**Supplementary Table 4 | Consensus binding motifs for transcription factors of the indicated families.** Consensus motifs were obtained from ref. [39]

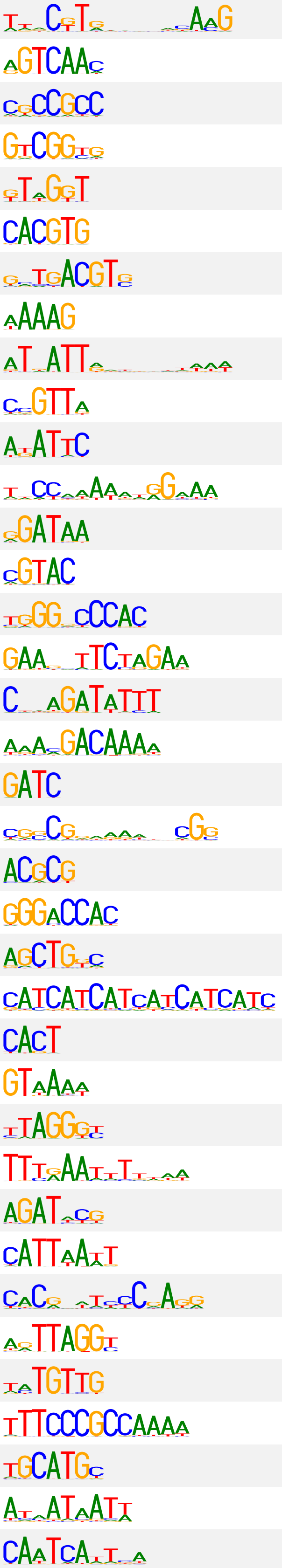

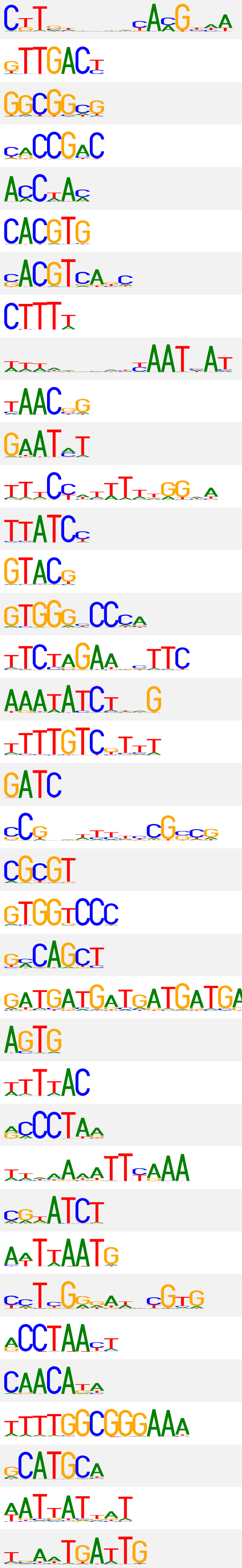

| motif ID | motif logo (forward) | motif logo (reverse-complement) | transcription factor families |
|---|---|---|---|
| 1 | | | NAC |
| 2 | | | WRKY, C3H |
| 3 | | | ERF |
| 4 | | | ERF, ARF, bHLH |
| 5 | | | MYB, MYB-related |
| 6 | | | bHLH, BES1, bZIP, Trihelix |
| 7 | | | bZIP |
| 8 | | | Dof, C3H |
| 9 | | | ZF-HD, HD-ZIP |
| 10 | | | MYB, Trihelix |
| 11 | | | G2-like |
| 12 | | | MIKC-MADS |
| 13 | | | MYB-related, MYB, GeBP |
| 14 | | | SBP, AP2, C2H2, bHLH |
| 15 | | | TCP |
| 16 | | | HSF, S1Fa-like |
| 17 | | | MYB-related, C2H2, AP2 |
| 18 | | | C2H2, GRAS |
| 19 | | | GATA |
| 20 | | | LBD, C3H |
| 21 | | | MYB, CAMTA, FAR1 |
| 22 | | | TCP |
| 23 | | | bZIP, C2H2 |
| 24 | | | GATA, MIKC-MADS |
| 25 | | | C2H2 |
| 26 | | | Trihelix |
| 27 | | | MYB-related, Trihelix |
| 28 | | | CPP |
| 29 | | | ARR-B |
| 30 | | | HD-ZIP |
| 31 | | | AP2 |
| 32 | | | MYB |
| 33 | | | RAV |
| 34 | | | E2F, DP |
| 35 | | | B3 |
| 36 | | | HD-ZIP, YABBY |
| 37 | | | WOX, HD-ZIP |

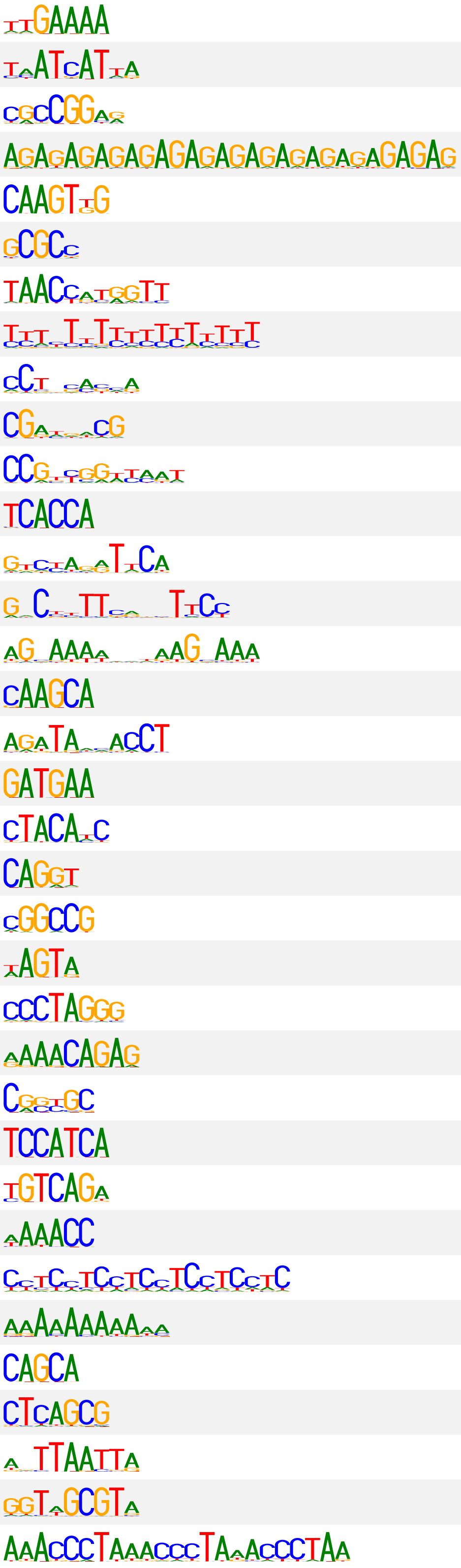

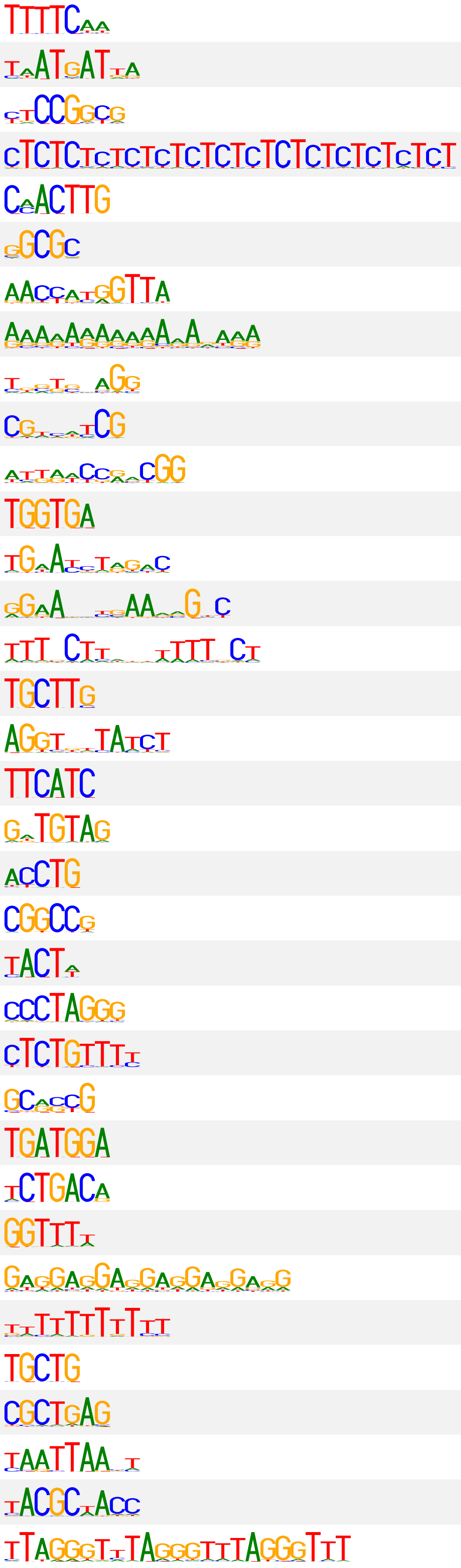

| motif ID | motif logo (forward) | motif logo (reverse-complement) | transcription factor families |
|---|---|---|---|
| 38 | | | NF-YB, C2H2, G2-like |
| 39 | | | HD-ZIP, YABBY |
| 40 | | | Trihelix |
| 41 | | | BBR-BPC |
| 42 | | | bHLH |
| 43 | | | E2F, DP |
| 44 | | | Trihelix |
| 45 | | | MIKC-MADS, AP2 |
| 46 | | | C2H2 |
| 47 | | | C2H2 |
| 48 | | | GATA, LFY |
| 49 | | | M-type-MADS |
| 50 | | | EIL |
| 51 | | | Nin-like |
| 52 | | | B3 |
| 53 | | | B3 |
| 54 | | | C2H2 |
| 55 | | | B3 |
| 56 | | | B3 |
| 57 | | | B3 |
| 58 | | | GeBP |
| 59 | | | C2H2 |
| 60 | | | SRS |
| 61 | | | C2H2 |
| 62 | | | ERF |
| 63 | | | NF-YB |
| 64 | | | GRF |
| 65 | | | Trihelix |
| 66 | | | C2H2 |
| 67 | | | B3 |
| 68 | | | Nin-like |
| 69 | | | MYB |
| 70 | | | WOX |
| 71 | | | VOZ |
| 72 | | | SRS |

**Supplementary Table 5 | Oligonucleotides used in this study.**

| sequence | purpose |
|---|---|
| GTTCCACTTCTGTATAGGAGCCAATTG | linearize pPSup to replace terminator (fwd primer) |
| TTACTTGTACAGCTCGTCCATGC | linearize pPSup to replace terminator (rev primer) |
| TGGACGAGCTGTACAAGTAAGCTCTAGCTAGAGTCGATCGACAAGCTCGAGTTTCTCCATAAT<br>AATGTGTGAGTAGTTCCCAGATAAGGGAATTAGGGTTCCTATAGGGTTTCGCTCATGTGTTGA<br>GCATATAAGAAACCCTTAGTATGTATTTGTATTTGTAAAATACTTCTATCAATAAAATTTCTA<br>ATTCCTAAAACCAAAATCCAGTACTAAAATCCAGATAGGTGTTCCACTTCTGTATAGGAGC | 35S terminator for pPSnt |
| AATTCCACTCCGAGACCACAAGGCGCGCCTAGTGGTCTCCCTGTGCAACCGTCTTCAGTGCCT<br>GCA | replace BsaI cassette for pPSntF (fwd primer) |
| GGCACTGAAGACGGTTGCACAGGGAGACCACTAGGCGCGCCTTGTGGTCTCGGAGTGG | replace BsaI cassette for pPSntF (rev primer) |
| AATTCCACAGCGAGACCACAAGGCGCGCCTAGTGGTCTCCGAGTGCAACCGTCTTCAGTGCCT<br>GCA | replace BsaI cassette for pPSntR (fwd primer) |
| GGCACTGAAGACGGTTGCACTCGGAGACCACTAGGCGCGCCTTGTGGTCTCGCTGTGG | replace BsaI cassette for pPSntR (rev primer) |
| GCAAGACCCTTCCTCTATATAAGGAAGTTCATTTCATTTGGAGAGGACACGCCCGTCCGAACT<br>CCGAACCCCAGAACAGAGCAAAGCCTCCTCGGCCTCCCTGTCCCCAGCCTTCCCCGATG | template 35S minimal promoter + Zm00001d041672 5′ UTR |
| GAGAGGAAGACCC-GCAAGACCCTTCCTCTATATAAGGAAGTTCATTTC | amplify 35S minimal promoter + 5′ UTR (fwd primer) |
| GAGAGGAAGACGGTCAC-NNBNNCNNBNNTNNBNNB-CATCGGGGAAGGCTGGG | amplify 35S minimal promoter + Zm00001d041672 5′ UTR (rev primer for pPSntF) |
| GAGAGGAAGACGGTCAC-NNBNNTNNBNNCNNBNNB-CATCGGGGAAGGCTGGG | amplify 35S minimal promoter + Zm00001d041672 5′ UTR (rev primer for pPSntR) |
| GAGAGGAAGACGGTCAC-NNBNNGNNBNNGNNBNNB-CATCGGGGAAGGCTGGG | amplify 35S minimal promoter + Zm00001d041672 5′ UTR (rev primer for no-enhancer control) |
| CCTTCCTCTATATAAGGAAGTTCATTTCATTTGGAGAGGACACG | linearize pDL to replace BsaI cassette (fwd primer) |
| GGAATTCGATATCAAGCTTATCGATACCG | linearize pDL to replace BsaI cassette (rev primer) |
| CGGTATCGATAAGCTTGATATCGAATTCC | amplify BsaI cassette + 35S minimal promoter from pPSnt (fwd primer) |
| CTCCAAATGAAATGAACTTCCTTATATAGAGGAAG | amplify BsaI cassette + 35S minimal promoter from pPSnt (rev primer) |
| GAGCGGTCTCCACTC<170-bp enhancer candidate>CTGTAGAGACCGGGC | oligo pool enhancer candidates |
| GAGCGGTCTCCACTC | amplify oligo pool sequences (fwd primer) |
| GCCCGGTCTCTACAG | amplify oligo pool sequences (rev primer) |
| CAATCCGCCCTCACTACAACCGACAGGAAACAGCTATGACCATGATTACGCCAAGCTTGCATG<br>CCTGCAGGTCGACGGTCTCCACTCATTCTAAAGGTCAAAGCTTGATCCGAGCAGAGTATTACT<br>GATTGTATATAATATTCCTAAGTAAATTGAAACTGACATGCATTCAAAGTCAACCTTGCTGCT<br>ACTGCTTATCCGAAGCATCATGGTACTCATCCTTATCTCCCTCTTCTTCAGCCACCTCAGGCA<br>CTGTACTGTCGAGACCTCTAGAGGATCCCCGGGTACCGAGCTCGAATTCACTGGCCGTCGTTT<br>TACAACGCTACTCTGGCGTCGATGAGGGA | gene fragment At-9661 |
| CAATCCGCCCTCACTACAACCGACAGGAAACAGCTATGACCATGATTACGCCAAGCTTGCATG<br>CCTGCAGGTCGACGGTCTCCACTCATAGAAAAATTAGGTAAAGAGTCAGTGTCGTTATGTTAT<br>GGAAGATGTGAATGAAGTTTGACTTCTCATTGTATATGAGTAAAATCTTTTCTTACAAGGGAA<br>GTCCCCAATTGGTCAACATGTGAAAGCACGTGTCATGTTCTTACTTTTGTTTGGGTAATCTTC<br>TAATTCTGTCGAGACCTCTAGAGGATCCCCGGGTACCGAGCTCGAATTCACTGGCCGTCGTTT<br>TACAACGCTACTCTGGCGTCGATGAGGGA | gene fragment Sl-12881 |
| CAATCCGCCCTCACTACAACCGACAGGAAACAGCTATGACCATGATTACGCCAAGCTTGCATG<br>CCTGCAGGTCGACGGTCTCCACTCAATTTAAGAGTAAATGAGAGAAGTTGAATTAGGGCCTTT<br>CGTGGACCAAATTCGTTGCTGCTTTTGGGGTTTAACCCAATTCCTCTTTTTGTATATATCTGT<br>TGTATATCAGTGTATATCGGATGTATACAACCCATTTTTTGTTTTTCCAGACCTTTTGGGGAG<br>CAGAGCTGTCGAGACCTCTAGAGGATCCCCGGGTACCGAGCTCGAATTCACTGGCCGTCGTTT<br>TACAACGCTACTCTGGCGTCGATGAGGGA | gene fragment At-11716 |
| CAATCCGCCCTCACTACAACCGACAGGAAACAGCTATGACCATGATTACGCCAAGCTTGCATG<br>CCTGCAGGTCGACGGTCTCCACTCACTCTTTAGCAACCATCGTCGTCATGCGGCCCATCAACA<br>CACGTGATATCTTATCTTCTCGGGAATTGAGTGCACGCGAGGCGGCCCATTCCCAACAGCTTT<br>TGTTTCGCTGGAGTATCTTGGCGTGCCAGACTGCCAGTGCGTCATGTGATGACCGGTGCAGTG<br>CCCTACTGTCGAGACCTCTAGAGGATCCCCGGGTACCGAGCTCGAATTCACTGGCCGTCGTTT<br>TACAACGCTACTCTGGCGTCGATGAGGGA | gene fragment At-12871 |
| CAATCCGCCCTCACTACAACCGACAGGAAACAGCTATGACCATGATTACGCCAAGCTTGCATG<br>CCTGCAGGTCGACGGTCTCCACTCCATGGCATCATGCTATCATGTGTGGTTGTCGAAGTCTCC<br>ATCCGTTGCGCTTGCGGTACGCCGTTGACCCGGGCAACGCGTGTTCCTTGCCACGCAGGCGAT<br>GATGGATCGGACGGTGCAGACAGCCTCGAGATCCGTGGATCCAGCGGGTAACTAGCCTTCCCC<br>TGTCCCTGTCGAGACCTCTAGAGGATCCCCGGGTACCGAGCTCGAATTCACTGGCCGTCGTTT<br>TACAACGCTACTCTGGCGTCGATGAGGGA | gene fragment At-13524 |
| CAATCCGCCCTCACTACAACCGACAGGAAACAGCTATGACCATGATTACGCCAAGCTTGCATG<br>CCTGCAGGTCGACGGTCTCCACTCATCCAATAGTCATCTTTGTAAAGATTTTTTGTTCGTGTG<br>TGGCGGATAAGCAAAAACAGAATAACATAATCTCTTATCCAACTTATATTGCCACGTGGACAG<br>ATATAGTTGGTGAAGCAGATACTAATCTAATCTGAGCAAAACCAGTTGGGCTATTTGAAAGCC<br>ACAAGCTGTCGAGACCTCTAGAGGATCCCCGGGTACCGAGCTCGAATTCACTGGCCGTCGTTT<br>TACAACGCTACTCTGGCGTCGATGAGGGA | gene fragment At-2020 |

| sequence | purpose |
|---|---|
| CAATCCGCCCTCACTACAACCGACAGGAAACAGCTATGACCATGATTACGCCAAGCTTGCATG<br>CCTGCAGGTCGACGGTCTCCACTCAATAGACATGGACATTAGGATAGAGAAATTGGATCAAAT<br>TTTCAACTAATTCTCATCCTCAATAGAAGTGGGACCCTTTGAAATCCATTTGACTACAACTTT<br>ATCCAATCAAAAACAAGTAATAAAGATGATGTTTATAATTTTAGCCAATTAAAATACAGATAA<br>GGCCTCTGTCGAGACCTCTAGAGGATCCCCGGGTACCGAGCTCGAATTCACTGGCCGTCGTTT<br>TACAACGCTACTCTGGCGTCGATGAGGGA | gene fragment At-1 |
| CAATCCGCCCTCACTACAACCGACAGGAAACAGCTATGACCATGATTACGCCAAGCTTGCATG<br>CCTGCAGGTCGACGGTCTCCACTCTAAGGTTCCATAAGATCCACAAGAAAAGGATAAAACGGT<br>AAGGAGTGTGCAATGTGCACCACTTAAAATCTATTGCTATTCCAGGCTTAGGCCCATTCTCGG<br>ACCCAATTTCAGTTTTGTTTGGAAGTGTTTCAACTTTTTCTTTTTTTATGTTAACCGATGTGA<br>TTCAACTGTCGAGACCTCTAGAGGATCCCCGGGTACCGAGCTCGAATTCACTGGCCGTCGTTT<br>TACAACGCTACTCTGGCGTCGATGAGGGA | gene fragment Sl-774 |
| CAATCCGCCCTCACTACAACCGACAGGAAACAGCTATGACCATGATTACGCCAAGCTTGCATG<br>CCTGCAGGTCGACGGTCTCCACTCAACGCACCAACAAACAAAATAGTTTCAAAACAAAACTTG<br>AAAATGTGGGGTCCACGTAAGAATCTCCATGGAAGAACTTGAGCCAATCAGAGTGCAAAAACG<br>GAGATGACAGTATATTTCAACCAATAAGGAGGCAGAATCAGATCACCAGTGGGTTCCATTATA<br>AGTTTCTGTCGAGACCTCTAGAGGATCCCCGGGTACCGAGCTCGAATTCACTGGCCGTCGTTT<br>TACAACGCTACTCTGGCGTCGATGAGGGA | gene fragment Sb-11289 |
| CAATCCGCCCTCACTACAACCGACAGGAAACAGCTATGACCATGATTACGCCAAGCTTGCATG<br>CCTGCAGGTCGACGGTCTCCACTCCACACTGTTGCGTAGTTTCATGAGATTCTATCATACACG<br>TACTTTCTTATGATATATTATTAAATTTTGTTGAAACGATAAGATGAAATTGATCCTTGTAAG<br>TCATGAGTCACCAGTTTGATCGATCCGATGATTGAGTTTATAACATTATTTTTATGTGGATCA<br>GTCTACTGTCGAGACCTCTAGAGGATCCCCGGGTACCGAGCTCGAATTCACTGGCCGTCGTTT<br>TACAACGCTACTCTGGCGTCGATGAGGGA | gene fragment Zm-23177 |
| GAACTTGTGGCCGTTTACG | reverse transcription primer for barcode sequencing |
| AATGATACGGCGACCACCGAGATCTACAC<8-bpindex2>CCCTCGAGGTCGACGGTATC | amplify pPSnt constructs for subassembly (fwd primer) |
| AATGATACGGCGACCACCGAGATCTACAC<8-bpindex2>CCTCGGCCTCCCTGTCC | amplify pPSnt constructs or reporter cDNA for barcode sequencing (fwd primer) |
| CAAGCAGAAGACGGCATACGAGAT<8-bpindex1>CACCCCGGTGAACAGCTCC | amplify pPSnt constructs or reporter cDNA for subassembly or barcode sequencing (rev primer) |
| CTCGAGGTCGACGGTATCGATAAGCTTGATATCGAATTCCAC | NGS read 1 primer for subassembly |
| CTCCAAATGAAATGAACTTCCTTATATAGAGGAAGGGTCTTGCAC | NGS read 2 primer for subassembly |
| CCTCGGCCTCCCTGTCCCCAGCCTTCCCCGATG | NGS index 1 primer for subassembly and NGS read 1 primer for barcode sequencing |
| CACCCCGGTGAACAGCTCCTCGCCCTTGCTCAC | NGS index 2 primer for subassembly and NGS read 2 primer for barcode sequencing |
| GTGAGCAAGGGCGAGGAGCTGTTCACCGGGGTG | NGS index 1 primer for barcode sequencing |
| CATCGGGGAAGGCTGGGGACAGGGAGGCCGAGG | NGS index 2 primer for barcode sequencing |
